# Cross-cohort optimal transport maps macrophage plasticity and routes to inflammation and fibrosis in human atherosclerosis

**DOI:** 10.64898/2026.05.15.725379

**Authors:** Sergio Vazquez Montes de Oca, Ariadna Acedo Terrades, Jose F Carreño Martinez, Philipp Kirchner, Tiit Örd, Minna U Kaikkonen, Siting Wei, Stefan Freigang, Inti Zlobec, María Rodríguez Martínez

## Abstract

**Background:** Single-cell transcriptomics has revealed extensive macrophage heterogeneity in atherosclerotic plaques, but how disease-associated programs reshape cellular identity remains unclear. Connecting macrophage states to these molecular changes could help identify candidate routes through which macrophages contribute to inflammation and fibrosis.

**Methods:** We developed a computational method combining optimal transport, RNA velocity and comparative program analysis across 34,533 monocytes and macrophages from seven human cohorts. Optimal transport quantified all 15 pairwise relationships among six macrophage states, and target-association gradients supported source- and destination-program comparisons across 11 relationships. Expression contrasts were calculated within libraries and source subtypes. Robustness was assessed through resampling, cohort omission and held-out-gene analyses. Paired RNA and protein measurements provided complementary molecular evidence, and patient-label permutation tests evaluated exploratory clinical associations in 18 patients. We also applied the method to an independent human liver dataset.

**Results:** Scavenging and inflammatory macrophages formed the closest transcriptional pair in the full atlas and every cohort-omission analysis. Scavenging macrophages retained selected source features alongside inflammatory programming, whereas fibrotic association coincided with attenuation of the broader scavenging signature. This signature showed a lower change along fibrotic than inflammatory association in all 19 matched libraries. Protein measurements corroborated selected changes and revealed coexisting scavenging and activation features within individual cells. Symptomatic patients had a higher fraction of monocytes with strong inflammatory association than asymptomatic patients (median difference, 13.0 percentage points; unadjusted one-sided permutation *p* = 0.0017). Analysis of 8,536 liver mononuclear phagocytes further resolved contrasting source-marker responses along scar-associated macrophage association.

**Conclusions:** These findings connect macrophage heterogeneity to selective identity remodeling across disease-associated states. Integrating population relationships with RNA, protein and clinical measurements provides a strategy for prioritizing candidate pathogenic routes and their associated molecular programs for functional investigation.

## 1 Background

Atherosclerosis underlies most cardiovascular deaths worldwide and is increasingly recognized as a chronic inflammatory disease of the arterial wall [1–3]. Despite aggressive lipid-lowering therapy, cardiovascular events persist [4], highlighting the need to understand inflammatory programs that are not directly targeted by current treatments. Macrophages are central to this inflammatory process: they dominate the immune compartment of human atherosclerotic plaques and contribute throughout disease progression, from lipid uptake and foam-cell formation to plaque destabilization, rupture, and inflammation resolution [5–8]. Reflecting these diverse roles, single-cell RNA sequencing has revealed a macrophage landscape far more heterogeneous than anticipated by the classical M1/M2 paradigm [9, 10], with transcriptionally distinct populations, including resident-like, TREM2^hi^ lipid-associated, foamy, inflammatory, and interferon-stimulated macrophages, recurrently identified across cohorts [11–15].

Recent atlases have further resolved disease-associated differences in macrophage abundance and transcriptional programs [16], emphasizing the need to understand both the populations present in a lesion and the relationships among their molecular states.

Characterizing the relationships among these states also requires understanding how their gene programs differ. Because macrophage activation involves both shared and stimulusspecific transcriptional programs [10], a central question in plaques is whether the acquisition of inflammatory or fibrosis-associated programs accompanies the attenuation of existing source-associated features, or whether these features are retained or reinforced. These patterns may vary with both the starting state and the disease-associated program being acquired. We therefore ask whether distinct macrophage populations show similar identity changes in association with a common disease-associated state, and whether a single population shows different changes across alternative states.

Mapping these relationships requires a methodology that compares predefined cell states and connects their relationships to changes in gene programs. Pseudotime methods organize cells along inferred trajectories [17], whereas RNA velocity estimates local directions of transcriptional change [18]. CellRank and CellRank 2 combine cell-level transition models with fate-probability estimation and identification of lineage-associated genes [19, 20]. Our focus is complementary: to quantify relationships among predefined macrophage populations and compare how source-associated and destination-associated programs vary across these relationships.

Optimal transport (OT) addresses this challenge by identifying the minimum-cost mapping between two cell populations in transcriptomic space. This provides a principled, multivariate measure of how closely the populations are related without imposing a predefined trajectory topology or assuming a fixed direction of change [21–23]. Although OT has transformed the analysis of developmental trajectories [21], its application to chronic disease remains limited, particularly in systems characterized by reversible and cyclic state transitions. We therefore combine pairwise OT with RNA velocity, inferred using UniTVelo’s unified latent-time model [24], to characterize connectivity and estimate orientations between macrophage states. OT quantifies population-level transcriptomic relatedness, while RNA velocity provides complementary directional estimates. We assess variability in OT divergence through cell-level and library-level resampling and evaluate sensitivity to cohort composition through leave-one-cohort-out (LOCO) analyses. Velocity model settings and initialization sensitivity are described in the Methods and Supplementary Table 5.

We apply this framework to a unified atlas of 34,533 monocytes and macrophages assembled from seven publicly available scRNA-seq cohorts of human atherosclerotic plaques (including carotid and coronary specimens). Across six macrophage meta-clusters, we characterize all 15 pairwise relationships using OT. UniTVelo identifies significant directional asymmetry in 13 of these pairs in the primary analysis. We further investigate source and destination-associated gene programs across 11 comparisons organized into three biological axes: monocyte diversification, inflammatory programming, and fibrotic-associated remodeling.

To characterize the transcriptional changes associated with these relationships, we rank source cells using OT-derived target-association scores and examine how source and destinationassociated gene programs vary along the resulting gradients. Classical monocyte markers decline along Scavenging and Inflammatory associations but increase along Fibrotic association.

Scavenging macrophages show maintenance and reinforcement of selected source-associated features alongside inflammatory programming, whereas several of these features are attenuated along Fibrotic association. Direct comparison across 19 matched libraries reveals consistently lower changes in the broader scavenging signature along Fibrotic than inflammatory association. Together, these findings characterize macrophage plasticity through selective remodeling of existing programs and show that a common source population can exhibit distinct molecular patterns across disease-associated relationships.

To examine whether these transcriptional patterns are reflected at the protein level, we analyze paired RNA and surface-protein measurements from an existing human plaque CITE-seq dataset [25]. These measurements allow us to assess whether selected scavengingassociated proteins are retained or attenuated along inflammatory and fibrosis-associated target-association gradients and whether scavenging and activation proteins are coexpressed in individual macrophages along the inflammatory gradient. This directly addresses whether features of the source state persist in the same cells that display activation-associated features, providing protein-level corroboration of selected patterns identified in the transcriptomic analysis.

Exploratory integration with patient-level clinical metadata further identifies a higher fraction of monocytes with strong inflammatory-state association in patients with symptomatic versus asymptomatic carotid disease, illustrating how the framework can connect within-state transcriptional variation to clinical presentation. To examine macrophage plasticity beyond atherosclerosis, we extend the OT-based connectivity and gene-program analyses to a published single-cell atlas of healthy and cirrhotic human liver [26]. We compare tissue monocytes and Kupffer cells in relation to scar-associated macrophages, asking whether the retention and attenuation of source-associated features can also be resolved in a distinct tissue and disease context. Together, these analyses provide a framework for comparing macrophage plasticity across states and tissues while relating selected molecular patterns to clinical disease.

## 2 Methods

### 2.1 Data collection and preprocessing

Raw scRNA-seq data were obtained from seven publicly available human atherosclerosis datasets: Alsaigh [27], Bashore [25], Pan [28], Jaiswal [29], Pauli [30], Fernandez [12], and Wirka [31]. Sequencing runs belonging to the same library were processed together, avoiding separate representation of cells sequenced across multiple lanes. Reads were processed using Cell Ranger v10.1.0 with the GRCh38-2020-A reference. BAM files were generated for quantification of spliced, unspliced, and ambiguous transcripts using velocyto [32].

Ambient RNA correction was performed per library using CellBender [33], with the unfiltered Cell Ranger matrix as input. The expected number of cells was obtained from the corresponding Cell Ranger metrics; settings were a false-positive rate of 0.01, 20,000 included droplets, and 150 training epochs. All libraries retained in the final atlas had CellBendercorrected expression matrices. The corrected RNA counts were used for expression analysis, whereas the spliced, unspliced, and ambiguous layers were obtained separately from velocyto and matched by library, barcode, and gene identifier. CellBender correction was not applied to these splicing layers.

### 2.2 Quality control

Quality control was performed using Scanpy [34]. Initial filtering retained cells with at least 200 detected genes and less than 25% mitochondrial counts. SOLO [35] was then trained separately for each library through scvi-tools [36], using an scVI model followed by SOLO.from_scvi_model. Both soft scores and hard classifications were retained.

For final doublet filtering, cells were ranked by their SOLO score within each library. The expected doublet fraction was defined as

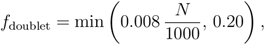

where *N* is the number of prefiltered cells in that library. The highest-scoring round(*Nf*_doublet_) cells were removed. Thus, filtering used an expected-rate ranking rather than a universal score threshold or direct removal of all SOLO hard calls.

Libraries containing fewer than 1,000 prefiltered cells were excluded. Following doublet filtering, cell-level quality control was applied separately within each retained library. Cells were excluded if log-transformed total counts or detected-gene counts fell more than five median absolute deviations below their respective medians, or if mitochondrial percentage exceeded five median absolute deviations above the library median. A hard mitochondrial threshold of less than 15% was additionally imposed. Libraries with fewer than 500 median detected genes after filtering were excluded. Per-library counts, exclusion flags, and executed settings are reported in Supplementary Table 9; retained-cell quality distributions are shown in Supplementary Figure 1.

### 2.3 Integration and clustering

Myeloid cells were identified by coarse lineage annotation and integrated across cohorts. Genes shared across the input datasets were retained. CellBender-corrected counts were normalized to 10,000 counts per cell and transformed using the natural logarithm of one plus normalized expression. The top 5,000 highly variable genes were selected using the Seurat implementation in Scanpy with cohort-aware selection.

Scanorama [37] was applied using exact neighbor search, 100 integration dimensions, 50 neighbors, *σ* = 15, and *α* = 0.10. The integrated representation was stored separately from the log-normalized expression matrix, which was retained for gene-level analyses. A 50-neighbor graph constructed from the integrated representation was used for UMAP visualization and Leiden clustering [38].

Clustering was examined across resolutions, with marker expression and cohort composition used to assess the resulting populations (Supplementary Figure 2). The final annotation used resolution-0.8 cluster assignments with additional subclustering. Previously reviewed annotations were propagated through shared cells and cluster-level agreement, with marker inspection supporting the final assignments. The retained atlas contained nine annotated subtypes consolidated into six meta-clusters: Monocytes, Scavenging/C1q^+^, Resident/Quiescent, Inflammatory, Lipid-Stressed/Foam, and Fibrotic/Hypoxic macrophages. The final dataset comprised 34,533 cells from 50 libraries. Subtype-to-meta-cluster mappings, stored cluster identifiers, and marker-expression summaries are reported in Supplementary Table 3 and Supplementary Figure 8.

Integration was compared with Harmony [39], BBKNN [40], and scVI [41] on the same 20,576-cell subset, obtained by capping sampling at approximately 5,000 cells per cohort while preserving meta-cluster proportions. Evaluation included batch silhouette width, graph connectivity, and principal-component-regression-based batch correction scores (Supplementary Table 1). Metric definitions and the distinction between graph-based and embedding-based methods are described in the Supplementary Methods. Local cohort mixing in the final atlas was additionally quantified within each meta-cluster using the local inverse Simpson index, with perplexity 30 (Supplementary Table 2).

### 2.4 Optimal transport analysis

Optimal transport (OT) was used to compare the transcriptional distributions of the six meta-clusters. Population-level OT divergence and cell-level target-association scores were calculated separately: the former describes relative separation between populations, whereas the latter ranks source cells within a specified source–destination comparison.

#### 2.4.1 Pairwise divergence estimation

All 15 unordered meta-cluster pairs were evaluated in the 100-dimensional Scanorama representation. For the reference analysis, each population was sampled without replacement to a maximum of 5,000 cells. Uniform source and target marginals were used.

The cost matrix contained pairwise Euclidean distances and was divided by its maximum before fitting. Transport plans were estimated using the Sinkhorn algorithm implemented in Python Optimal Transport [42], with entropic regularization *ε* = 0.01 and a maximum of 2,000 iterations. After fitting, the transport cost was restored to the original distance scale:

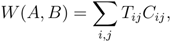

where *C* denotes the unscaled Euclidean cost matrix. The reported cost excludes the entropy term used to estimate the regularized transport plan.

Self-transport costs were computed using the same procedure. The self-cost-corrected statistic, referred to here as OT divergence, was

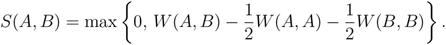

The diagonal was set to zero. Sampling procedures and random seeds are documented with the analysis code.

#### 2.4.2 Resampling assessment

Resampling variability was evaluated using 500 cell-level and 200 library-level bootstrap iterations. Cell-level iterations sampled cells with replacement within each state, with a maximum of 2,000 cells per state. Library-level iterations sampled libraries with replacement, concatenated their cells, and then sampled up to 2,000 cells per state without replacement from the resulting pool. Library resamples containing fewer than 50 cells in any state were excluded.

Divergences were recomputed in each iteration. Bootstrap means, standard deviations, and 2.5th–97.5th percentile intervals are reported in Supplementary Table 4 and Supplementary Figure 3. Because the reference and resampling analyses used different cell caps and sampling schemes, the bootstrap distributions are reported separately from the 5,000-cell reference estimates.

#### 2.4.3 Label-permutation null model

Meta-cluster labels were randomly permuted across cells 50 times, preserving the total number of cells assigned to each label. Divergences were recomputed with a maximum of 2,000 cells per state. Departure from this label-mixing null was summarized as

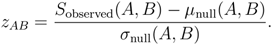

The implemented network-retention criterion was *|z_AB_| >* 2. This diagnostic assesses whether labeled populations differ from randomly mixed populations; it is not a test of biological interconversion.

#### 2.4.4 Connectivity transformation

For network visualization, retained divergences were transformed using

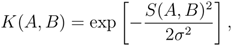

with *σ* set to half the median positive off-diagonal divergence among pairs passing the labelnull criterion. The diagonal and pairs failing that criterion were set to zero. Connectivity therefore increases with transcriptional proximity under the chosen representation; it is not a transition probability or an estimate of cellular flux.

#### 2.4.5 Sensitivity analyses

Leave-one-cohort-out analyses were performed for all seven cohorts. Each analysis removed one cohort from the existing integrated representation and repeated the OT calculations; integration itself was not refitted. Sensitivity was assessed using changes in raw divergence, pairwise ranks, and Spearman correlation with the full-cohort ranking (Supplementary Figure 4).

Regularization sensitivity was evaluated at *ε ∈ {*0.005, 0.01, 0.025, 0.05, 0.1, 0.5*}* using the same representation and reference sampling scheme (Supplementary Figure 9). Comparisons focused on divergence rankings rather than connectivity values rescaled with different runspecific bandwidths.

### 2.5 RNA velocity

RNA velocity was estimated separately for the 15 meta-cluster pairs using UniTVelo [24] in unified-time mode, with the designated source population supplied as the root for the primary analysis. Preprocessing of spliced and unspliced counts, gene filtering and neighborhood construction are described in Supplementary Methods.

Directional asymmetry was calculated from the scVelo transition matrix derived from each fitted velocity graph [18]. For each orientation, we calculated the proportion of interface cells whose transition mass towards the other population exceeded that population’s representation within their transition-matrix support. The difference between these proportions defined the signed asymmetry.

Significance was assessed using 100 sign-flip permutations, randomly changing the signs of individual cell velocity vectors and rebuilding the velocity graph and transition matrix. Twosided permutation *p*-values used a plus-one correction. The sign of the observed asymmetry defined the reported orientation when the unadjusted *p*-value was below 0.05; other pairs were classified as unresolved. The full interface definition, initialization-sensitivity analysis and pairwise results are provided in Supplementary Methods and Supplementary Table 5. UMAP stream plots were used for visualization rather than as the directionality statistic (Supplementary Figure 5).

### 2.6 OT target-association gradient analysis

Target-association gradients were calculated for 11 source–destination comparisons: Monocytes with each of the other five states; Scavenging and Resident macrophages with the Inflammatory state; and Scavenging, Resident, Inflammatory, and Foam macrophages with the Fibrotic state. Source–destination notation specifies the population being scored and its reference population; it does not by itself establish temporal direction.

#### 2.6.1 Per-cell target-association scoring

A separate transport plan was estimated for each comparison using all available source and target cells, the first 50 Scanorama dimensions, squared Euclidean costs normalized by their maximum, and *ε* = 0.01. Source marginals were uniform.

Target marginals were proportional to an inverse local-distance estimate. For each target cell, the mean Euclidean distance to the ten returned nearest neighbors in the target population was calculated; the query included the cell itself. Inverse mean distances, with a numerical offset of 10*^−^*^10^, were normalized to sum to one, defining target weights *b_j_*.

For source cell *i*, the association score was

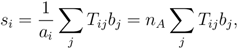

where *a_i_* = 1*/n_A_* and *n_A_* is the number of source cells. Thus, the score measures the targetdensity weight associated with a source cell under the conditional coupling. It is distinct from transport row mass, which is constrained by the source marginal.

#### 2.6.2 Count correction and quartile stratification

Within each source–destination comparison, scores were regressed against total RNA counts using ordinary least squares with an intercept. Residuals were retained as corrected scores, and source cells were divided into four approximately equal-sized quartiles, Q1–Q4. The saved corrected scores and quartile assignments were used consistently in downstream analyses.

Associations between total counts and scores before and after correction were summarized using Pearson and Spearman correlations (Supplementary Figure 10). No additional rankbased correction was applied to the current analyses.

#### 2.6.3 Gene-level expression contrasts

Expression contrasts used the retained natural-log-transformed, library-size-normalized RNA matrix. Individual expression values were returned to the linear normalized scale using expm1 before calculating means.

Comparisons were performed within each library and annotated source subtype. A stratum was eligible if it contained at least 20 source cells in both Q1 and Q4. For gene *g*, library *ℓ*, and subtype *h*, the effect was

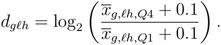

Eligible subtype effects were averaged equally within each library, and the median library effect was used for the principal summaries. Libraries therefore contributed equally regardless of their cell counts. Coverage and per-library effects are reported in Supplementary Table 6 and its associated source data.

These contrasts compare different cells at the extremes of an association gradient. Marker panels report effect sizes without pooled-cell significance tests.

#### 2.6.4 Marker panels and broader signatures

Canonical marker panels were curated using established macrophage markers and the atlas annotation. Panel development was exploratory; the displayed genes were not treated as a prospectively specified hypothesis set. The same source panel was retained across destinations where direct comparison was intended.

Broader signatures were derived by one-versus-rest Wilcoxon ranking for each metacluster. Among the top 300 ranked genes, candidates were required to have a reported log fold change above 0.5 and an adjusted *p*-value below 0.01. Genes matching mitochondrial or ribosomal prefixes, *MALAT1*, or *NEAT1* were excluded. The first 80 passing genes per state were considered, genes shared with another statés candidate list were removed, and up to 50 remaining genes were retained. The resulting signatures contained 50 genes for Monocytes, Scavenging, Resident, and Inflammatory macrophages, 45 for Foam cells, and 49 for Fibrotic macrophages.

Signature effects were calculated by averaging gene-level log_2_ fold changes within each eligible stratum, followed by equal subtype weighting within libraries and median aggregation across libraries. The S100-family summary used the same procedure for *S100A8*, *S100A9*, and *S100A12*. These summaries are means of gene-level effects, rather than fold changes of a summed expression score or averages of the displayed gene medians. Complete signature definitions and results are provided in Supplementary Table 6 and Supplementary Figure 11.

#### 2.6.5 Matched destination comparisons

To compare Scavenging associations with Inflammatory and Fibrotic macrophages, RNA analyses were restricted to library–subtype combinations eligible for both comparisons. Each comparison retained its own original quartile assignments. Signature effects were calculated using the same gene sets and aggregation procedure, then paired within libraries. This analysis used 25 matched strata from 19 libraries.

Additional target-resemblance analyses compared Q1 and Q4 source-cell profiles along the Mono–Fibro and Scav–Fibro target-association gradients with same-library target-state reference profiles in a shared 244-gene space. We quantified the reduction in distance from Q1 to Q4 and compared the reduction towards Fibrotic macrophages with that towards alternative states, using the same source cells and expression scaling for each comparison. Gene-panel construction, distance calculations, eligibility criteria and aggregation are described in the Supplementary Methods.

#### 2.6.6 Plasticity pattern characterization

Source-associated features were characterized by their attenuation, maintenance or reinforcement alongside destination-associated expression. Patterns were evaluated at both individual-gene and broader-signature levels. No numerical cutoff was used to assign every route to a fixed category, and near-zero effects were not treated as formal evidence of equivalence or complete identity retention. For Mono–Foam, effects were additionally inspected by library and monocyte subtype to distinguish consistent marker responses from near-zero summaries arising from opposing changes.

### 2.7 Held-out-feature and doublet-score analyses

Genes were considered strictly held out from integration if they were not selected as highly variable and had not been classified as highly variable in any cohort. For each source population, held-out genes detected in at least 20 cells and at least 1% of source cells were tested using the same library–subtype Q4-versus-Q1 expression contrast described above.

Before inference, library-level effects from eligible paired fresh and frozen preparations in the Jaiswal cohort were averaged with equal weight into one analysis unit; other libraries remained separate units. Eligibility, source-subtype weighting and quartile assignments were unchanged. Inference tested the mean analysis-unit effect using two-sided sign flips, preserving all gene effects from a unit together. Exact sign enumeration was used for 3–12 units and 9,999 Monte Carlo sign patterns for larger samples, with a plus-one correction for Monte Carlo *p*-values. Fewer than three units were considered insufficient for inference. Benjamini–Hochberg correction was applied across all tested gene–route combinations.

A second analysis measured cosine distance from source cells to the mean log-normalized expression of target cells from the same library. Feature panels comprised the top 100, top 500, or all expressed, variable held-out genes, ranked by variance across the combined source and target populations without reference to OT scores. At least 20 target cells and 20 valid source cells in each endpoint quartile were required. Within-stratum Spearman correlations between corrected OT scores and target distance were averaged within libraries and then within known paired preparations. Mean analysis-unit correlations were tested using the same sign-flip procedure, with Benjamini–Hochberg correction across route–panel tests. Supplementary Figure 6 displays median analysis-unit correlations.

SOLO scores among retained cells were examined across OT quartiles for the three inflammatory comparisons. Q4-minus-Q1 mean-score differences were calculated within eligible library–subtype strata, averaged within libraries, and then grouped using the same pairing rule. Inference used analysis-unit sign flips, with correction across the three routes; 10,000 bootstrap resamples of analysis units were used for intervals around mean effects. Supplementary Figure 7 retains descriptive pooled distributions and individual-library summaries. This analysis assessed score variation among retained cells and did not establish the absence of residual doublets.

### 2.8 CITE-seq protein analysis

Antibody-derived tag counts from the Bashore CITE-seq libraries were obtained from the processed data associated with GSE253902. RNA and protein measurements were matched by library and cell barcode, and analyses were restricted to cells present in the current RNA atlas and the corresponding gradient tables.

Protein counts were transformed using a per-cell centered log ratio:

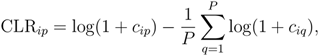

where *c_ip_*is the count for protein *p* in cell *i*, and *P* is the number of antibody features.

Protein effects were calculated as mean CLR in Q4 minus mean CLR in Q1 within each eligible library–subtype stratum, requiring at least 20 cells per endpoint. Subtype effects were averaged within libraries, and median library effects were displayed. Protein differences were not expressed as RNA log_2_ fold changes.

The panel included source-associated proteins, activation-associated proteins, lineage controls, and available isotype controls. The Scavenging-to-Inflammatory analysis contained three eligible libraries. The paired Inflammatory-versus-Fibrotic comparison used the two libraries with eligible matched source-subtype strata for both destinations.

For the descriptive same-cell analysis, a source-associated score was calculated as the mean CLR of CD163, CD206, and MERTK, and an activation-associated score as the mean CLR of HLA-DR and CD38. Within each library, the Q1 median of each score defined its reference threshold. The percentage of cells exceeding both thresholds was calculated separately in Q1 and Q4. These thresholds summarize joint expression and were not treated as validated functional gates.

### 2.9 Transcription-factor activity inference

Transcription-factor activities were inferred from the current atlas log-normalized expression matrix using the univariate linear model implemented in decoupleR v1.8.0 [43], with human signed regulons from CollecTRI [44]. Complexes were retained, and a minimum of five measured target genes per regulator was required. The network used for inference was retained with the analysis outputs.

For each route, mean Q4-minus-Q1 activity differences were calculated within eligible library–subtype strata and averaged within libraries. Median library effects were reported. Two-sided Wilcoxon signed-rank tests were applied to library effects where at least three libraries were available, followed by Benjamini–Hochberg correction across all tested regulator– route combinations. The complete results are provided in Supplementary Table 7; effectranked display selections were not restricted to significant regulators. Inferred activities reflect regulon-associated expression and were not interpreted as direct measurements of regulator expression or causal activity.

### 2.10 Clinical association analysis

Clinical analyses used the Bashore cohort, with 18 mapped patients: 11 symptomatic and seven asymptomatic. Libraries were linked to patients and symptom status using the library manifest and BioSample metadata (Supplementary Table 8).

For each of the Monocyte-to-Inflammatory and Scavenging-to-Inflammatory comparisons, the 90th percentile of corrected association scores was calculated across mapped Bashore source cells. Each patient’s outcome was the fraction of their source cells at or above this common, route-specific threshold. The threshold was therefore defined across the mapped cohort, rather than separately within each patient or across the complete atlas. Patients with source cells and known symptom status contributed one value per comparison.

The test statistic was the difference between symptomatic and asymptomatic patient medians. Symptom labels were permuted among patients 20,000 times, preserving group sizes. One-sided *p*-values evaluated higher fractions in symptomatic patients and used a plusone correction. Reported clinical *p*-values were unadjusted. These analyses were exploratory and were not considered confirmatory tests or assessments of prospective prediction.

### 2.11 Application to human liver macrophages

Processed human liver expression matrices from Ramachandran et al. [26] were obtained from GSE136103, including healthy and cirrhotic liver samples. Cells with fewer than 500 detected genes or at least 20% mitochondrial counts were excluded. Expression was normalized to 10,000 counts per cell and log-transformed.

Integration was performed by library using 2,000 highly variable genes and a 50-dimensional Scanorama representation. Broad lineage annotation identified the myeloid compartment, which was re-integrated and clustered. Marker-guided review and additional subclustering defined tissue monocytes, Kupffer cells, C1q^+^/MHC-II macrophages, and scar-associated macrophages. Dendritic and nonmyeloid populations were excluded, yielding 8,536 retained cells.

Pairwise OT and target-association analyses used the corresponding liver representation and state definitions. Gene-expression contrasts for tissue monocytes and Kupffer cells in relation to scar-associated macrophages used the same linearized-expression log_2_ fold-change estimator as the plaque analysis. Comparisons required at least 20 Q1 and 20 Q4 cells within a library and were summarized by the median library effect, without the plaque-specific subtype stratification. Library coverage and healthy/cirrhotic contributions were recorded separately. The liver analysis evaluated applicability of the comparison strategy in another tissue and did not include RNA velocity. For CD9, expression and the fraction of cells with detectable expression were additionally summarized by library and association quartile. CD9 expression in annotated Kupffer cells and SAM was compared descriptively within the same libraries, with SAM references containing fewer than 20 cells flagged as having limited coverage.

### 2.12 Statistical analysis and reporting

Cells, library–subtype strata, libraries, and patients were treated as distinct analysis units. RNA, protein, and regulatory contrasts used library-level summaries; clinical comparisons used patients. A targeted sensitivity analysis grouped eligible paired fresh and frozen preparations from the Jaiswal cohort by donor, averaging their saved library-level effects with equal weight while leaving other libraries unchanged. Broad-signature summaries and transcriptionfactor Wilcoxon tests were recomputed, with Benjamini–Hochberg correction across all tested regulator–route combinations. This analysis assessed sensitivity to the known Jaiswal pairing rather than constituting a complete donor-level reanalysis.

The testing procedures and multiplicity corrections were defined separately for each analysis. Gene and protein marker panels were primarily descriptive; held-out-gene, held-outdistance, SOLO-score, and transcription-factor analyses used the testing families specified above. Velocity and clinical permutation *p*-values were reported as unadjusted. Missing or ineligible comparisons were retained as unavailable rather than assigned zero effects. Detailed gene sets, coverage, sensitivity results, and supporting numerical outputs are provided in the Supplementary Materials.

### 2.13 Use of AI-assisted tools

Codex (OpenAI; Astra) and Claude (Anthropic; Opus 4.8) were used to assist with manuscript preparation and computational implementation. The iterative instructions are summarized here: revise supplied text for clarity, continuity, grammar and formatting while preserving its scientific meaning; develop and debug analysis code; prepare plotting and table-export code; and review consistency between analytical outputs, their interpretation and manuscript statements. These descriptions summarize multiple interactions.

The authors reviewed and verified the resulting text, code and analytical outputs, made the final methodological and interpretative decisions, and take responsibility for the work.

## 3 Results

### 3.1 A unified single-cell atlas of macrophages in human atherosclerotic plaques

To comprehensively map macrophage heterogeneity in human atherosclerosis, we gathered and reprocessed raw scRNA-seq data from seven publicly available datasets: Alsaigh [27], Bashore [25], Pan [28], Jaiswal [29], Pauli [30], Fernandez [12], and Wirka [31] (Fig. 1A). All datasets were reprocessed from FASTQ files using a unified preprocessing procedure including ambient RNA removal with CellBender [33], doublet detection with SOLO [35], and per-sample quality assessment (Supplementary Figure 1; see Methods). Following quality control and annotation, the final atlas comprised 34,533 monocytes and macrophages from 50 libraries across the seven cohorts.

**Figure 1:**
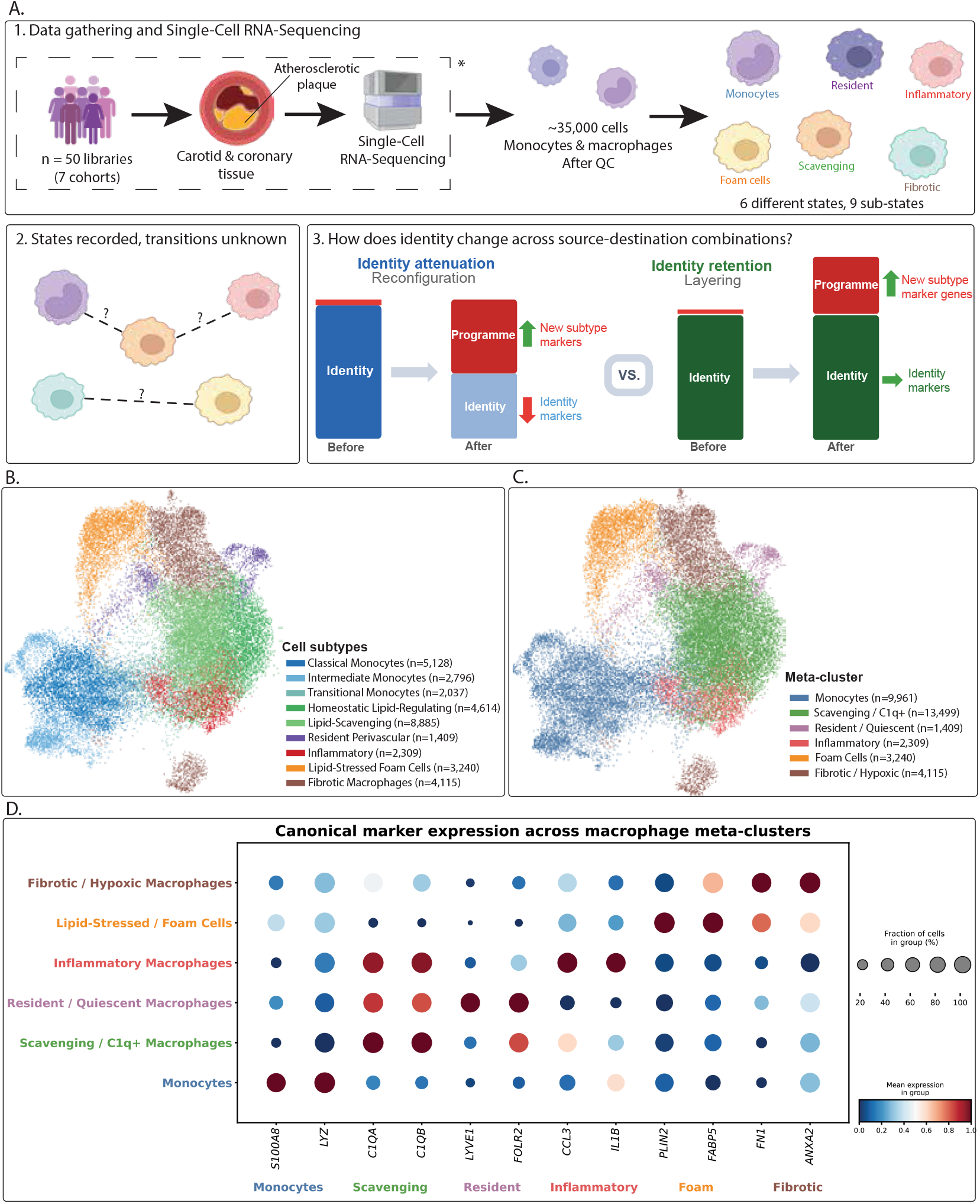
A cross-cohort atlas for comparing macrophage states in human atherosclerosis. **(A)** Overview of the study design. Single-cell RNA sequencing data from seven publicly available cohorts of human atherosclerotic plaques (Alsaigh, Bashore, Pan, Jaiswal, Pauli, Fernandez, Wirka; including carotid and coronary specimens) were reprocessed from raw FASTQ files through a unified bioinformatics pipeline (see Methods). The asterisk denotes steps performed by the original cohort consortia (patient recruitment, tissue collection, and droplet-based sequencing). Following quality control and annotation, the final atlas comprised 34,533 monocytes and macrophages from 50 libraries. **(2 & 3)** Conceptual illustration of the study questions: how macrophage states are connected, and whether destination-associated programs accompany attenuation, retention or reinforcement of source-associated features. Selective attenuation and maintenance of source-associated features are illustrated as candidate patterns to be examined across source–destination combinations. **(B)** UMAP embedding colored by nine annotated cell subtypes, with cell counts indicated in the legend. **(C)** The same UMAP embedding colored by the six macrophage metaclusters used for the plaque analyses: Monocytes (*n* = 9,961), Scavenging/C1q^+^ (*n* = 13,499), Resident/Quiescent (*n* = 1,409), Inflammatory (*n* = 2,309), Lipid-Stressed/Foam Cells (*n* = 3,240), and Fibrotic/Hypoxic (*n* = 4,115). **(D)** Dot plot of canonical marker gene expression across meta-clusters, including the resident-associated markers *LYVE1* and *FOLR2*. Dot size indicates the percentage of cells expressing each gene; color indicates mean expression scaled from 0 to 1 for each gene.

Batch-effect correction was performed using Scanorama [37] and benchmarked against Harmony [39], BBKNN [40], and scVI [41] on the same subset of 20,576 cells from all 50 libraries and seven cohorts. Sampling was capped at 5,000 cells per cohort and stratified by meta-cluster within each cohort to preserve population proportions (Supplementary Tables 1–2; Supplementary Methods).

Across the four methods, batch average silhouette width (ASW) scores ranged from 0.911 to 0.934 and graph connectivity scores from 0.985 to 0.998. Scanorama had the highest observed scores for these two metrics (0.934 and 0.998, respectively), whereas Harmony had a higher principal component regression (PCR) batch-removal score (0.655 versus 0.424) and the highest composite batch score, calculated as the mean of the three metrics (0.855 versus 0.786 for Scanorama). Scanorama was retained as the shared representation for downstream analyses, given its high batch ASW and graph connectivity. Its integrated embedding was used to construct cell neighbourhoods and quantify relationships between populations, while gene-expression and spliced/unspliced measurements were retained separately.

Clustering was examined across Leiden resolutions (Supplementary Figure 2; Supplementary Methods). The final atlas comprised nine annotated cell subtypes (Fig. 1B), which were grouped into six meta-clusters by manual annotation against canonical macrophage marker panels established in the plaque scRNA-seq literature [11, 12, 15, 16, 45, 46], combining subtypes with related expression profiles. The marker panels with per-subtype expression values are provided in Supplementary Table 3 and Supplementary Figure 8. The six meta-clusters were Monocytes (Mono; *n* = 9,961), Scavenging/C1q^+^ Macrophages (Scav; *n* = 13,499), Resident/Quiescent Macrophages (Res; *n* = 1,409), Inflammatory Macrophages (Inflam; *n* = 2,309), Lipid-Stressed/Foam Cells (Foam; *n* = 3,240), and Fibrotic/Hypoxic Macrophages (Fibro; *n* = 4,115) (Fig. 1C).

The Scavenging meta-cluster is named for its coordinate expression of the complement component C1q genes (*C1QA*, *C1QB*) alongside *APOE* and *SELENOP* (Fig. 1D; Supplementary Table 3). Throughout, capitalised names refer to the six expression-defined metaclusters described here, whereas lowercase usage refers to macrophage phenotypes in the general sense.

Each meta-cluster displayed a coherent marker profile (Fig. 1D), consistent with established macrophage populations in human atherosclerotic plaques [15] and the heterogeneity described in the integrated human plaque atlas of Träuble et al. [16]. Building on this description of plaque heterogeneity, we used our atlas as the basis for a method that quantifies relationships between macrophage states and characterizes changes in source- and destinationassociated gene programs (Fig. 2A). This establishes a common setting for comparing different source populations associated with the same destination, and the same source population associated with different destinations.

**Figure 2:**
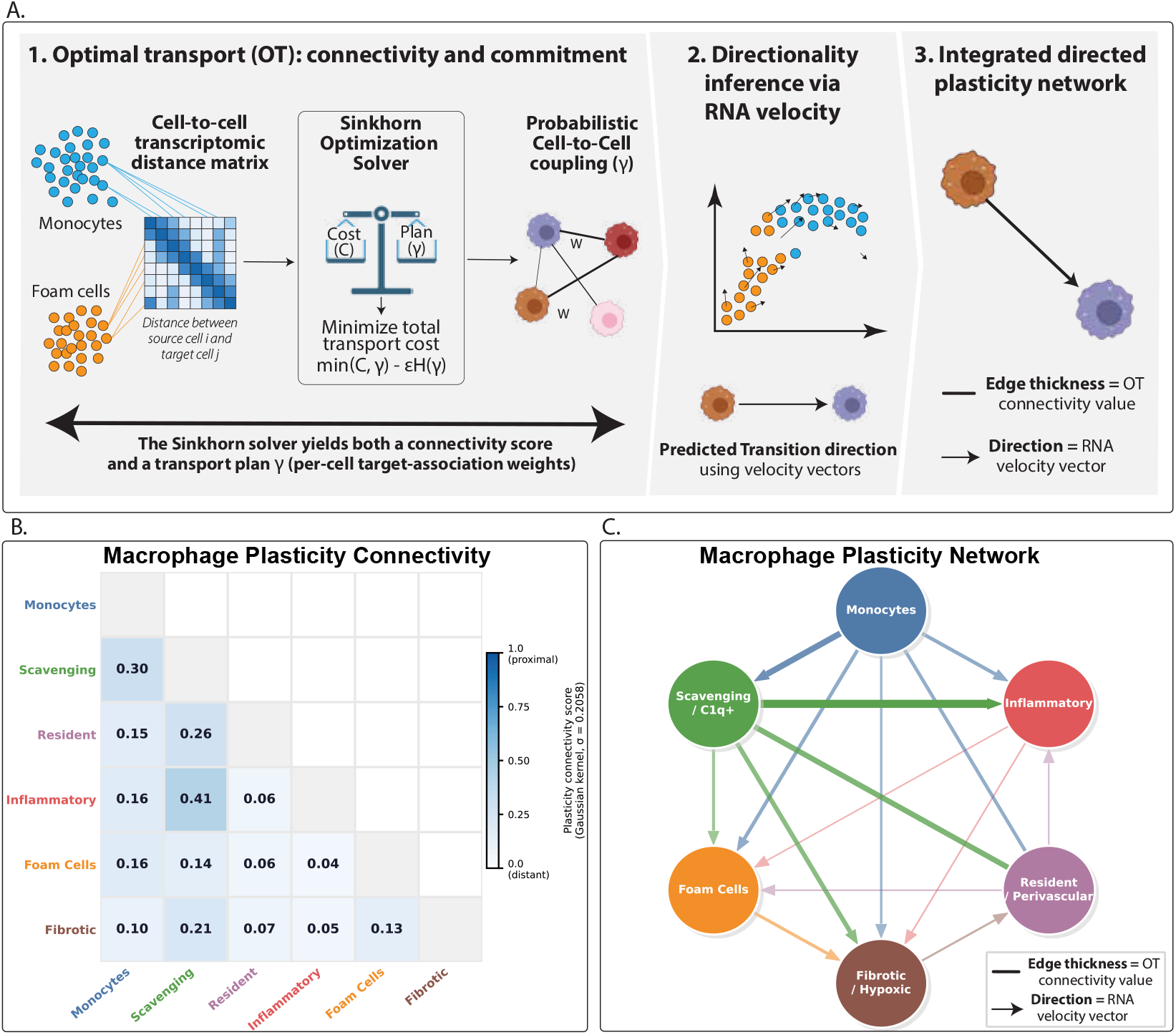
Optimal transport maps macrophage state connectivity, with model-dependent RNA velocity orientations. **(A)** Overview of the computational method. For each pair of macrophage states, optimal transport uses transcriptomic distances between source and target cells to estimate population-level connectivity. A separate cell-level transport calculation yields target-association scores for the expression analyses in Figs. 3–5. Pairwise RNA velocity models provide complementary estimates of orientation under specified initialization assumptions. **(B)** Pairwise OT-based connectivity scores among the six macrophage meta-clusters. Higher values indicate closer cell-population distributions in the integrated transcriptomic space. These scores describe relative relatedness rather than transition probabilities. Label-permutation results are reported in Supplementary Table 4. **(C)** Macrophage connectivity network. Edge thickness represents OT-based connectivity, and arrowheads indicate orientations from the source-root UniTVelo analysis and its sign-flip test. Mono–Res and Scav–Res lacked significant asymmetry and are shown without arrowheads. Orientations are conditional on the fitted model; initialization sensitivity is reported in Supplementary Table 5 and the Supplementary Methods.

### 3.2 Optimal transport and RNA velocity characterize relationships between macrophage states

To systematically map transcriptomic relationships between macrophage states, we computed pairwise OT-based divergences between all meta-cluster pairs on the Scanorama-corrected embedding (Fig. 2A; see Methods). For each pair of macrophage states, we constructed a cost matrix representing the transcriptomic distance between source and target cells. The Sinkhorn algorithm [47] then identified an entropically regularized transport plan matching the two populations. Transport costs, corrected for within-population self-costs, yielded a divergence summarising their distributional separation. This approach compares cell-population distributions while retaining their internal heterogeneity. Entropic regularization made these comparisons computationally tractable, with up to 5,000 cells per state in the population-level analysis.

A low divergence indicates that two states are close in the integrated transcriptomic space, whereas a high divergence indicates greater separation. Divergences were converted to connectivity scores using a Gaussian kernel, with higher scores indicating closer cell-population distributions (Fig. 2B; see Methods). All 15 pairs showed greater separation than populations formed by shuffled labels (Supplementary Table 4). This supports the transcriptional distinction between the annotated states, while connectivity provides a relative measure of their relatedness.

The resulting network (Fig. 2C) places each meta-cluster in relation to multiple other macrophage states. The strongest OT connection was Scav–Inflam (connectivity = 0.412), followed by Mono–Scav (0.298) and Res–Scav (0.263). These relationships identify closely related populations for detailed analysis but do not quantify their contributions to the formation of other states.

We used connectivity to prioritize candidate relationships for further characterization, without interpreting it as a rate or probability of interconversion. Robustness was assessed using cell-level and library-level bootstrap resampling, label permutation, and leave-one-cohort-out (LOCO) analyses across all seven cohorts (Supplementary Table 4; Supplementary Figures 3–4). Scav–Inflam had the lowest divergence in the full analysis and retained this rank in every LOCO analysis. Overall rank correlations with the full analysis ranged from 0.843 to 0.996, although individual relationships were more sensitive to cohort composition, with a maximum shift of seven positions for Foam–Fibro upon Bashore exclusion. Removing the Wirka cohort changed divergences by approximately 0.006 on average and at most 0.022. These results assess sensitivity to cohort composition without establishing vascular-bed independence. We also varied the regularization parameter *ɛ*, which controls how broadly transport is distributed among possible cell matches, with larger values producing smoother transport plans. Pairwise rankings were closely preserved across *ɛ* = 0.005–0.025, but changed more substantially at higher values (Supplementary Figure 9).

**Figure 3:**
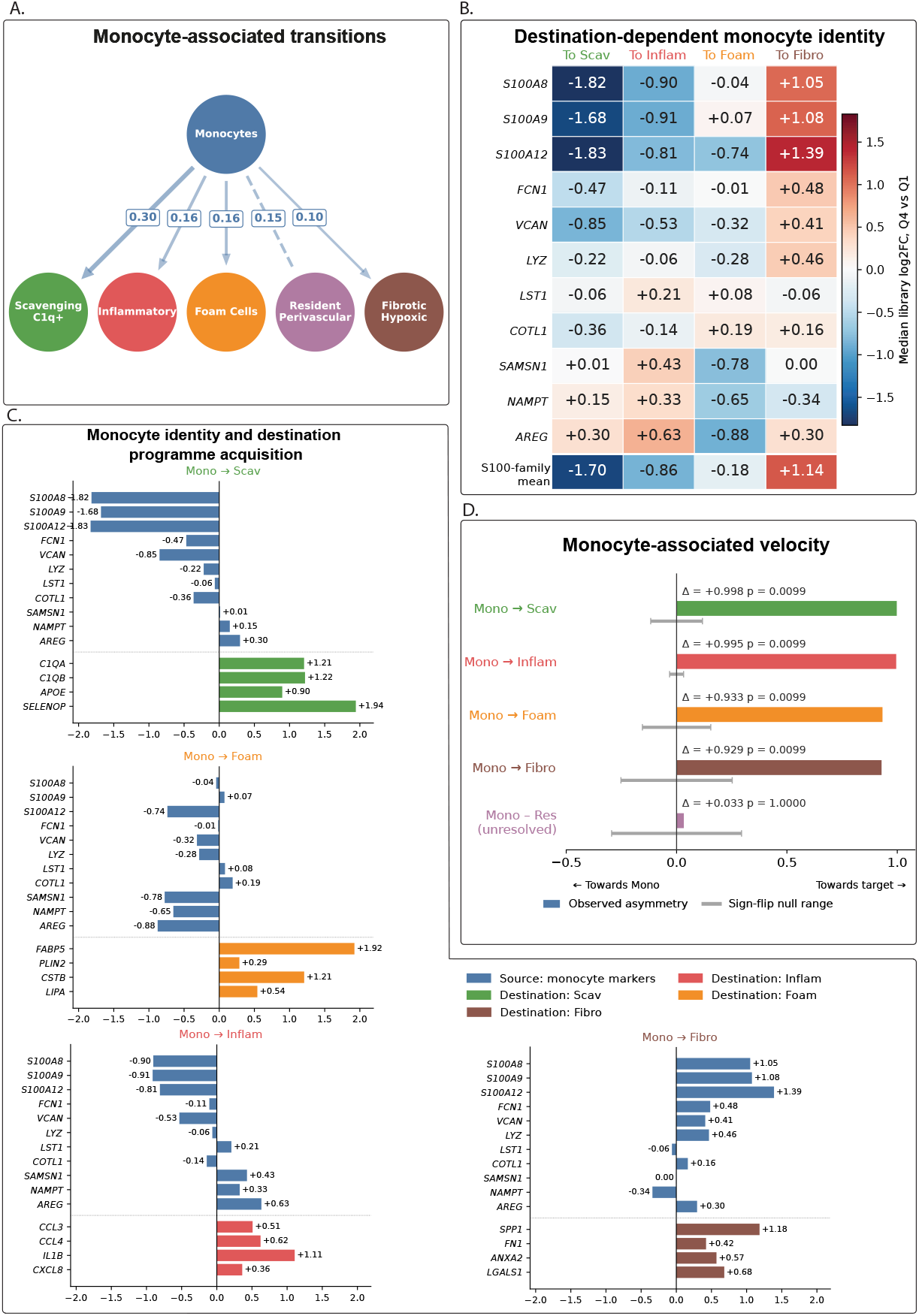
Destination-associated changes in monocyte identity (Biological Axis A). **(A)** Monocyte-associated subnetwork. Edge labels indicate OT-based connectivity; arrowheads represent UniTVelo orientations. Mono–Res remained unresolved. **(B)** Q4-versus-Q1 changes in 11 monocyte-associated genes across Scav, Inflam, Foam and Fibro associations. Q1 and Q4 denote the lowest and highest OT target-association quartiles. Values are median library log_2_ fold changes from linearised normalised expression (pseudocount, 0.1). Library/subtype comparisons required at least 20 cells per quartile; subtype effects were averaged within libraries before calculating the median across libraries. Coverage (libraries/library–subtype combinations): Scav, 17/22; Inflam, 18/23; Foam, 10/16; Fibro, 12/14. The S100-family row averages *S100A8*, *S100A9* and *S100A12* effects before library aggregation. **(C)** The same monocyte-marker changes alongside four destination-associated markers per target. Source markers are blue; destination markers use the corresponding state colour. Positive values indicate higher expression in Q4. **(D)** Signed velocity asymmetry at monocyte–macrophage interfaces. Coloured bars show observed asymmetry, with positive values indicating orientation towards the macrophage state. Grey intervals show symmetric sign-flip null ranges (plus or minus the 95th percentile of absolute null asymmetry), not confidence intervals. Unadjusted, two-sided *p* values use 100 randomizations and a plus-one correction. All velocity results use monocyte-root initialization; details and initialization sensitivity are reported in Supplementary Table 5.

**Figure 4:**
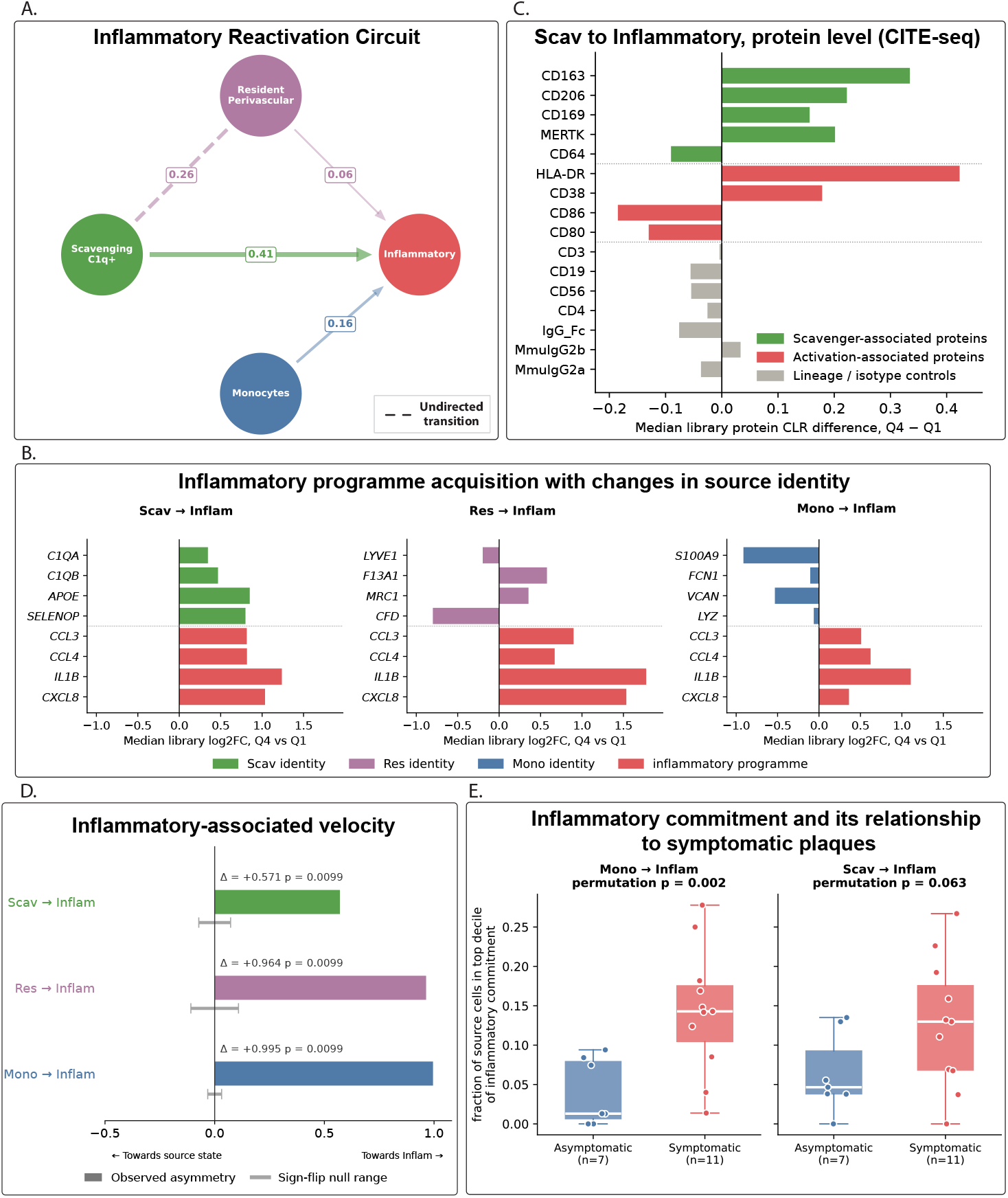
Inflammatory programming across distinct source states, with protein corroboration and clinical association (Biological Axis B). **(A)** Inflammatory-axis subnetwork. Edge labels show OT connectivity; arrowheads show source-root UniTVelo orientations. **(B)** Source-marker and inflammatory-marker changes between the highest and lowest OT target-association quartiles (Q4 versus Q1). Bars show median library log_2_ fold changes from linearised normalised expression (pseudocount, 0.1). Comparisons required *≥* 20 cells per quartile within library/subtype; subtype effects were averaged within libraries. Coverage (libraries/library–subtype combinations): Scav, 26/33; Res, 5/5; Mono, 18/23. **(C)** Scav–Inflam protein changes within the Lipid-Scavenging (C1q^+^/HLA-DR^+^) subtype in three Bashore libraries. Bars show median library Q4-minus-Q1 differences in mean centred log-ratio (CLR) abundance. Q1/Q4 counts: GSM8029956, 57/180; GSM8029958, 99/115; GSM8029960, 130/184. Green denotes scavenger-associated proteins, red activation-associated proteins, and grey lineage markers and comparison antibodies. **(D)** Signed velocity asymmetry for the three inflammatory routes under source-root initialization. Positive values favour Inflam; grey intervals show symmetric sign-flip null ranges (plus or minus the 95th percentile of absolute null asymmetry), not confidence intervals. Unadjusted, two-sided *p* values use 100 randomizations with a plus-one correction; sensitivity is reported in Supplementary Table 5. **(E)** Bashore clinical comparison (7 asymptomatic, 11 symptomatic patients). Points show patient-specific fractions at or above a common source-specific threshold: the 90th percentile of corrected inflammatory target-association scores among mapped Bashore source cells. Boxes show medians and interquartile ranges; whiskers extend to observations within 1.5 times that range. Unadjusted, one-sided patient-label permutation tests compare symptomatic-minus-asymptomatic differences in medians (20,000 permutations).

To connect population-level relatedness to expression changes within source states, we derived cell-level target-association scores from separate OT computations without subsampling, using uniform source marginals and density-weighted target marginals (see Methods). For each source cell, the score summarizes the target density weights reached by its conditional transport coupling. After correction for total UMI counts (Supplementary Figure 10), source cells were stratified into association quartiles. Comparing the highest and lowest quartiles (Q4 versus Q1) allowed us to characterize how source markers and destination-associated programmes vary across these cross-sectional gradients (Figs. 3–5). This provides a common basis for comparing selective attenuation, retention or reinforcement of source-associated features across different source–destination relationships.

**Figure 5:**
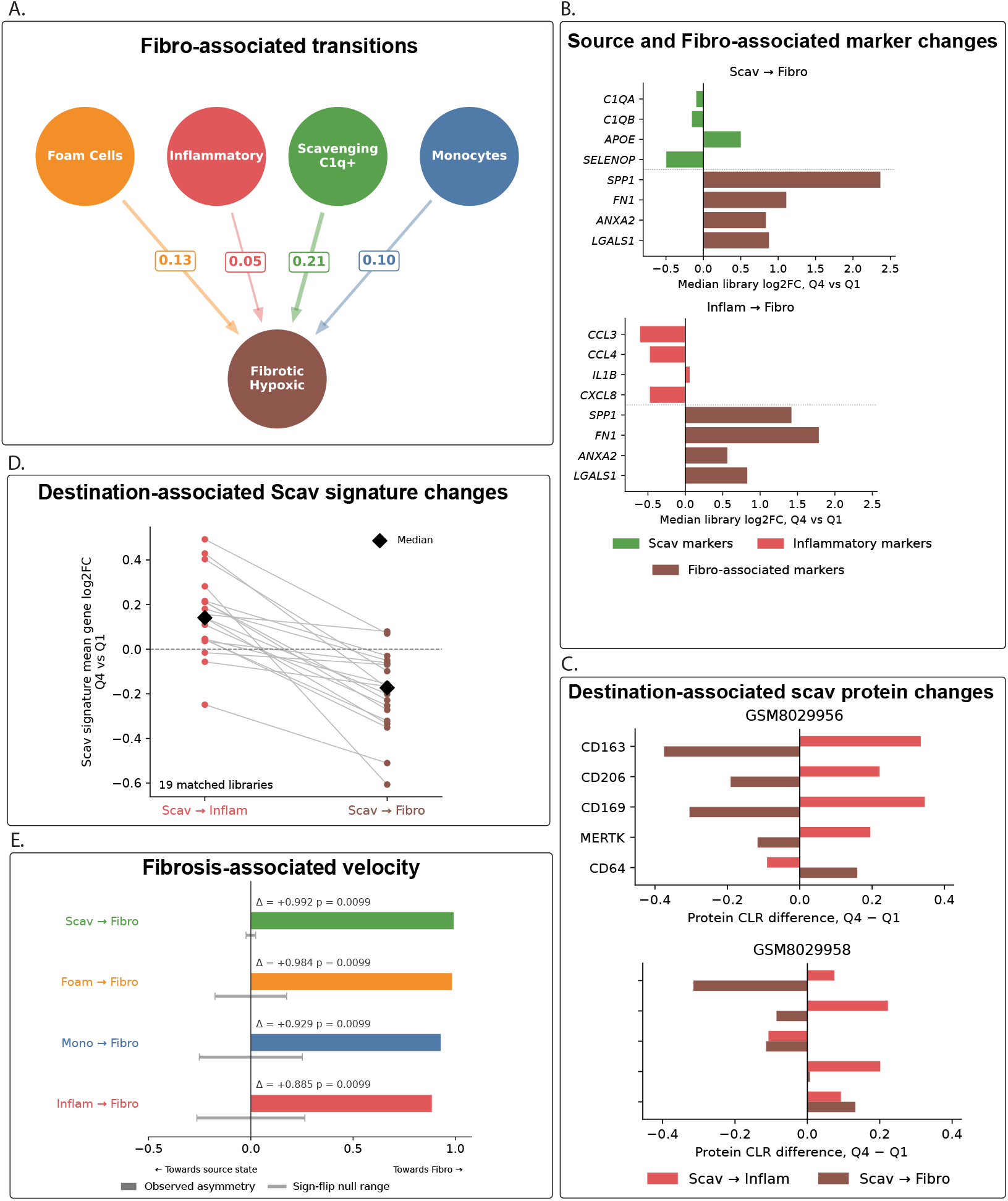
Source-identity remodelling along Fibrotic association (Biological Axis C). **(A)** Fibro-associated subnetwork. Edge labels indicate OT connectivity; arrowheads represent source-root UniTVelo orientations. **(B)** Source-marker and Fibro-associated marker changes along Scav–Fibro and Inflam–Fibro association. Bars show median library log_2_ fold changes between the highest and lowest OT target-association quartiles (Q4 versus Q1), using linearised normalised expression (pseudocount, 0.1). Comparisons required *≥* 20 cells per quartile within library/subtype; subtype effects were averaged within libraries. Coverage (libraries/library–subtype combinations): Scav, 21/28; Inflam, 6/6. Source markers use state colours; Fibro-associated markers are brown. **(C)** Scavenger-associated protein changes in two matched Bashore libraries, within the Lipid-Scavenging (C1q^+^/HLA-DR^+^) subtype. Bars show Q4-minus-Q1 differences in mean centred log-ratio (CLR) abundance for Scav–Inflam (red) and Scav–Fibro (brown). **(D)** Broader Scav RNA-signature changes across both associations in 19 matched libraries/25 library–subtype combinations. Points show library effects, averaging gene log_2_ fold changes within subtypes, then subtype effects within libraries. Lines connect matched library effects; black diamonds show medians across libraries. **(E)** Signed velocity asymmetry for the four pairs in A, using Scav, Foam, Mono or Inflam as the source root. Positive values favour Fibro. Grey intervals show symmetric sign-flip null ranges (plus or minus the 95th percentile of absolute null asymmetry), not confidence intervals. Unadjusted, two-sided *p* values use 100 randomizations with a plus-one correction. Results for all 15 pairs and initialization sensitivity are reported in Supplementary Table 5.

Because the population-level OT divergence is symmetric and does not encode directionality, we used UniTVelo [24] to infer RNA velocity for each pairwise meta-cluster combination and assessed orientation using a bidirectional asymmetry test (Methods; Supplementary Methods; Supplementary Table 5; Supplementary Figure 5). For each population, we calculated the fraction of interface cells whose velocity-derived transition mass towards the other population exceeded its representation among supported neighbours. The difference between these fractions defines the signed asymmetry, with significance assessed against a null obtained by randomly reversing cell velocity vectors.

In the primary UniTVelo analysis, 13 of the 15 pairs showed significant asymmetry: 12 followed the designated source-to-target orientation and Res–Fibro favoured Fibro-to-Res (Fig. 2C). Mono–Res and Scav–Res remained unresolved. Orientations depended on initialization, and are therefore interpreted as complementary estimates of direction; the complete sensitivity analysis is reported in Supplementary Table 5. The expression-gradient analyses provide comparisons of source cells with differing target association.

We organized the subsequent analyses around monocyte-associated, inflammatory, and Fibrotic-associated relationships.

### 3.3 Biological Axis A: Destination-associated changes in monocyte identity

The first axis characterizes how monocyte identity varies with association to different macrophage states (Fig. 3A). The atlas contained 9,961 monocytes. Their strongest OT connection was with Scavenging/C1q^+^ macrophages (connectivity = 0.298), followed by Inflammatory macrophages and Foam Cells (both 0.160), Resident/Quiescent macrophages (0.151), and Fibrotic/Hypoxic macrophages (0.103). These relationships motivated a comparison of monocyte expression across different destination associations, rather than assuming a single trajectory of source-identity loss.

Connectivity robustness was assessed across all seven LOCO analyses (Supplementary Table 4). For example, the Mono–Foam connectivity was similar with and without the Alsaigh cohort (0.160 and approximately 0.153, respectively). In the primary UniTVelo analysis, the Scav, Inflam, Foam and Fibro pairs were oriented towards the macrophage population, whereas Mono–Res remained unresolved (Supplementary Table 5).

#### 3.3.1 OT target-association gradients reveal selective changes in monocyte source and destination programmes

Using the cell-level target-association scores described above, monocytes were divided into quartiles after correction for total UMI counts. We compared Q4 with Q1 for the same 11 monocyte-associated genes across the Scav, Inflam, Foam and Fibro gradients, together with markers of each destination state (Fig. 3B,C; Supplementary Table 6). Log_2_ fold changes were calculated within eligible library/subtype combinations, averaged across eligible subtypes within each library, and summarized by the median across libraries. Between 10 and 18 libraries contributed to each destination comparison, reducing the influence of differences in library and monocyte-subtype composition. Marker panels combine established identity genes with cluster-enrichment information (Methods, “Marker panels and broader signatures”; Supplementary Table 3).

Monocytes with greater Scav or Inflam association showed lower expression of the S100-family markers (median library mean-gene log_2_FC: *−*1.70 and *−*0.86, respectively; Fig. 3B).

*FCN1* and *VCAN* also decreased along both gradients, although their effect sizes differed between destinations. These changes accompanied higher expression of the respective destination markers: *C1QA* (+1.21), *C1QB* (+1.22), *APOE* (+0.90) and *SELENOP* (+1.94) along Scav association, and *CCL3* (+0.51), *CCL4* (+0.62), *IL1B* (+1.11) and *CXCL8* (+0.36) along Inflam association (Fig. 3C). Other monocyte-associated genes were preserved or in-creased; for example, *NAMPT*, *SAMSN1* and *AREG* increased along Inflam association. Thus, destination-associated expression accompanied selective attenuation of source features rather than uniform loss of monocyte-associated expression.

The Foam-associated gradient showed a different pattern. The S100-family summary decreased modestly overall (*−*0.18), with near-zero median changes in *S100A8* and *S100A9* but lower *S100A12* expression (*−*0.74). However, the near-zero medians did not reflect consistent marker maintenance: *S100A8*, *S100A9* and *FCN1* each increased in five libraries and decreased in five. Subtype-stratified inspection likewise revealed heterogeneous marker responses, while the broader monocyte signature had a negative median change in each of the three monocyte subtypes. *NAMPT*, *SAMSN1* and *AREG* decreased (*−*0.65, *−*0.78 and *−*0.88, respectively), alongside increased *FABP5* (+1.92), *PLIN2* (+0.29), *CSTB* (+1.21) and *LIPA* (+0.54). All four displayed Foam markers increased in every eligible library. Thus, Foam association accompanied consistent destination-marker enrichment and heterogeneous changes within the monocyte programme, rather than general preservation of monocyte identity.

Fibro association was accompanied by reinforcement of several monocyte markers. The S100-family summary increased (+1.14), together with *FCN1* (+0.48), *VCAN* (+0.41) and *LYZ* (+0.46), while the selected destination markers *SPP1*, *FN1*, *ANXA2* and *LGALS1* also increased (+1.18, +0.42, +0.57 and +0.68, respectively). *NAMPT* decreased modestly, *SAMSN1* was close to unchanged, and *AREG* increased, providing no evidence for a uniform reciprocal loss of an activation programme. In an additional analysis restricted within monocyte subtypes, Q4 cells were closer to the Fibro reference than Q1 cells in all seven eligible libraries on a shared 244-gene panel. However, the additional reduction in distance towards Fibro relative to Foam was small in the five matched libraries (median paired advantage 0.006; Fibro favoured in three of five). This supports increased Fibro resemblance while limiting a claim of exclusive Fibro destination specificity.

Analysis of broader state-associated signatures extended these comparisons beyond the selected marker panels (Supplementary Figure 11). Destination-signature changes were positive in every eligible library for each of the five monocyte-associated routes. The broader monocyte signature decreased along Scav and Foam association (median library mean-gene log_2_FC of *−*0.19 and *−*0.26, respectively), but increased along Inflam association (+0.14), with positive changes in 15 of 18 libraries. Its median changes along Res and Fibro association were smaller (both approximately +0.07). In particular, the increase in the broader monocyte signature along Inflam association, despite lower S100-family and *VCAN* expression, supports selective remodelling within the source programme rather than uniform loss of monocyte identity.

To assess whether the expression associations extended beyond the features used to construct the embedding, we examined genes excluded from highly variable gene selection in every cohort and from the final embedding feature set (Supplementary Figure 6; Supplementary Methods). Four of the five monocyte-associated routes showed Q4-versus-Q1 expression differences in held-out genes, with 91–342 significant genes per supported route after library/subtype matching, grouping known paired preparations, and global Benjamini– Hochberg correction (*q <* 0.05). Mono–Foam did not retain gene-level support at this adjusted threshold. These results show that OT-associated expression differences extend beyond direct reuse of the embedding features.

We also assessed whether higher OT association accompanied smaller cosine distance to the same-library target population in held-out-gene expression space, using the top 100, top 500 or all eligible variable genes. After grouping known paired preparations, median correlations were negative for all five monocyte-associated routes. Mono–Fibro retained adjusted statistical support across all three panels, whereas Mono–Scav and Mono–Inflam retained support only with the top 100 genes. Mono–Foam did not reach the adjusted threshold, and the two eligible Mono–Res libraries represented one paired donor and were assessed as a single descriptive unit. Thus, held-out expression supported target resemblance for several monocyte associations, with the extent of support varying across destinations, feature panels and available analysis units (Supplementary Figure 6).

Taken together, the monocyte-associated gradients show that destination programmes are accompanied by different patterns of source-marker attenuation, retention and reinforcement. Because these are cross-sectional comparisons, they do not establish that an initial activation state determines a subsequent fate. This destination-dependent variation motivates the complementary comparison in the next axis: whether different source populations show similar or distinct identity changes when associated with the same inflammatory state.

### 3.4 Biological Axis B: Inflammatory programming across distinct source states

The second axis examines inflammatory programming across Scavenging/C1q^+^ macrophages, Resident/Quiescent macrophages and monocytes (Fig. 4A). Whereas the preceding axis compared one source population across different destinations, this analysis asks how different source identities relate to the same inflammatory state. Scav–Inflam was the strongest OT connection in the atlas (connectivity = 0.412), followed within this axis by Mono–Inflam (0.160) and Res–Inflam (0.061). Resident and Scavenging macrophages were also closely related transcriptionally (connectivity = 0.263). These values quantify population-level transcriptomic relatedness; they do not estimate the proportion of inflammatory macrophages derived from each source.

The Scav–Inflam connection retained the highest rank across all seven LOCO analyses (Supplementary Table 4). Its connectivity was approximately 0.371 after Alsaigh exclusion, compared with 0.412 in the full analysis. The primary UniTVelo analysis oriented all three inflammatory pairs towards Inflam, while Res–Scav remained unresolved (Supplementary Table 5). We next examined how inflammatory association was accompanied by changes in source-identity features at the RNA and protein levels.

#### 3.4.1 Inflammatory programmes accompany different changes in source identity

To assess whether inflammatory programming is accompanied by a common pattern of sourceidentity change, we examined the same four inflammatory markers (*CCL3*, *CCL4*, *IL1B* and *CXCL8*) across all three source populations, alongside their respective identity markers (Fig. 4B; Supplementary Table 6). Using the library/subtype-balanced comparison described above, all four inflammatory genes had positive median Q4-versus-Q1 log_2_ fold changes in each population. For example, *IL1B* increased by +1.24 in Scav, +1.78 in Res and +1.11 in Mono. This shared inflammatory expression pattern was accompanied by different sourcemarker responses.

In monocytes, the strongest decreases among the displayed source markers were *S100A9* (*−*0.91) and *VCAN* (*−*0.53), whereas *FCN1* and *LYZ* changed less (*−*0.11 and *−*0.06, respectively). Scavenging macrophages showed positive changes in all four displayed source markers: *C1QA* (+0.35), *C1QB* (+0.47), *APOE* (+0.85) and *SELENOP* (+0.80), along-side increased inflammatory markers. Resident macrophages showed a mixed response, with increased *F13A1* (+0.58) and *MRC1* (+0.35), but decreased *LYVE1* (*−*0.20) and *CFD* (*−*0.80).

Broader state-associated signatures supported inflammatory enrichment across all three source populations, with positive inflammatory-signature changes in every eligible library (Supplementary Figure 11). Source-signature responses differed: the Scav signature in-creased along inflammatory association (median library mean-gene log_2_FC +0.13), whereas the Res signature decreased (*−*0.24), despite increases in individual resident markers. As described above, the broader monocyte signature also increased despite attenuation of selected S100-family and other source markers. These comparisons show that shared inflammatory enrichment accompanies distinct changes within source-associated programmes.

The Scav pattern indicates maintenance and reinforcement of selected source-associated features alongside increased inflammatory expression. The mixed Res response and selective monocyte-marker decreases show that inflammatory association is compatible with several forms of source-identity remodelling. This interpretation is consistent with the coexistence of lipid-associated and inflammatory features reported for PLIN2^hi^/TREM1^hi^ macrophages in human plaques by Dib et al. [48]. Their TLR-dependent findings provide related bio-logical context, without establishing that their population corresponds to our Scav–Inflam relationship.

#### 3.4.2 CITE-seq corroborates coexisting source-associated and activation-associated protein features

We next examined whether the Scav expression pattern was accompanied by corresponding protein changes, using paired RNA and antibody measurements from the Bashore cohort [25]. Three libraries contained sufficient Q1 and Q4 cells within the Lipid-Scavenging (C1q^+^/HLA-DR^+^) subtype for matched comparisons (Fig. 4C). CD163, CD206 and MERTK increased in every eligible library, with median Q4 minus Q1 CLR differences of +0.334, +0.223 and +0.201, respectively. HLA-DR and CD38 also increased in all three libraries (median differences +0.423 and +0.179). The protein changes were selective: CD169 and CD64 varied between libraries, while CD86 and CD80 decreased consistently (median differences *−*0.185 and *−*0.130). Lineage markers and comparison antibodies generally showed smaller median changes, although some varied between libraries.

To determine whether elevated source-associated and activation-associated protein features occurred within the same cells, we performed a descriptive coexpression analysis using the mean CLR abundance of CD163/CD206/MERTK and HLA-DR/CD38, respectively. Each library’s Q1 medians defined the two reference thresholds. The fraction of cells above both thresholds increased from 26.3% to 61.7%, from 30.3% to 49.6%, and from 23.8% to 56.0% between Q1 and Q4 in the three libraries. These measurements corroborate joint elevation of selected protein features in individual antibody-tagged cells. They extend the RNA observations to a second modality within one cohort, while remaining cross-sectional measurements of phenotype.

#### 3.4.3 Supporting analyses of inflammatory association

The held-out-gene analysis also supported the Scav–Inflam association (Supplementary Figure 6). Among 26 eligible libraries, 760 genes excluded from the embedding showed Q4-versus-Q1 changes at global *q <* 0.05. Higher target association also accompanied smaller distance to the inflammatory reference in held-out-gene expression space, with adjusted sta-tistical support across all three gene panels in the analysis of 17 eligible libraries. Res–Inflam did not reach the adjusted threshold in either analysis, with coverage limited to four analysis units from five libraries for expression changes and three units for target resemblance. These controls corroborate the Scav–Inflam association beyond the embedding features, while support for the Res association remains more limited.

To assess whether greater inflammatory association coincided with higher residual doublet scores, we compared saved SOLO scores across the corrected OT quartiles in the current atlas (Supplementary Figure 7). Cell-ID-matched, within-library/subtype comparisons included 26 Scav, five Res and 18 Mono libraries, requiring at least 20 Q1 and 20 Q4 cells per eligible subtype/library combination. Grouping known paired preparations yielded 26, four and 17 analysis units, respectively, for statistical inference. This analysis evaluates residual score enrichment among retained cells; it does not establish the absence of doublets.

Mean SOLO score changes across analysis units were modest for Scav–Inflam (+0.011) and Mono–Inflam (+0.024), with neither comparison significant after multiple-testing correction (*q* = 0.471 and *q* = 0.444, respectively). For Res–Inflam, scores increased in all four analysis units (mean change +0.044; exact *p* = 0.125, adjusted *q* = 0.375; Supplementary Figure 7). This consistent increase warrants caution when interpreting the Resident comparison.

#### 3.4.4 Inflammatory target association is linked to symptomatic plaque status

We then asked whether the distribution of inflammatory association within source populations differed with clinical presentation. In the 18 Bashore patients with mapped clinical status (11 symptomatic and 7 asymptomatic), we calculated each patient’s fraction of monocytes or Scavenging macrophages in the upper decile of the corresponding inflammatory-association distribution (Fig. 4E; Supplementary Table 8). Thresholds were defined separately for the two source populations using cells from the mapped Bashore patients.

Symptomatic patients had a higher fraction of monocytes with high inflammatory association, with a symptomatic-minus-asymptomatic median difference of 13.0 percentage points (one-sided patient-label permutation *p* = 0.0017). The corresponding Scav comparison showed an estimated difference of 8.3 percentage points (*p* = 0.0632). These exploratory, unadjusted tests do not establish a difference between the two route effects, but identify an association between monocyte inflammatory programming and symptomatic status in this cohort. They assess the distribution of cell states at sampling, rather than prospective event risk or the rate of cell-state conversion.

Taken together, this axis links inflammatory-associated expression to distinct patterns of source-identity remodelling, corroborates selected Scav features at the protein level, and identifies clinical relevance for variation in monocyte inflammatory association. The strongest OT connection and the clearest symptom association involve different source populations, reflecting the distinct population-level and patient-level questions addressed by the method. Having compared several sources associated with the same inflammatory destination, we next examine whether Scav source features change differently along inflammatory and Fibroassociated gradients.

### 3.5 Biological Axis C: Source-identity remodelling along Fibrotic association

The third axis examines how association with Fibrotic/Hypoxic macrophages relates to changes in source identity (Fig. 5A). Scav–Fibro showed the strongest OT connectivity with the Fibro state (0.212), followed by Foam–Fibro (0.134), Mono–Fibro (0.103), Res–Fibro (0.070) and Inflam–Fibro (0.049; Supplementary Table 4). These relationships identify transcriptionally related populations for comparison without estimating their contributions to the Fibro compartment.

The primary UniTVelo analysis oriented the Scav, Mono, Foam and Inflam pairs towards Fibro (Fig. 5E; Supplementary Table 5). The expression analyses below characterize source cells across Fibro association using the same framework applied to the preceding axes.

#### 3.5.1 Fibro-associated expression accompanies selective changes in source programmes

Along Scav–Fibro association, the four displayed destination markers increased: *SPP1* (+2.37), *FN1* (+1.11), *ANXA2* (+0.84) and *LGALS1* (+0.88 median library log_2_FC; Fig. 5B; Supplementary Table 6). These changes accompanied lower *C1QA*, *C1QB* and *SELENOP* expression (*−*0.10, *−*0.15 and *−*0.50, respectively), while *APOE* increased (+0.50). Thus, Fibro association involved selective attenuation of scavenger-associated features alongside retained or reinforced expression of others.

Inflammatory macrophages also showed increased expression of all four displayed Fibroassociated markers, including *SPP1* (+1.42) and *FN1* (+1.78). Source-marker changes were selective: *CCL3*, *CCL4* and *CXCL8* decreased, whereas *IL1B* showed little median change (+0.06; Fig. 5B). Across the wider set of source populations, the broader Fibro signature increased in every eligible library, while source-signature responses varied (Supplementary Figure 11). These results extend the destination-associated expression pattern beyond individual markers without implying uniform loss of the source programme.

The Scav–Fibro association was also supported by the held-out-gene analyses. Among 21 eligible libraries, 806 genes excluded from the embedding showed Q4-versus-Q1 differences at global *q <* 0.05. Higher Fibro association accompanied greater resemblance to the Fibro reference, with adjusted statistical support across all three held-out gene panels in the analysis of 15 eligible libraries (Supplementary Figure 6). In a separate comparison using a shared 244-gene panel, Q4 cells were closer than Q1 cells to both Fibro and Foam references, with a larger reduction towards Fibro in eight of nine matched libraries. This supports preferential Fibro resemblance while recognizing shared expression features between the two destinations.

#### 3.5.2 Matched RNA and protein comparisons distinguish inflammatory and Fibro-associated remodelling

The contrasting Scav marker responses along inflammatory and Fibro association prompted a direct comparison within the same source population. We restricted this analysis to 19 libraries from four cohorts, comprising 25 library–subtype combinations eligible for both gradients. The broader Scav signature increased along inflammatory association but decreased along Fibro association (median library mean-gene log_2_FC +0.14 and *−*0.17, respectively; Fig. 5D). The change in the Scav signature was lower along Fibro than inflammatory association in all 19 matched libraries. In 14 of these libraries, the signature increased along inflammatory association and decreased along Fibro association. This comparison shows that the destination-associated difference persists after matching library and subtype composition. We then examined whether the corresponding contrast was present at the protein level.

Two Bashore libraries contained sufficient Q1 and Q4 cells for both gradients within the LipidScavenging subtype (Fig. 5C). CD163 and CD206 increased along inflammatory association and decreased along Fibro association in both libraries. CD169 and MERTK changes were also lower along Fibro association in both libraries, although their signs were less consistent. CD64 increased along Fibro association, indicating that the contrast did not extend uniformly across all scavenger-associated proteins. These measurements corroborate selected sourceprotein changes across destinations; the measured antibody panel did not directly test the Fibro-associated destination markers.

Taken together, the matched RNA and protein comparisons show that the same expressiondefined source population can display different patterns of source-feature remodelling depending on its destination association. Scavenger-associated features were reinforced alongside inflammatory expression, whereas several were attenuated alongside Fibro-associated expression. Together with the monocyte comparisons, these findings place selective retention, reinforcement and attenuation of source programmes within a common description of macrophage plasticity. They describe cross-sectional molecular patterns rather than demonstrated conversion or resolution pathways.

### 3.6 A spectrum of macrophage programme remodelling

Across the three biological axes, OT target-association gradients revealed different combinations of source-feature attenuation, retention and reinforcement alongside destinationassociated expression (Figs. 3B,C, 4B,C and 5B–D). Broader state-associated signatures showed positive destination-programme changes in every eligible library across the 11 analysed relationships, whereas source-programme changes varied in direction and magnitude (Supplementary Figure 11). Along Mono–Inflam association, for example, S100-family and *VCAN* expression decreased while the broader monocyte signature increased. The matched Scav comparison further showed that source-programme responses differed between inflammatory and Fibro association. Thus, the observed pattern depends on the particular source– destination relationship, and cannot be assigned solely from the source-state label.

To examine the regulatory associations accompanying these expression patterns, we inferred transcription factor activities using decoupleR [43] with the CollecTRI prior network [44] across all 11 relationships. Q4-versus-Q1 activity differences were calculated within eligible library/subtype combinations and compared across libraries, with multiple-testing correction across regulators and routes (Supplementary Table 7; Supplementary Methods). These estimates summarize expression patterns among regulator target genes.

Both Mono–Inflam and Scav–Inflam showed increased inferred activity of *RELA*, *NFKB1*, *REL* and *RELB*, together with *CIITA* and the RFX regulators *RFX5*, *RFXAP* and *RFXANK* (all global adjusted *q <* 0.05). The corresponding effects were also positive along Res–Inflam, although they did not reach the adjusted threshold in its five eligible libraries. Increased CI-ITA and RFX activities were additionally observed along Mono–Scav association, indicating that these regulatory features were shared across several macrophage relationships rather than restricted to inflammatory association.

Fibro-associated regulatory changes were also dependent on the source population. Scav– Fibro showed lower inferred activity of NF-*κ*B family members alongside increased *MYC* and *HOXA7* activity. Increased MYC and HOXA7 activities were also supported along Mono– Fibro and Mono–Foam association (global adjusted *q <* 0.05). However, MYC activity also increased along Scav–Inflam, and the decreases in NF-*κ*B activities along Mono–Foam did not reach the adjusted threshold. These findings identify shared and contrasting regulatory associations across routes, without supporting a universal opposition between inflammatory and Fibro-associated programmes or establishing that foam-cell formation requires inflammatory silencing.

In a targeted sensitivity analysis combining paired fresh and frozen preparations from the Jaiswal cohort, all broad source- and destination-signature medians retained their sign, and the principal regulatory associations described above retained adjusted statistical support.

Together, the marker, broader-signature and inferred regulatory analyses characterize macrophage plasticity through selective remodelling of overlapping programmes. Selected RNA patterns were corroborated by protein measurements, including the destination-associated contrast within the Scav population. The associated regulator profiles identify candidates for functional investigation of these selective changes in source and destination programmes.

### 3.7 Extension of macrophage programme analysis to healthy and cirrhotic human liver

To examine whether the analytical framework could be extended beyond atherosclerotic plaques, we applied the OT connectivity and target-association analyses to an independent scRNA-seq dataset of healthy and cirrhotic human liver from Ramachandran et al. [26] (GSE136103). The original study included five healthy and five cirrhotic liver donors. Our annotated mononuclear phagocyte compartment comprised 8,536 cells distributed across four populations: tissue monocytes, Kupffer cells, C1q^+^/MHC-II macrophages and scar-associated macrophages (SAM; Fig. 6A). This setting allowed comparison of distinct hepatic source populations associated with a common disease-associated macrophage state.

**Figure 6:**
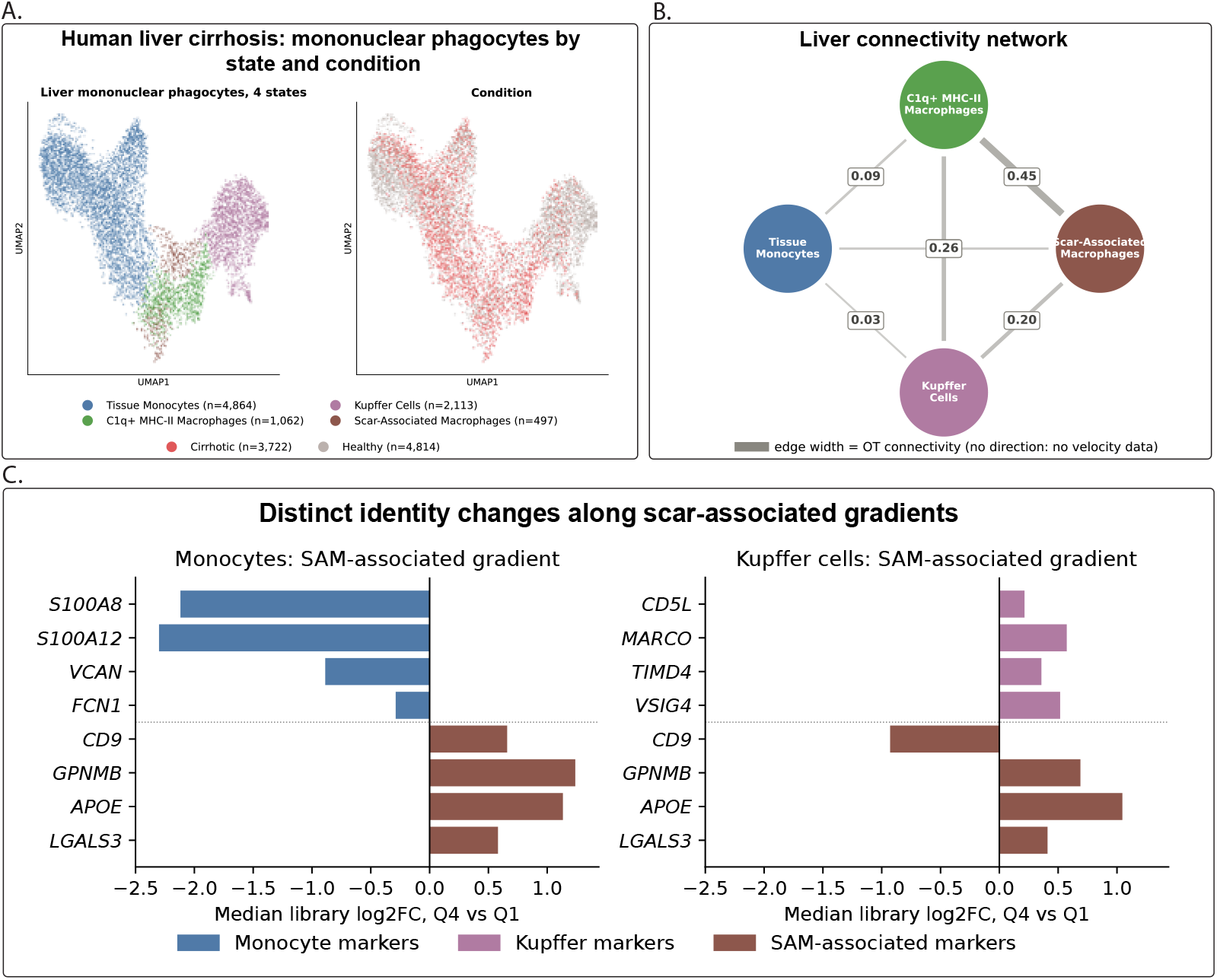
Macrophage state relationships and source-associated programmes in human liver cirrhosis. **(A)** UMAP embeddings of 8,536 liver mononuclear phagocytes, coloured by annotated state and disease condition. The analysis comprised Tissue Monocytes (*n* = 4,864), Kupffer Cells (*n* = 2,113), C1q^+^/MHC-II Macrophages (*n* = 1,062) and Scar-Associated Macrophages (SAM; *n* = 497), including 4,814 cells from healthy and 3,722 from cirrhotic liver samples. **(B)** OT-based connectivity among the four liver populations. Edge thickness and labels represent population-level connectivity. The network is undirected because RNA velocity was not evaluated in this dataset. **(C)** Source-marker and SAM-associated marker changes along Tissue Monocyte–SAM and Kupffer Cell–SAM target-association gradients. Bars show median library log_2_ fold changes between Q4 and Q1, calculated from linearised normalised expression with a pseudocount of 0.1. Comparisons required at least 20 source cells in each quartile within an eligible library. Source markers are blue for tissue monocytes and purple for Kupffer cells; SAM-associated markers are brown. Coverage was 13 monocyte libraries (seven healthy and six cirrhotic) and five Kupffer-cell libraries (four healthy and one cirrhotic).

Within the liver analysis, SAM showed their strongest OT connection with C1q^+^/MHC-II macrophages (connectivity = 0.454), followed by Kupffer cells (0.200) and tissue monocytes (0.083; Fig. 6B). Kupffer cells were also connected to the C1q^+^/MHC-II population (0.256). These relationships describe the relative transcriptional proximity of the hepatic populations. They provide a basis for examining SAM-associated expression within different source compartments without specifying a temporal sequence of cell-state conversion.

We next compared source-associated and SAM-associated expression along the tissuemonocyte and Kupffer-cell target-association gradients (Fig. 6C). Monocyte markers comprised *S100A8*, *S100A12*, *VCAN* and *FCN1*, while Kupffer-cell markers comprised *CD5L*, *MARCO*, *TIMD4* and *VSIG4*. The same four SAM-associated genes, *CD9*, *GPNMB*, *APOE* and *LGALS3*, were examined in both source populations. As in the plaque analyses, expression differences were calculated between highand low-association source cells within eligible libraries and summarized across libraries.

The comparisons included 13 eligible monocyte libraries (seven healthy and six cirrhotic) and five Kupffer-cell libraries (four healthy and one cirrhotic). Along monocyte–SAM association, all four selected monocyte markers had negative median changes, while all four SAM-associated markers had positive median changes. Kupffer cells instead showed positive median changes in all four selected source markers. Among the SAM-associated markers, *GPNMB* and *APOE* increased in all five Kupffer-cell comparisons and *LGALS3* in four. In contrast, *CD9* decreased in four of five libraries, accompanied by a lower fraction of CD9-expressing cells in the same four comparisons (median library log_2_ fold change = *−*0.93; Fig. 6C). Annotated SAM showed substantially higher CD9 expression than Kupffer cells in matched libraries, including all four references containing at least 20 SAM cells. The Kupffer-cell gradient therefore captures increased expression of selected SAM-associated features alongside maintained source markers, rather than uniform convergence towards the complete SAM marker profile. The predominantly healthy Kupffer-cell coverage limits conclusions specific to cirrhosis.

The liver analysis extends the comparative framework to a second tissue and disease context, allowing source-associated and disease-associated expression to be examined within distinct hepatic macrophage populations. As in plaques, destination-associated expression accompanied distinct source-marker responses: attenuation of selected monocyte features and reinforcement of Kupffer-cell markers alongside partial SAM-associated expression. Any correspondence between the two tissues should therefore be interpreted as recurrence of expression patterns, rather than evidence that a shared developmental origin determines a universal mechanism of macrophage plasticity.

## 4 Discussion

In this study, we developed a computational method combining pairwise optimal transport and target-association gradient analysis to characterize relationships among macrophage states in human atherosclerotic plaques. Across seven cohorts, the resulting network organized these relationships around three biological axes: monocyte diversification, inflammatory programming, and fibrotic-associated remodeling. By examining source and destination programs together, we identified distinct patterns of selective attenuation, retention, and reinforcement of source-state features. Selected patterns were corroborated at the protein level, while patient-level analysis linked inflammatory-associated monocyte states to symptomatic disease. Together, these findings provide a comparative framework for investigating macrophage plasticity and prioritizing candidate pathogenic routes between established cell states. Its biological value lies in connecting the populations present in a plaque to the molecular changes through which they may participate in disease.

Two features make this method particularly useful for investigating these relationships. First, population-level pairwise transport compares cell states without requiring a predefined branching topology or a terminal population. Second, target-association gradients characterize variation within each source population in relation to a specified destination, allowing source and destination programs to be examined along the same axis. This combination of population-level and within-state analyses is particularly relevant in atherosclerosis, where macrophage heterogeneity is increasingly well resolved across single-cell studies [12, 16], but the relationships among these populations remain less well understood. The integrated atlas of Traeuble et al. [16] illustrates that disease-associated differences involve both changes in the abundance of selected macrophage subsets and changes in their transcriptional programs. Our approach addresses the complementary question of how such programs vary across candidate connections between states. Comparing all 15 pairwise relationships within a common framework, followed by source–destination program analysis, provides a systematic basis for selecting populations and molecular features for experimental investigation.

An important consideration is that OT divergence measures distributional distance rather than cell fate. Transcriptionally proximal populations may share environmental responses or developmental history without undergoing interconversion. We therefore interpret the network as a map of candidate relationships whose biological relevance must be assessed through additional evidence. Within-library and subtype-matched comparisons identified destination-associated expression changes alongside distinct source-program responses, while analyses of genes excluded from the embedding supported aspects of these relationships beyond the features used to construct it. Broader signatures also allowed us to distinguish changes in individual markers from changes extending across a source-associated program. These analyses strengthen the characterization of the candidate routes, although they remain based on correlated measurements from the same transcriptomes.

A central finding is that macrophage plasticity is not well described by a binary switch between complete cell identities. This is consistent with experimental descriptions of macrophage activation as a spectrum of transcriptional programs [10]. Our analysis extends this perspective by explicitly comparing the behavior of source-associated features across different destinations. Maintenance, reinforcement, and selective attenuation of source-associated features can coexist within the same comparison, alongside increased destination-associated expression. For example, monocytes with stronger inflammatory association showed lower expression of classical markers, including S100-family genes and *VCAN*, despite an increase in the broader monocyte-associated signature. Conversely, monocyte associations with the scavenging and foam states involved decreases in the broader source signature. Thus, loss of a familiar marker does not necessarily imply loss of the complete source program, and the relevant unit of plasticity may be a set of selectively changing features within an otherwise recognizable cellular state.

The comparison between inflammatory and Fibrotic associations within the same scavenging population provides a particularly informative example. The scavenging signature increased along the inflammatory association but decreased along the Fibrotic association, with a lower Fibrotic-associated change in all 19 matched libraries. This contrast was accompanied by increased inflammatory or Fibrotic-associated expression, respectively. Because the source population and contributing libraries were matched, the comparison supports destination-associated differences in source-program remodeling beyond a simple contrast between unrelated populations. It also shows why the extent of source-identity maintenance cannot be assigned as a fixed property of monocytes or tissue-adapted macrophages. A population retaining recognizable scavenging features can exhibit different molecular relationships with different disease-associated states.

CITE-seq provided a complementary test of these patterns at the protein level. Along the scavenging-to-inflammatory association, CD163, CD206, and MERTK increased together with HLA-DR and CD38 across the three eligible libraries. The increased fraction of cells scoring above the reference thresholds for both protein groups further supported their coexistence within individual cells. The response was selective, with CD80 and CD86 decreasing, emphasizing that inflammatory-associated programming does not entail uniform increases in all activation markers. Across the two libraries eligible for the matched destination comparison, CD163 and CD206 increased along the inflammatory association and decreased along the Fibrotic association. These observations corroborate selected source-feature changes in a second molecular modality. They support the biological relevance of the RNA patterns while defining their scope: the available protein measurements characterize source and activationassociated features, rather than directly establishing acquisition of a fibrogenic function.

One possible explanation for these selective responses is that macrophage identity and stimulus-responsive programs are regulated through partially separable mechanisms. Experimental studies have shown that the local microenvironment shapes macrophage enhancer landscapes and that factors such as ZEB2 help maintain tissue-specific identity [49, 50]. Such organization could permit new functional programs to coexist with established features while allowing other components to change under different environmental conditions. Our observations are compatible with this model, but neither chromatin regulation nor developmental ancestry was measured here. In particular, the expression-defined Resident and Scavenging populations should not be interpreted as lineage-traced populations. The differences observed across source–destination combinations instead motivate experiments testing how local signals interact with the existing transcriptional state to determine which macrophage features are maintained or remodeled.

Transcription-factor activity inference provides candidate regulatory programs for these experiments. Monocyte and scavenging associations with the inflammatory state were accompanied by increased inferred NF-*κ*B and RFX–CIITA activity, linking inflammatory and antigen-presentation programs across distinct source populations. The scavenging-to-Fibrotic association instead showed lower inferred NF-*κ*B activity together with increased MYC and HOXA7 activity. However, these programs were overlapping rather than mutually exclusive: MYC-associated activity also increased along the scavenging-to-inflammatory association, and some routes had insufficient library-level support to establish comparable regulatory changes. These results suggest that common regulatory components may be combined differently across candidate routes. Their value is to identify specific hypotheses for perturbation, including whether interfering with a destination-associated program changes the response while preserving selected source features.

The topology of the network helps prioritize where such experiments may be most informative. The scavenging–inflammatory relationship showed the strongest OT connectivity and remained the closest pair in every leave-one-cohort-out analysis. Together with the RNA and protein findings, this makes the relationship a strong candidate for investigating how inflammatory features emerge within macrophages that retain lipid-handling and scavenging-associated characteristics. This interpretation is biologically consistent with the TLR2-dependent transition of lipid-associated macrophages described by Dib et al. [48], although their experimentally defined populations cannot be assumed to correspond exactly to our meta-clusters. The strength of an OT connection also does not quantify the number of cells taking that route. Its role is to prioritize a relationship for investigation, alongside the molecular programs and clinical associations that may make it relevant to disease.

The patient-level analysis adds a distinct clinical perspective. In the Bashore cohort, symptomatic patients had a higher fraction of monocytes with strong inflammatory association, with a median difference of 13 percentage points. The corresponding scavenging comparison was less statistically resolved. Thus, the closest population-level relationship and the clearest symptom-associated cellular pattern addressed different aspects of the inflammatory axis. One identifies transcriptionally related states across the atlas; the other identifies variation within a source population associated with clinical presentation. This distinction is useful when prioritizing candidate pathogenic routes, because transcriptional proximity alone need not determine clinical relevance. The clinical findings remain exploratory, given the 18patient sample, the examination of alternative endpoints, and the absence of an independent clinical replication cohort.

Application to the human liver dataset extended the same analytical strategy to a separate tissue context containing monocytes, Kupffer cells, and scar-associated macrophages [26]. Along association with scar-associated macrophages, selected monocyte markers declined as SAM-associated markers increased, whereas Kupffer-cell markers increased alongside only part of the SAM-associated program. The decrease in CD9 further showed that the Kupffercell gradient did not represent uniform convergence towards the full SAM marker profile. These contrasting responses illustrate how the method can resolve selective source-feature changes in a separate tissue context, although the predominantly healthy Kupffer-cell coverage limits conclusions specific to cirrhosis. Establishing which relationships or regulatory mechanisms are shared between plaque and liver macrophages will require direct comparison across appropriately matched datasets and experimental systems.

These findings may also inform how macrophage-directed interventions are conceived. Trials such as CANTOS established that targeting inflammation can reduce recurrent cardiovascular events [4]. A complementary biological question is whether particular routes into disease-associated macrophage states can be perturbed while preserving other functions of the participating populations. The coexistence of source and destination features makes this question especially relevant: a macrophage displaying scavenging-associated markers may also express an inflammatory program, so its functional contribution cannot be inferred from the source label alone. The experimentally characterized TLR2-dependent switch [48] provides one precedent for investigating such selectivity. Our comparative analysis identifies additional source–destination combinations and regulatory candidates for testing whether disease-associated programming can be modified while maintaining functions such as lipid handling and clearance of dying cells. Demonstrating that separation will require functional measurements alongside changes in state-associated markers.

Several limitations should be considered. First, cross-sectional scRNA-seq data cannot directly establish temporal lineage relationships, transition rates, or the functional consequences of a candidate route. Expression of Fibrotic-associated genes, for example, does not by itself establish a fibrogenic effect. Second, protein corroboration was restricted to one cohort and a small number of eligible libraries, and the RNA, protein, and clinical analyses partly reused the same biological material. Third, the clinical association requires confirmation in larger, prospectively characterized cohorts. Fourth, the contributing datasets differed in vascular territory and sample preparation, including fresh and cryopreserved specimens. Although the closest state pair was stable across cohort omissions, other relationships showed greater sensitivity, particularly to omission of the Bashore and Jaiswal cohorts; generalizability across vascular territories remains to be established. Direct evaluation of the prioritized routes could combine perturbation of candidate regulators with RNA, protein, and functional readouts. Time-stamped approaches such as Zman-seq [51] and inducible fate mapping in atherosclerosis models could additionally test the temporal relationships that remain unresolved in human cross-sectional tissue.

## 5 Conclusions

These results connect the description of macrophage heterogeneity to experimentally testable questions about disease-associated state change. The method brings together populationlevel relationships, selective source-program remodeling, destination-associated expression, and clinical context to prioritize candidate pathogenic routes. This approach is particularly relevant to chronic inflammatory disease, where recognizable macrophage populations may contain functionally different cellular states. Characterizing those relationships provides a basis for investigating how pathological programs arise and, ultimately, whether they can be altered while preserving beneficial macrophage functions.

## List of abbreviations

ADT, antibody-derived tag; ASW, average silhouette width; BBKNN, batch-balanced knearest neighbors; CITE-seq, cellular indexing of transcriptomes and epitopes by sequencing; CLR, centered log ratio; HVG, highly variable gene; iLISI, integration local inverse Simpson index; LOCO, leave-one-cohort-out; NF-*κ*B, nuclear factor kappa B; OT, optimal transport; PCR, principal component regression; SAM, scar-associated macrophage; scRNA-seq, single-cell RNA sequencing; scVI, single-cell variational inference; TF, transcription factor; UMAP, uniform manifold approximation and projection; UMI, unique molecular identifier.

Meta-cluster abbreviations: Mono, Monocytes; Scav, Scavenging/C1q^+^ Macrophages; Res, Resident/Quiescent Macrophages; Inflam, Inflammatory Macrophages; Foam, LipidStressed/Foam Cells; Fibro, Fibrotic/Hypoxic Macrophages.

## Declarations

### Ethics approval and consent to participate

This study reanalysed publicly available, de-identified human scRNA-seq and CITE-seq datasets. No new participants were recruited and no new human specimens were collected. In accordance with institutional requirements, no additional ethics approval was required for this secondary analysis. Ethics approval and participant-consent procedures for the original studies are described in the corresponding publications.

## Consent for publication

Not applicable. The manuscript contains no identifiable individual participant information.

## Availability of data and materials

The scRNA-seq and CITE-seq datasets analyzed in this study are publicly available through the repositories associated with their original publications. Library mappings, analysis coverage, and supporting numerical outputs are provided in Additional file 1 and the associated source data.

Analysis code is available at https://github.com/sergiovmdo/macrophage-ot-atlas and archived at Zenodo (https://doi.org/10.5281/zenodo.21940799).

## Competing interests

The authors declare no competing interests.

## Funding

S.V.M.O. was supported by the European Innovation Council (EIC) program (MIRACLE, 101115381). T.Ö. was supported by the Research Council of Finland (Grant No. 352968).

M.U.K. was supported by the Sigrid Juselius Foundation, the Finnish Foundation for Cardiovascular Research, the European Innovation Council (EIC) program (MIRACLE, 101115381), and the European Research Council of the European Union (SECRET, 101125115). Views and opinions expressed are, however, those of the author(s) only and do not necessarily reflect those of the European Union or the European Research Council. Neither the European Union nor the granting authority can be held responsible for them. S.W. was supported by the China Scholarship Council (CSC) Fellowship Programme (No. 202306160023).

## Authors’ contributions

S.V.M.O., M.U.K., I.Z. and M.R.M. conceived and designed the study. S.V.M.O. developed the computational framework, implemented the analysis pipeline, performed all optimal transport, RNA velocity and target-association gradient analyses, produced the figures, and wrote the manuscript. A.A.T. and J.F.C.M. contributed to data integration and preprocessing. P.K. contributed to interpretation of sc-RNA-Seq results. T.Ö. contributed biological expertise on plaque macrophage phenotypes. S.W. contributed to the clinical interpretation of the results. S.F. contributed to the biological interpretation of the transition landscape and to revision of the manuscript. I.Z. and M.R.M. supervised the study. M.R.M. acquired funding and contributed to manuscript writing. All authors read, revised and approved the final manuscript.

## Acknowledgements

The authors acknowledge the use of Codex (OpenAI; Astra) and Claude (Anthropic; Opus 4.8) for assistance with manuscript and code preparation, as described in the Methods.

We thank Michal Mokry (Central Diagnostics Laboratory, University Medical Center Utrecht) for valuable discussions and contributions to the clinical interpretation of the results. We also thank the investigators of the seven original cohorts for making their data publicly available, and acknowledge the UBELIX high-performance computing cluster at the University of Bern, on which all analyses were run.

## Additional files

**Additional file 1 (PDF): Supplementary Information.** Supplementary Methods; Supplementary Tables 1–9; Supplementary Figures 1–11; and the supplementary note on sequencingdepth correction of OT target-association scores.

## Notes

### Competing Interest Statement

The authors have declared no competing interest.

### Summary of Updates

Inclusion of new studies including the expansion to a different disease and some clinical experiments that have been conducted using the framework.

