## Supplementary text & materials for "Cross-cohort optimal transport maps macrophage plasticity and routes to inflammation and fibrosis in human atherosclerosis"

### Additional file 1: Supplementary Information

#### Supplementary Methods

##### Dataset scope, preprocessing and quality control

The plaque analyses use the rebuilt atlas of 34,533 monocytes and macrophages from 50 libraries across seven cohorts (Alsaigh, Bashore, Fernandez, Jaiswal, Pan, Pauli and Wirka). Sequencing runs belonging to the same library were processed together, so that a cell sequenced across multiple lanes was represented once. A sequencing run, library and patient are distinct units; patient-level analyses used the explicit mapping in Supplementary Table 8.

Reads were processed using Cell Ranger v10.1.0 and the GRCh38-2020-A reference. Ambient RNA correction was performed per library with CellBender [1], using the unfiltered matrix, the Cell Ranger expected-cell estimate, a false-positive rate of 0.01, 20,000 included droplets and 150 epochs. All 50 libraries represented in the final atlas had CellBender-corrected RNA matrices. Spliced, unspliced and ambiguous counts were generated separately with velocyto [2] and matched by library, barcode and gene identifier; CellBender correction of the RNA matrix was not applied to these splicing layers.

Initial filtering retained cells with at least 200 detected genes and less than 25% mitochondrial counts. SOLO [3] was trained separately within each library using scvi-tools [4]. Cells were ranked by the saved SOLO doublet score, and the highest-scoring $\mathrm{round}(Nf)$ cells were removed, where $N$ is the number of prefiltered cells and

$$f = \min\left(0.008\,\frac{N}{1000},\, 0.20\right).$$

This procedure uses a library-specific expected doublet fraction rather than a universal score threshold. Both soft scores and hard classifications were retained in the metadata, but the hard calls were not the final filtering rule.

Libraries with fewer than 1,000 input cells were excluded. Within each retained library, cells were filtered for low log-transformed counts or detected genes (more than five median absolute deviations below the respective median), high mitochondrial percentage (more than five median absolute deviations above its median), or mitochondrial percentage of at least 15%. A post-filter median below 500 detected genes triggered whole-library exclusion. Removal flags can overlap and were not summed as independent exclusions.

Supplementary Table 9 reports the executed settings and cell accounting: 242,433 cells in 62 QC-input libraries, 201,858 all-lineage cells retained across 50 libraries, and 34,533 cells in the final monocyte/macrophage atlas. Eleven libraries failed the minimum input-size criterion and one failed the post-filter median-gene criterion. The difference between all-lineage QC retention and final atlas size includes lineage selection and subsequent annotation review, and is not reported as additional QC loss. Supplementary Figure 1 describes the retained atlas, not the discarded cells.

##### Integration, subtype annotation and meta-cluster consolidation

CellBender-corrected RNA counts were normalized to 10,000 counts per cell and transformed using $\log_e(1+x)$. Coarse lineage annotation identified the myeloid compartment. Cohort-aware highly variable gene selection used the Seurat implementation in Scanpy [5], requesting 5,000 genes. Scanorama [6] integration used exact neighbor search, 100 dimensions, 50 neighbors, $\sigma = 15$ and $\alpha = 0.10$. The integrated representation was retained separately from the log-normalized expression matrix. A 50-neighbor graph supported UMAP visualization and Leiden clustering [7].

Cluster annotation combined canonical markers, differential-expression review and cohort composition.

Previously reviewed annotations were propagated through shared cells and cluster-level agreement, followed by marker inspection and additional subclustering. The final stored assignments are based on resolution-0.8 clusters and their subclusters. The resolution sweep in Supplementary Figure 2 is a diagnostic on the current graph; it is not a clustering tree or evidence that all final annotations arose from a single resolution-0.5 partition.

Annotation was anchored in established plaque, blood-monocyte and resident-macrophage marker panels [8–12]. This marker-guided approach allowed reviewed functional annotations to be retained across the rebuilt datasets without treating a transferred label as definitive. Supplementary Table 3 gives the stored cluster identifiers, final labels, cell counts and marker-expression summaries. Supplementary Figure 8 displays the corresponding fine-subtype marker profiles.

Nine nonempty annotated subtypes were consolidated into six meta-clusters:

- **Monocytes (Mono; $n = 9{,}961$):** Classical Monocytes ($n = 5{,}128$; *S100A8*, *S100A9*, *FCN1*); Intermediate/Non-Classical Monocytes ($n = 2{,}796$; *LST1*, *S100A4*, *FCGR3A*); and Transitional Monocytes ($n = 2{,}037$; *VCAN*, *ZEB2*).
- **Scavenging/C1q$^+$ macrophages (Scav; $n = 13{,}499$):** Homeostatic Lipid-Regulating Macrophages ($n = 4{,}614$; *APOE*, *SELENOP*) and Lipid-Scavenging Macrophages ($n = 8{,}885$; *C1QA*, *C1QB*, *HLA-DRA*, *MSR1*).
- **Resident/Quiescent macrophages (Res; $n = 1{,}409$):** the retained Resident Perivascular population, characterized by *LYVE1*, *FOLR2* and *F13A1*.
- **Inflammatory macrophages (Inflam; $n = 2{,}309$):** the *CCL3*/*CCL4*/*IL1B*/*CXCL8*-associated population.
- **Lipid-Stressed/Foam cells (Foam; $n = 3{,}240$):** the *FABP5*/*CSTB*/*PLIN2*/*LIPA*-associated population.
- **Fibrotic/Hypoxic macrophages (Fibro; $n = 4{,}115$):** the *SPP1*/*FN1*-associated population, also characterized using *ANXA2* and *LGALS1*.

Meta-cluster names describe expression-defined states. In particular, the Resident label does not establish developmental origin, and the Fibrotic label is not a direct assay of matrix production. All plaque OT analyses use these six meta-clusters; annotated subtypes are additionally used to stratify source-cell expression comparisons.

##### Batch integration quality assessment

*Comparison with alternative integration methods.* Scanorama, Harmony [13], BBKNN [14] and scVI [15] were evaluated on the same 20,576 cells, spanning seven cohorts and all 50 libraries. Sampling capped cohort contributions at approximately 5,000 cells while retaining meta-cluster proportions. The subset contained 4,999 Alsaigh, 4,999 Bashore, 4,488 Jaiswal, 2,384 Wirka, 2,037 Pan, 1,350 Fernandez and 319 Pauli cells.

Batch silhouette width, graph connectivity and principal-component-regression (PCR) scores were calculated using scIB [16]. Batch silhouette width measures mixing within annotated cell types; graph connectivity measures connectivity within these groups. These two metrics therefore use the existing annotations. For methods producing embeddings, neighbor graphs used 15 neighbors; BBKNN retained its corrected graph. PCR compared cohort-associated variation in each benchmark object's expression matrix with its supplied embedding. The saved composite was the arithmetic mean of the three metrics.

BBKNN's saved PCR value of zero is a consequence of this graph-based comparison setup and is not interpreted as a comparable corrected-embedding result; its PCR and composite are omitted from the printed comparison but retained in the source data.

Batch silhouette scores ranged from 0.911 to 0.934 and graph connectivity from 0.985 to 0.998 (Supplementary Table 1). Harmony had the highest saved composite, followed by scVI and Scanorama. Scanorama was retained as the representation used throughout the comparative OT analysis, with RNA and splicing matrices stored separately for downstream analyses. The benchmark assesses the stated integration properties, not superiority for transition inference.

*Local cohort mixing and cohort representation.* The local inverse Simpson index (iLISI) was calculated using `harmonypy` with perplexity 30, separately within each meta-cluster in the full Scanorama embedding, without cell downsampling. Raw iLISI is the effective local number of cohorts; normalized values were $(\mathrm{iLISI}-1)/6$ for seven cohorts. State-wide cohort diversity was calculated separately as $1/\sum_c p_c^2$, where $p_c$ is the fraction of cells from cohort $c$. This composition statistic is not a local-mixing ceiling. Cohort enrichment was the percentage-point difference between a cohort's fraction within a state and its fraction in the entire atlas. UMAP spread was the unweighted mean pairwise distance between cohort centroids.

All six states contained cells from all seven cohorts, but local cohort concentration remained substantial (median iLISI, 1.015–1.448; Supplementary Table 2). These quantities describe local mixing and representation; differences can reflect technical effects, sampled biological heterogeneity or both. Library-balanced expression comparisons and cohort-omission analyses address complementary aspects of this heterogeneity.

##### Optimal transport divergence computation

Population-level OT divergence was computed for all 15 unordered meta-cluster pairs in the 100-dimensional Scanorama representation. The reference calculation sampled each state without replacement to a maximum of 5,000 cells (seed 42). Source and target marginals were uniform. Pairwise Euclidean distances defined the cost matrix $C$. Each cross- or self-cost matrix was divided by its own maximum before fitting with the Sinkhorn solver in Python Optimal Transport [17], using $\varepsilon = 0.01$ and at most 2,000 iterations.

After fitting, the transport-cost component was returned to the original distance scale:

$$W(A,B) = \langle T_{AB}, C_{AB}/\max(C_{AB})\rangle \max(C_{AB}) = \langle T_{AB}, C_{AB}\rangle.$$

The reported scalar excludes the entropy term used to fit the plan. Self-transport costs were evaluated by the same procedure, and the reported self-cost-corrected OT divergence was

$$S(A,B) = \max\left\{0,\, W(A,B) - \tfrac{1}{2}W(A,A) - \tfrac{1}{2}W(B,B)\right\}.$$

Diagonal entries were set to zero. Here "OT divergence" refers to this implemented transport-cost statistic, rather than an entropy-inclusive objective.

*Cell- and library-level resampling.* The reference run included 500 cell-bootstrap and 200 library-bootstrap iterations. Cell bootstraps sampled with replacement within states, to a maximum of 2,000 cells per state. Library bootstraps sampled library identifiers with replacement, concatenated their cells, and sampled up to 2,000 rows per state without replacement from that pool; resamples with fewer than 50 cells in any state were excluded. Cross- and self-costs were recomputed in each retained iteration. Means, standard deviations and 2.5th–97.5th percentile intervals summarize the resulting distributions.

Supplementary Table 4 and Supplementary Figure 3 emphasize library-resampling summaries. Their cell cap and sampling scheme differ from the reference estimate, so the percentile intervals describe the resampling distribution and are reported separately from the 5,000-cell reference.

*Label-permutation null model.* Meta-cluster labels were permuted across cells 50 times, preserving label counts and embedding coordinates. Divergences were recomputed with a maximum of 2,000 cells per state. Departure from random label mixing was summarized by

$$z_{AB} = \frac{S_{\text{observed}}(A,B) - \mu_{\text{null}}(A,B)}{\sigma_{\text{null}}(A,B)}.$$

The implemented retention criterion was $|z_{AB}| > 2$. All 15 pairs met this criterion, with $z$ ranging from 81.02 to 232.41. These standardized deviations quantify separation of labeled populations from a label-mixing null, not the probability of a biological transition; they are not empirical permutation $p$-values.

*Connectivity transformation.* For network visualization, reference divergences passing the label-null criterion were transformed using

$$K(A,B) = \exp\left[-\frac{S(A,B)^2}{2\sigma^2}\right], \qquad \sigma = \tfrac{1}{2}\,\text{median}\{S(A,B) > 0 :\ |z_{AB}| > 2\}.$$

The full-atlas bandwidth was $\sigma = 0.205758$. The diagonal and pairs not passing the criterion were assigned zero connectivity. Higher connectivity denotes greater relative transcriptional proximity in this representation; it is not a calibrated transition probability or a measure of cellular flux. Scav–Inflam had the smallest reference divergence (0.2739) and highest connectivity (0.4124).

#### Cohort-omission and regularization sensitivity

Leave-one-cohort-out (LOCO) calculations were performed for all seven cohorts by removing their cells from the existing integrated representation and repeating the OT analysis. Integration was not refitted. Reference divergence changes and ascending-rank changes were calculated relative to the full-cohort result; rank 1 denotes the closest pair. Run-specific connectivity bandwidths were not used to compare effect magnitudes across omissions.

Rank correlations with the full result ranged from 0.843 after omitting Bashore to 0.996 after omitting Fernandez or Pauli. The largest absolute divergence changes were 0.104 after Bashore omission and 0.079 after Jaiswal omission; maximum rank changes were seven and three positions, respectively. Scav–Inflam remained the closest pair in all seven runs. Supplementary Figure 4 displays both signed changes, while Supplementary Table 4 retains all 105 pair-by-omission estimates.

Regularization sensitivity used $\varepsilon \in \{0.005, 0.01, 0.025, 0.05, 0.1, 0.5\}$ with the same representation and reference sampling scheme. Spearman correlations with the selected $\varepsilon = 0.01$ ranking were 0.993, 1.000, 1.000, 0.954, 0.854 and 0.654, respectively (Supplementary Figure 9). These results describe stability over the evaluated settings; stronger regularization changed the ordering of several pairs.

#### OT target-association gradients

*Cell-level score.* A separate balanced transport calculation ranked source cells within each source–destination comparison. All available source and target cells were used, with the first 50 Scanorama dimensions, squared Euclidean costs normalized by their maximum, $\varepsilon = 0.01$, at most 2,000 iterations and convergence tolerance $10^{-9}$. Source marginals were $a_i = 1/n_A$. Target marginals were proportional to inverse local distance:

$$b_j = \frac{(\bar{d}_j + 10^{-10})^{-1}}{\sum_k (\bar{d}_k + 10^{-10})^{-1}},$$

where $\bar{d}_j$ is the mean Euclidean distance to the ten returned nearest neighbors within the target population, including the query cell itself. For source cell $i$, the score was

$$s_i = \frac{1}{a_i} \sum_j T_{ij} b_j = n_A \sum_j T_{ij} b_j.$$

It is therefore the expected target-density weight under the source cell's conditional coupling. It is not the row sum of a balanced plan, which is fixed by the source marginal, and does not directly measure elapsed time or fate probability.

*Count correction and quartiles.* Within each comparison, scores were regressed on total RNA counts using ordinary least squares with an intercept. Residuals defined the corrected scores; source cells were divided into four approximately equal-sized quartiles (Q1–Q4). Saved corrected scores and quartiles were reused across the analyses. No rank-based fallback was applied in the current analysis. Supplementary Figure 10 and the accompanying Note report Pearson and Spearman associations with counts before and after correction.

*Comparison inventory.* Eleven unique source–destination comparisons were analyzed: Mono with Scav, Inflam, Foam, Res and Fibro; Scav and Res with Inflam; and Scav, Res, Foam and Inflam with Fibro. Mono–Inflam is shared between the Monocyte and Inflammatory figure narratives, and Mono–Fibro between the Monocyte and Fibrotic narratives; these are not additional independent comparisons. An arrow in an expression comparison specifies the source population and the target used to construct the gradient. Temporal orientation is evaluated separately by the velocity analysis.

##### Gene-level contrasts, marker panels and broader signatures

Log-normalized expression values were returned to the linear normalized scale using $\mathrm{expm1}$ before calculating RNA means. Q1 and Q4 cells were compared within each library and annotated source subtype. A library–subtype stratum was eligible if it contained at least 20 cells in each endpoint quartile. For gene $g$, library $\ell$ and subtype $h$,

$$d_{g\ell h} = \log_2 \left( \frac{\bar{x}_{g,\ell h,Q4} + 0.1}{\bar{x}_{g,\ell h,Q1} + 0.1} \right).$$

Eligible subtype effects were averaged equally within each library, and the median library effect was displayed. This gives each library equal weight, irrespective of cell yield. Coverage, per-stratum and per-library effects are retained in the source data for Supplementary Table 6. Canonical panels report descriptive effect sizes; inference for held-out genes is described below.

Canonical panels were curated from established markers and the atlas annotation; panel development was exploratory. The same source panel was used across destinations where direct comparison was intended. Broader state-associated signatures were derived by one-versus-rest Wilcoxon ranking. Among the top 300 ranked genes per state, candidates required a reported log fold change above 0.5 and adjusted $p < 0.01$. Mitochondrial and ribosomal prefixes (`MT-`, `RPL`, `RPS`), *MALAT1* and *NEAT1* were excluded. The first 80 passing candidates per state were considered; genes shared with another state's candidate list were removed, and up to 50 remaining genes were retained.

Final signatures contained 50 genes each for Mono, Scav, Res and Inflam, 45 for Foam and 49 for Fibro. Signature effects were means of gene-level $\log_2$ fold changes within each stratum, followed by equal subtype averaging within libraries and the median across libraries. The S100-family summary used the same procedure for *S100A8*, *S100A9* and *S100A12*. These summaries are neither fold changes of a summed score nor averages of the displayed gene medians. Full gene sets accompany Supplementary Table 6; broader signature effects are shown in Supplementary Figure 11. Because these signatures were derived from the same atlas, they broaden marker coverage rather than provide an independent validation dataset.

*Matched destination comparison.* The Scav–Inflam versus Scav–Fibro RNA comparison was restricted to library–subtype strata eligible for both gradients. Each gradient retained its own quartiles. Signature effects were calculated using the same gene sets and paired within libraries, using 25 matched strata across 19 libraries (main Fig. 5D). This matched set differs from the separately eligible sets used in Supplementary Figure 11.

*Descriptive plasticity patterns.* Layering denotes increased destination-associated expression together with retained or reinforced source features; selective reconfiguration denotes decreases in particular source features alongside destination-associated increases. The terms are descriptive and nonexclusive. No numerical rule assigned every comparison to a fixed mechanism, and near-zero effects were not treated as an equivalence test or evidence of complete identity preservation. Gene-specific and broader-signature results were considered together.

##### Target resemblance and destination specificity in a shared gene space

Q1 and Q4 source-cell profiles were compared with same-library target reference populations using the union of the exclusive Scav, Inflam, Res, Foam and Fibro signatures, excluding the Mono signature. Genes present in the atlas with expression standard deviation above $10^{-8}$ were retained, yielding a shared 244-gene panel. This is a same-atlas signature comparison, distinct from the held-out-feature analysis below.

Analyses used atlas $\log_e(1+x)$-normalized expression. Each gene was divided by its standard deviation across all atlas cells, without mean centering or conversion back to the linear scale. The scale was fixed across all references within the analysis. The Mono comparison used sample standard deviations (`ddof=1`); the Scav comparison used population standard deviations (`ddof=0`).

Each eligible source library–subtype stratum contained at least 20 Q1 and 20 Q4 cells. Target references were mean profiles of all cells assigned to the target state within the same library, requiring at least 20 target cells. Distance was the root mean squared difference between the source-quartile and target mean profiles:

$$d_{q,\ell h,t} = \sqrt{\frac{1}{G}\sum_{g=1}^{G}\left(\bar{z}_{g,q,\ell h} - \bar{z}_{g,t,\ell}\right)^2}, \qquad G = 244.$$

Here $q$ denotes source quartile, $\ell$ library, $h$ source subtype and $t$ target state. This is Euclidean distance between mean profiles divided by $\sqrt{G}$, not the mean of individual-cell distances. Distance reduction was $\Delta_{\ell h,t} = d_{Q1,\ell h,t} - d_{Q4,\ell h,t}$; positive values indicate greater target resemblance in Q4. Percentage gap closed was $100\Delta_{\ell h,t}/d_{Q1,\ell h,t}$, evaluated for baseline distances above $10^{-8}$ in the Mono analysis and above zero in the Scav analysis.

The Mono–Fibro Q1/Q4 groups were held fixed when comparing Fibro with Scav, Inflam, Foam or Res references wherever eligible. The Scav–Fibro Q1/Q4 groups were held fixed when comparing Fibro and Foam references. Each paired comparison required both references and used identical source strata. Preferential Fibro resemblance was $\Delta_{\ell h,\mathrm{Fibro}} - \Delta_{\ell h,\mathrm{alternative}}$. Reductions, percentages and paired differences were calculated per stratum, averaged equally across eligible subtypes within libraries, and summarized by medians across libraries and the fraction of positive paired differences.

The Mono analysis included seven libraries with Fibro references; paired comparisons used seven libraries for Scav and five each for Inflam and Foam. Median Fibro advantages were 0.144, 0.170 and 0.006, respectively, with positive advantages in 100%, 100% and 60% of libraries. No eligible Mono comparison with a Res reference was reported. The Scav comparison included nine libraries, with median reductions of 0.187 towards Fibro and 0.166 towards Foam; the median paired advantage was 0.044 and favored Fibro in eight of nine libraries. These are descriptive paired effects without a significance test in this analysis.

##### Held-out-feature corroboration of OT associations

Genes were strictly held out if `highly_variable=False` and `highly_variable_nbatches=0`, excluding genes selected as highly variable in any cohort as well as genes in the final integration feature set. Two complementary analyses reused the corrected OT scores and source quartiles (Supplementary Figure 6).

*Gene-level endpoint differences.* Held-out genes detected in at least 20 source cells and at least 1% of source cells were tested with the linearized-expression Q4/Q1 estimator above, requiring at least 20 cells per endpoint within each library–subtype stratum. Eligible subtype effects were averaged within libraries. Before inference, effects from eligible paired fresh and frozen preparations in the Jaiswal cohort were averaged with equal weight into one analysis unit; other libraries remained separate. The only pair with both preparations eligible in these analyses was GSM6976306/GSM6976307. Eligibility, quartile assignments and source-subtype weighting were unchanged. Two-sided tests assessed the mean analysis-unit effect using sign flips that preserved the vector of gene effects from each unit. Exact enumeration was used for 3–12 units and 9,999 Monte Carlo sign patterns for larger samples, with a plus-one correction for Monte Carlo $p$-values. Comparisons with fewer than three units were descriptive only. Unchanged raw test results were retained, and Benjamini–Hochberg adjustment was recalculated across the complete gene–route testing family.

There were 42,415 tested gene–route entries and 2,513 discoveries at global $q < 0.05$: 342 for Mono–Scav, 262 for Mono–Inflam, 91 for Mono–Res, 252 for Mono–Fibro, 760 for Scav–Inflam and 806 for Scav–Fibro. The remaining comparisons, including Mono–Foam, had no discoveries at this threshold. These counts refer to gene–route combinations, not unique genes. Limited numbers of analysis units restrict the resolution of exact two-sided tests: with four nonzero unit effects the minimum attainable $p$ is 0.125, and with three it is 0.25, before multiplicity adjustment. A lack of discoveries therefore does not establish absence of expression differences.

*Same-library target distance.* Panels comprised the top 100, top 500 or all expressed, variable held-out genes ranked by variance across the combined source and target populations, without using OT scores or quartile labels. For each source cell, cosine distance was calculated from its log-normalized expression vector to the mean target-state profile from the same library. References required at least 20 target cells. Cells with zero vector norm were unevaluable; strata additionally required at least 20 valid cells in each endpoint quartile. Within each eligible library–subtype stratum, Spearman correlation was calculated across valid source cells between corrected OT score and target distance. Subtype correlations were averaged equally within libraries and then within known paired preparations using the rule above; medians across analysis units were displayed.

The mean analysis-unit correlation was tested with the sign-flip procedure above, with a separate Benjamini–Hochberg family across route–panel tests. All ten estimable comparisons had negative median correlations across the three panels. Mono–Fibro, Scav–Inflam and Scav–Fibro retained adjusted support across all three panels; Mono–Scav and Mono–Inflam retained support only with the top 100 genes. Mono–Scav had 15 units from 16 libraries. Mono–Res had one unit from two paired libraries and was descriptive only; Res–Fibro had no eligible target-distance comparison. These analyses assess held-out expression features in the same cells, rather than independent cohorts, temporal transitions or exclusive destination specificity.

#### SOLO scores along inflammatory association gradients

Saved SOLO scores from retained atlas cells were joined by exact cell identifier to the Scav–Inflam, Res–Inflam and Mono–Inflam gradient tables. The pooled Q1–Q4 distributions in Supplementary Figure 7A–C are descriptive. Q4-minus-Q1 mean-score differences were calculated within library–subtype strata containing at least 20 cells in each endpoint and averaged equally over eligible subtypes within libraries. This yielded 26, five and 18 matched libraries, respectively. Supplementary Figure 7D–F displays individual paired library means and their arithmetic mean, rather than confidence intervals around pooled cells.

For inference, library differences were grouped using the known-pair rule above, yielding 26, four and 17 analysis units for Scav–Inflam, Res–Inflam and Mono–Inflam, respectively. Two-sided sign-flip tests assessed mean unit differences using exact enumeration for 3–12 units or 9,999 Monte Carlo patterns with a plus-one correction for larger samples. Benjamini–Hochberg adjustment was applied across the three routes. Percentile intervals for the mean used 10,000 bootstrap draws of analysis units. The tests assume independent analysis units and sign-symmetric effects under the null. Supplementary Figure 7 retains the descriptive library-level display; the grouped estimates below need not equal its black-diamond library means.

The mean analysis-unit score changes for Scav–Inflam, Res–Inflam and Mono–Inflam were $+0.011067$, $+0.043974$ and $+0.024451$, respectively. Their 95% percentile-bootstrap intervals were $[-0.017159, 0.039472]$, $[0.026960, 0.060988]$ and $[-0.018392, 0.068475]$; the corresponding sign-flip $p$ values were 0.4713, 0.1250 and 0.2957, and adjusted $q$ values were 0.4713, 0.3750 and 0.44355. Res–Inflam scores increased in all five plotted libraries and all four grouped analysis units. With four nonzero unit effects, the minimum attainable two-sided exact sign-flip $p$ value is $2/2^4 = 0.125$. Bootstrap intervals and exact sign-flip tests use different finite-sample procedures and need not give identical threshold-based conclusions. These results do not establish equivalence or the absence of score enrichment or residual doublets. Original and grouped results, unit membership and per-unit effects are retained in the paired-controls source-data exports.

#### CITE-seq protein corroboration

Processed antibody-derived tag (ADT) counts from the Bashore CITE-seq data (GSE253902) were matched to RNA by library and cell barcode. Analyses retained cells present in the current RNA atlas and the relevant gradient table. The saved ADT object contained 5,455 cells and 283 antibody features; 3,707 cells overlapped the current RNA atlas across six mapped patients (Supplementary Table 8). ADT cells are a subset of RNA cells and were not added to RNA totals.

Raw ADT counts were retained alongside a per-cell centered log-ratio transformation:

$$\mathrm{CLR}_{ip} = \log(1 + c_{ip}) - \frac{1}{P}\sum_{q=1}^{P}\log(1 + c_{iq}),$$

where $c_{ip}$ is the count for antibody $p$ in cell $i$ and $P$ is the number of measured antibody features. The saved `clr` layer supplied the protein values. Effects were mean CLR in Q4 minus mean CLR in Q1, calculated within library–subtype strata with at least 20 cells in each endpoint; subtype effects were averaged equally within libraries and medians across libraries were displayed. CLR differences are relative to the antibody panel and are not RNA $\log_2$ fold changes.

The panel comprised source-associated CD163, CD206, CD169, MERTK and CD64; activation-associated HLA-DR, CD38, CD86 and CD80; lineage controls CD3, CD19, CD56 and CD4; and available isotype controls IgG_Fc, MmuIgG2b and MmuIgG2a. Available antibody identities were matched to their stored feature names. Protein panels were descriptive, without pooled-cell significance stars.

Three eligible libraries (GSM8029956, GSM8029958 and GSM8029960) supported the Scav–Inflam comparison (main Fig. 4C), each within the Lipid-Scavenging subtype. The matched Scav–Inflam versus Scav–Fibro comparison used GSM8029956 and GSM8029958, restricting to source-subtype strata eligible for both gradients while retaining each gradient's quartiles (main Fig. 5C). Individual-library effects are shown for this two-library comparison.

For the same-cell descriptive analysis, a source-associated score was the mean CLR of CD163, CD206 and MERTK, and an activation-associated score was the mean CLR of HLA-DR and CD38. Each library's Q1 median for each score defined its reference threshold. The fraction of cells exceeding both thresholds was calculated separately for Q1 and Q4. These thresholds summarize joint expression and are not validated functional gates.

##### RNA velocity analysis and directional asymmetry

UniTVelo [18] was fitted separately to all 15 meta-cluster pairs in unified-time mode (`FIT_OPTION=1`), with CPU execution and the designated source state supplied through `IROOT` for the primary analysis. Cells with no combined spliced/unspliced counts were removed from the pairwise subset. The predefined ambient-associated gene list was excluded before fitting. Highly variable gene selection requested 2,000 genes, neighbors were calculated in the Scanorama representation with $k = 30$, and spliced/unspliced moments were computed using scVelo [19]. Gene filtering and fitting can yield fewer fitted genes than the requested HVG count.

Saved source-root models supplied the quantitative asymmetry estimates and Supplementary Figure 5. The source mapping uses the v3 fits for Mono–Inflam, Scav–Inflam, Res–Inflam, Inflam–Fibro and Inflam–Foam, and the corresponding v2 fits for the other ten pairs; all use the rebuilt six-state atlas. Projected UMAP velocities were reused or calculated from saved fits for visualization, without refitting during figure generation.

*Transition-support statistic.* For each fitted velocity graph, the row-stochastic scVelo transition matrix $P$ was obtained. For cell $i$ in state $A$, let $N_i = \{j : P_{ij} > 0\}$ and retain interface cells for which $N_i$ contains at least one cell in state $B$. The target-mass enrichment was

$$r_{i,A\to B} = \frac{\sum_{j\in B} P_{ij}}{|N_i \cap B|/|N_i|}.$$

The proportion $q_{A\to B}$ of evaluable interface cells with $r_{i,A\to B} > 1$ was calculated, and the signed asymmetry was

$$\Delta_{AB} = q_{A\to B} - q_{B\to A}.$$

At least ten evaluable cells were required per orientation. The interface is defined by positive transition-matrix support and is velocity-dependent. The primary statistic is not a UMAP cosine difference; the script's separate $k = 20$ UMAP-neighbor diagnostic does not define this interface or classification.

*Sign-flip null and reporting.* Each null draw independently multiplied each cell's entire velocity vector by $+1$ or $-1$ with equal probability and rebuilt the velocity graph and transition matrix. One hundred draws were requested (seed 0); draws without adequate interface coverage were omitted. The two-sided empirical value was

$$p = \frac{1 + \sum_{b=1}^{B_{\text{valid}}} \mathbf{1}\{|\Delta_{AB}^{(b)}| \geq |\Delta_{AB}|\}}{1 + B_{\text{valid}}}.$$

For unadjusted $p < 0.05$, the sign specifies the conditional orientation; otherwise the pair is unresolved. No absolute 0.5 effect cutoff is applied. The minimum attainable value for 100 valid draws is $1/101 \simeq 0.0099$. The reported 95th percentile of null $|\Delta|$ is a null reference, not an estimate confidence interval.

Thirteen pairs had a resolved source-root asymmetry (12 in the designated A-to-B orientation and Res–Fibro in the opposite orientation); Mono–Res and Scav–Res were unresolved. Initialization sensitivity was assessed using opposite-root and unspecified-root fits, with all signs aligned to the same pair order in Supplementary Table 5. Eight of the 13 source-root-resolved orientations reversed under opposite-root initialization; Mono–Fibro, Foam–Fibro and Scav–Fibro retained the same resolved orientation across all three settings. These results describe conditional fitted velocity patterns and are reported separately from the undirected OT relationships and expression gradients. An unresolved asymmetry does not establish biological bidirectionality. Main Fig. 3D displays the five monocyte pairs, Fig. 4D the three inflammatory pairs and Fig. 5E the four Fibro-associated pairs displayed in Fig. 5A (Scav, Foam, Mono and Inflam paired with Fibro). Supplementary Table 5 retains all 15 tested pairs, including Res–Fibro, together with the initialization-sensitivity results.

#### Transcription-factor activity inference

Regulator activities were inferred from current atlas log-normalized expression using the univariate linear model in decoupleR v1.8.0 [20], with human signed CollecTRI regulons [21]. Complexes were retained, and at least five measured targets were required per regulator. The retained network contains 43,159 interactions and 1,186 regulator entries; 762 regulators met the analysis requirements.

For each of the 11 comparisons, mean Q4-minus-Q1 activity differences were calculated within eligible library–subtype strata (at least 20 cells per endpoint), then averaged equally within libraries. Median library effects were reported. Two-sided Wilcoxon signed-rank tests used library effects when at least three libraries were available (`zero_method='wilcox', method='auto'`); all-zero effects were assigned $p = 1$. Benjamini–Hochberg correction was applied jointly across 8,382 regulator–route tests. Supplementary Table 7 prints the five lowest and five highest median effects per route regardless of significance, with all tested regulators retained in the source data. Inferred activities represent regulon-associated expression, not direct TF abundance or experimental evidence of regulation.

#### Sensitivity of signature and regulator summaries to paired preparations

We assessed the influence of known paired fresh and frozen preparations in the Jaiswal cohort using saved library-level effects. Eligible preparations from the same donor were averaged with equal weight into one analysis unit; other libraries remained separate. Only GSM6976306/GSM6976307 contributed both preparations to an eligible comparison. Quartile assignments, source-subtype weighting and gene sets were unchanged. Broad-signature medians and transcription-factor Wilcoxon tests were recomputed using the grouped units, followed by Benjamini–Hochberg correction across all 8,382 regulator–route tests.

All 22 source- and destination-signature medians retained their sign. Among regulator–route tests, 192 no longer met $q < 0.05$ and 15 newly met this threshold; no comparison became untestable. The principal regulatory associations described in the main text retained adjusted statistical support. Supplementary Table 7 reports the primary library-level estimates. This targeted sensitivity analysis accounts for the documented Jaiswal pairing and does not constitute a complete donor-level reanalysis across all cohorts.

##### Clinical association analysis and patient metadata

The clinical analysis (main Fig. 4E) used the documented Bashore library-to-BioSample-to-patient mapping, comprising 18 patients (11 symptomatic, seven asymptomatic). The current atlas contains 16,336 RNA cells from these patients. Each mapped patient contributes one RNA library in the supplied atlas; repeated sequencing runs and ADT records were not counted as additional patients. Age, sex and symptom status were taken from the mapping rather than inferred from expression. Supplementary Table 8 lists all 18 patients, source-cell denominators, state composition and ADT overlap. Analyses were restricted to this mapped cohort; clinical availability in other cohorts was not used to infer missing donor labels.

For Mono–Inflam and Scav–Inflam, the 90th percentile of corrected association scores was calculated across mapped Bashore source cells. Each patient's endpoint was the fraction of their source cells at or above this common route-specific threshold. The threshold was defined across mapped Bashore cells, not separately per patient or across the entire seven-cohort atlas. Patients with at least one source cell and known status contributed one value per route. The 20-cell-per-quartile rule for expression contrasts was not applied to this clinical endpoint; source denominators remain explicit in Supplementary Table 8.

The statistic was the symptomatic-minus-asymptomatic difference in patient medians. Symptom labels were permuted 20,000 times while preserving group sizes. One-sided empirical $p$-values evaluated higher fractions in symptomatic patients, with a plus-one correction. Values were unadjusted. The comparisons were exploratory, without clinical covariate adjustment or an external predictive test.

##### Application to human liver macrophages

Processed human liver matrices from Ramachandran et al. [22] (GSE136103) included healthy and cirrhotic samples. Cells with fewer than 500 detected genes or at least 20% mitochondrial counts were excluded. Expression was normalized to 10,000 counts per cell and log-transformed. Library-based integration used 2,000 highly variable genes and a 50-dimensional Scanorama representation. Broad annotation identified the myeloid compartment, which was re-integrated and clustered. Marker-guided review and subclustering identified tissue monocytes, Kupffer cells, C1q$^+$/MHC-II macrophages and scar-associated macrophages; dendritic and nonmyeloid cells were excluded, leaving 8,536 cells.

Pairwise OT and target-association calculations used the liver representation and state labels. Tissue-monocyte and Kupffer comparisons with scar-associated macrophages used saved Q1/Q4 assignments and the same linearized-expression $\log_2$ fold-change estimator with pseudocount 0.1. Eligibility required at least 20 source cells in each endpoint within a library; effects were summarized by their median across libraries, without the plaque-specific fine-subtype stratification. The source panels were *S100A8*, *S100A12*, *VCAN*, *FCN1* for tissue monocytes and *CD5L*, *MARCO*, *TIMD4*, *VSIG4* for Kupffer cells; the scar-associated panel comprised *CD9*, *GPNMB*, *APOE* and *LGALS3*. This extension evaluated the comparison strategy in another tissue and did not include RNA velocity.

Thirteen monocyte libraries met the eligibility threshold (seven healthy, six cirrhotic), whereas five Kupffer-cell libraries were eligible (four healthy, one cirrhotic). These are library counts, not necessarily independent donor counts. The corrected median library effects showed decreases in all four selected monocyte markers and increases in all four scar-associated markers along monocyte–SAM association. Along

Kupffer-cell–SAM association, all four selected source markers increased, as did *GPNMB*, *APOE* and *LGALS3*, whereas *CD9* decreased (median $\log_2$ fold change $= -0.928829$). The Kupffer comparison therefore does not demonstrate replication across multiple cirrhotic donors, and scar-associated features were not uniformly acquired. Gene effects and eligibility are recorded in `fig6C_values.csv` and `fig6C_coverage.csv`; library-level effects are retained in `fig6C_per_library.csv`.

**Statistical units, software and source data**

Cells, library–subtype strata, libraries and patients were treated as separate units. RNA, protein and regulatory contrasts used library summaries; the symptom analysis used patients. For held-out-gene, held-out-distance and SOLO inference, known paired Jaiswal preparations were combined into analysis units as specified above; other libraries remained separate. These tests assume independence between analysis units and sign-symmetric effects under the null. This grouping accounts for the documented pairing but does not constitute a complete donor-level reanalysis across all cohorts. Gene/marker and protein summaries were primarily descriptive. Multiple-testing families were defined separately for held-out genes, held-out-distance panels, SOLO comparisons and regulator activities, as specified above. Clinical and primary velocity permutation values were unadjusted.

Missing or ineligible comparisons were retained as unavailable rather than assigned zero. Supplementary Tables 1–9 print the principal results and identify their complete machine-readable exports. Table 6 includes 123 canonical marker rows from 1,133 gene–route–role entries, and Table 7 includes 110 effect-ranked entries from 8,382 tests. Companion per-cell, per-stratum, per-library and provenance files retain the respective denominators and inputs. Supplementary figure source data retain plotted values and coverage. The table source-data folder names below follow the supplied `table_1`–`table_9` archive.

The table-export environment recorded NumPy 2.0.2, pandas 2.3.3, anndata 0.12.9, SciPy 1.15.3, harmonypy 0.0.10 and decoupleR 1.8.0. These versions describe that export environment, not every preprocessing or velocity environment. Analysis scripts and run-specific settings are retained with the study code and output provenance.

#### List of Supplementary Tables

**Supplementary Table 1.** Batch correction benchmark.

**Supplementary Table 2.** Cohort mixing and enrichment within meta-clusters.

**Supplementary Table 3.** Current atlas annotation and marker panels.

**Supplementary Table 4.** Pairwise optimal transport and leave-one-cohort-out analysis.

**Supplementary Table 5.** Conditional RNA velocity asymmetry and initialization sensitivity.

**Supplementary Table 6.** Gene-level expression changes along OT association gradients.

**Supplementary Table 7.** Inferred transcription factor activity along OT association gradients.

**Supplementary Table 8.** Bashore patient metadata and macrophage composition.

**Supplementary Table 9.** Quality control settings and cell accounting.

#### Supplementary Table 1

##### Batch correction benchmark.

Four integration methods were evaluated on the same 20,576 cells drawn from the current 34,533-cell atlas, comprising seven cohorts and 50 libraries. Values reproduce the saved benchmark.

| Method | Batch ASW | Graph connectivity | PCR | Composite |
|---|---|---|---|---|
| Harmony | 0.926 | 0.985 | 0.655 | 0.855 |
| scVI | 0.932 | 0.996 | 0.553 | 0.827 |
| Scanorama | 0.934 | 0.998 | 0.424 | 0.786 |
| BBKNN[†] | 0.911 | 0.996 | — | — |

**Metrics.** Batch ASW measures cohort mixing within the annotated cell types; graph connectivity measures connectivity within cell-type groups. Both use the existing annotations. PCR measures the reduction in variance associated with cohort. The composite is the arithmetic mean of the three saved metrics; larger values indicate better performance under their respective definitions.

[†]BBKNN produces a corrected neighbour graph rather than a corrected embedding. Its saved PCR value (0) and resulting composite (0.636) are retained in the source CSV but omitted from the embedding-based comparison here. These metrics do not establish comparative performance for downstream state-transition inference.

**Source data.** `table_1/Table_1.csv`, `table_1/benchmark_coverage.csv`, `table_1/benchmark_cohort_counts.csv`.

#### Supplementary Table 2

##### Cohort mixing and enrichment within meta-clusters.

Local Inverse Simpson Index (iLISI) was calculated with `harmonypy` (perplexity 30) in the Scanorama embedding, separately within each meta-cluster and without downsampling. All six populations contain cells from all seven cohorts.

| Meta-cluster | Cells | Median iLISI | Mean iLISI | Normalised median | Composition diversity | UMAP spread | Dominant cohort | Dominant fraction | $\Delta$ (pp) |
|---|---|---|---|---|---|---|---|---|---|
| Mono | 9,961 | 1.239 | 1.513 | 0.0399 | 4.051 | 1.882 | Jaiswal | 31.6% | +18.6 |
| Scav | 13,499 | 1.448 | 1.627 | 0.0747 | 2.579 | 1.092 | Bashore | 57.2% | +9.9 |
| Res | 1,409 | 1.017 | 1.142 | 0.0029 | 2.565 | 2.915 | Jaiswal | 47.6% | +34.6 |
| Inflam | 2,309 | 1.249 | 1.463 | 0.0415 | 1.839 | 0.959 | Bashore | 72.1% | +24.8 |
| Foam | 3,240 | 1.015 | 1.204 | 0.0025 | 1.864 | 1.190 | Alsaigh | 69.5% | +47.4 |
| Fibro | 4,115 | 1.155 | 1.415 | 0.0258 | 1.614 | 1.588 | Bashore | 77.7% | +30.4 |

**Definitions.** Raw iLISI is the effective local number of cohorts (range 1–7); normalised iLISI is $(\mathrm{iLISI}-1)/6$. Composition diversity is $1/\sum_c p_c^2$, calculated from the cohort fractions of the entire meta-cluster, and is a reference mixture rather than a local-mixing ceiling. UMAP spread is the unweighted mean pairwise distance between cohort centroids. Dominant-cohort enrichment is the percentage-point difference between that cohort's fraction in the meta-cluster and its fraction in the full atlas.

**Interpretation.** The observed local iLISI values indicate substantial cohort concentration within these broad populations. Cohort representation is unequal; the metrics alone do not distinguish technical effects from differences in the sampled cell states or tissue composition.

**Cohort composition of each meta-cluster (%).**

| Meta-cluster | Alsaigh | Bashore | Fernandez | Jaiswal | Pan | Pauli | Wirka |
|---|---|---|---|---|---|---|---|
| Mono | 19.7 | 30.2 | 12.5 | 31.6 | 2.9 | 0.2 | 3.0 |
| Scav | 19.8 | 57.2 | 0.2 | 1.7 | 10.5 | 0.7 | 9.7 |
| Res | 11.5 | 1.6 | 0.3 | 47.6 | 0.2 | 0.1 | 38.7 |
| Inflam | 5.1 | 72.1 | 0.0 | 1.2 | 11.8 | 1.0 | 8.8 |
| Foam | 69.5 | 22.3 | 1.4 | 6.2 | 0.1 | 0.3 | 0.2 |
| Fibro | 11.0 | 77.7 | 0.6 | 5.0 | 1.2 | 4.1 | 0.4 |

**Source data.** `table_2/Table_2.csv`, `table_2/cohort_enrichment.csv`, `table_2/cell_iLISI.csv`.

#### Supplementary Table 3

##### Current atlas annotation and marker panels.

Nine nonempty annotated subtypes map to six meta-clusters in the current atlas (34,533 cells, 50 libraries, seven cohorts). Cell counts, subtype assignments and cluster identifiers are read from the saved atlas. All nine subtypes occur in all seven cohorts.

| Meta-cluster | Annotated subtype | Cells | Atlas (%) | Libraries | Canonical marker panel | Stored cluster identifiers |
|---|---|---|---|---|---|---|
| Mono | Classical Monocytes | 5,128 | 14.85 | 49 | *S100A8*; *S100A9*; *FCN1* | r0.8:2 |
| Mono | Intermediate / Non-Classical Monocytes | 2,796 | 8.10 | 46 | *LST1*; *S100A4*; *FCGR3A* | r0.8:11; r0.8:13 |
| Mono | Transitional Monocytes | 2,037 | 5.90 | 46 | *VCAN*; *ZEB2* | r0.8:10 |
| Scav | Homeostatic Lipid-Regulating Macrophages | 4,614 | 13.36 | 44 | *APOE*; *SELENOP* | r0.8:3 |
| Scav | Lipid-Scavenging Macrophages | 8,885 | 25.73 | 48 | *C1QA*; *C1QB*; *HLA-DRA*; *MSR1* | r0.8:1; r0.8:s0,1; r0.8:s0,12; r0.8:s0,2; r0.8:s0,3; r0.8:s0,4; r0.8:s0,5; r0.8:s0,7 |
| Res | Resident Perivascular Macrophages | 1,409 | 4.08 | 29 | *LYVE1*; *FOLR2*; *F13A1* | r0.8:12 |
| Inflam | Inflammatory Macrophages | 2,309 | 6.69 | 44 | *CCL3*; *CCL4*; *IL1B*; *CXCL8* | r0.8:s0,0; r0.8:s0,10; r0.8:s0,11; r0.8:s0,6; r0.8:s0,8; r0.8:s0,9 |
| Foam | Lipid-Stressed Foam Cells | 3,240 | 9.38 | 37 | *FABP5*; *CSTB*; *PLIN2*; *LIPA* | r0.8:6 |
| Fibro | Fibrotic Macrophages | 4,115 | 11.92 | 46 | *SPP1*; *FN1*; *ANXA2*; *LGALS1* | r0.8:4 |

**Annotation and markers.** The listed genes are canonical annotation panels, not newly ranked differential-expression results. The stored `r0.8:` prefix denotes the resolution recorded with the cluster identifiers; `s0,` identifiers record subdivisions of the parent cluster. Subtype labels consolidate these stored clusters. Quantitative marker support reports mean $\log_e$(1+normalised expression) and the percentage of cells expressing each gene for every panel gene across every subtype (261 gene–subtype comparisons). Population names describe expression-defined states and do not establish cellular origin.

**Source data.** `table_3/Table_3.csv`, `table_3/marker_expression_all_subtypes.csv`, `table_3/meta_counts.csv`, `table_3/subtype_cohort_counts.csv`.

#### Supplementary Table 4

##### Pairwise optimal transport and leave-one-cohort-out analysis.

All 15 unordered meta-cluster pairs were evaluated using the current atlas and Scanorama embedding. The saved reference calculation uses up to 5,000 cells per state, entropic regularisation $\epsilon = 0.01$, and seed 42. Library-bootstrap summaries use 200 resampling iterations and up to 2,000 cells per state. The underlying run also contains 500 cell-bootstrap iterations and 50 label permutations.

**A. Reference estimates and library-resampling summaries.**

| Pair | Reference divergence | Connectivity | Bootstrap mean | Bootstrap SD | Bootstrap 2.5–97.5% interval | Null $z$ |
|---|---|---|---|---|---|---|
| Mono–Scav | 0.3203 | 0.2978 | 0.2707 | 0.0114 | [0.2509, 0.2968] | 132.96 |
| Mono–Res | 0.4002 | 0.1509 | 0.3746 | 0.0246 | [0.3302, 0.4235] | 139.12 |
| Mono–Inflam | 0.3940 | 0.1599 | 0.3492 | 0.0180 | [0.3156, 0.3805] | 158.76 |
| Mono–Foam | 0.3941 | 0.1597 | 0.3243 | 0.0280 | [0.2857, 0.3725] | 147.81 |
| Mono–Fibro | 0.4386 | 0.1031 | 0.3543 | 0.0206 | [0.3253, 0.4081] | 156.24 |
| Scav–Res | 0.3364 | 0.2628 | 0.3183 | 0.0104 | [0.2928, 0.3380] | 107.33 |
| Scav–Inflam | 0.2739 | 0.4124 | 0.2392 | 0.0171 | [0.2066, 0.2680] | 81.02 |
| Scav–Foam | 0.4115 | 0.1353 | 0.3508 | 0.0280 | [0.3123, 0.3986] | 158.32 |
| Scav–Fibro | 0.3622 | 0.2124 | 0.2833 | 0.0187 | [0.2552, 0.3237] | 137.23 |
| Res–Inflam | 0.4870 | 0.0607 | 0.4725 | 0.0163 | [0.4396, 0.4999] | 180.40 |
| Res–Foam | 0.4854 | 0.0619 | 0.4518 | 0.0395 | [0.3955, 0.5362] | 164.88 |
| Res–Fibro | 0.4743 | 0.0701 | 0.4210 | 0.0205 | [0.3863, 0.4624] | 143.23 |
| Inflam–Foam | 0.5204 | 0.0408 | 0.4631 | 0.0278 | [0.4144, 0.5145] | 232.41 |
| Inflam–Fibro | 0.5061 | 0.0485 | 0.4307 | 0.0227 | [0.3913, 0.4740] | 186.91 |
| Foam–Fibro | 0.4127 | 0.1338 | 0.3225 | 0.0400 | [0.2647, 0.3991] | 124.59 |

Connectivity is the saved Gaussian transformation $\exp[-d^2/(2\sigma^2)]$ of the reference divergence $d$, with $\sigma = 0.205758$. Higher connectivity denotes greater transcriptional proximity. Bootstrap intervals describe the library-resampling distribution under its smaller cell cap; they are reported separately from the reference estimates rather than as matched confidence bounds around them. The $z$-score compares reference divergence with the label-permutation null. It quantifies separation from label mixing, not evidence that a biological transition occurs, and is not an empirical permutation $p$-value.

**B. Leave-one-cohort-out reference divergences.**

| Pair | Alsaigh | Bashore | Fernandez | Jaiswal | Pan | Pauli | Wirka |
|---|---|---|---|---|---|---|---|
| Mono–Scav | 0.3489 | 0.4240 | 0.3235 | 0.3492 | 0.3295 | 0.3183 | 0.3261 |
| Mono–Res | 0.4050 | 0.4333 | 0.4072 | 0.4791 | 0.4024 | 0.3978 | 0.3830 |
| Mono–Inflam | 0.4108 | 0.4397 | 0.3963 | 0.4219 | 0.3980 | 0.3929 | 0.3962 |
| Mono–Foam | 0.3987 | 0.4301 | 0.4012 | 0.4226 | 0.3951 | 0.3928 | 0.3965 |
| Mono–Fibro | 0.4524 | 0.4745 | 0.4460 | 0.4652 | 0.4398 | 0.4356 | 0.4392 |
| Scav–Res | 0.3491 | 0.4037 | 0.3358 | 0.3488 | 0.3435 | 0.3351 | 0.3449 |
| Scav–Inflam | 0.2900 | 0.3495 | 0.2745 | 0.2736 | 0.2805 | 0.2721 | 0.2794 |
| Scav–Foam | 0.4186 | 0.4838 | 0.4128 | 0.4124 | 0.4147 | 0.4110 | 0.4171 |
| Scav–Fibro | 0.3760 | 0.4435 | 0.3615 | 0.3602 | 0.3667 | 0.3605 | 0.3696 |
| Res–Inflam | 0.4860 | 0.4947 | 0.4870 | 0.5062 | 0.4861 | 0.4870 | 0.4840 |
| Res–Foam | 0.4804 | 0.4882 | 0.4853 | 0.5323 | 0.4855 | 0.4853 | 0.4637 |
| Res–Fibro | 0.4712 | 0.4699 | 0.4746 | 0.5072 | 0.4749 | 0.4747 | 0.4664 |
| Inflam–Foam | 0.5096 | 0.5293 | 0.5205 | 0.5209 | 0.5195 | 0.5204 | 0.5203 |
| Inflam–Fibro | 0.5082 | 0.5123 | 0.5062 | 0.5052 | 0.5062 | 0.5048 | 0.5058 |
| Foam–Fibro | 0.4245 | 0.4009 | 0.4127 | 0.4127 | 0.4125 | 0.4143 | 0.4126 |

Each column omits the named cohort, with all other reference-run settings held constant. Raw divergences are reported because connectivity scaling is recalculated in each run.

**C. Pair-ranking sensitivity to cohort omission.**

| Omitted cohort | Retained cells | Rank correlation | Largest divergence change | Largest rank change |
|---|---|---|---|---|
| Alsaigh | 26,914 | 0.989 | 0.0286 | 2 |
| Bashore | 18,197 | 0.843 | 0.1037 | 7 |
| Fernandez | 33,183 | 0.996 | 0.0074 | 1 |
| Jaiswal | 30,045 | 0.889 | 0.0789 | 3 |
| Pan | 32,496 | 0.993 | 0.0092 | 1 |
| Pauli | 34,214 | 0.996 | 0.0030 | 1 |
| Wirka | 32,149 | 0.982 | 0.0217 | 2 |

Ranks increase with divergence (rank 1 is the closest pair). Correlations compare all 15 pair ranks with the reference run. Pair-level rank changes and run settings are retained in the source data. Velocity orientation is reported separately in Supplementary Table 5.

**Source data.** `table_4/Table_4.csv`, `table_4/all_LOCO_pairs.csv`, `table_4/run_configs.csv`.

#### Supplementary Table 5

##### Conditional RNA velocity asymmetry and initialization sensitivity.

Pairwise UniTVelo estimates are reported under source-root initialization (state $A$ specified as the root), together with alternate-root and unspecified-root sensitivity results. For each pair, $q_{A\to B}$ is the proportion of interface cells in $A$ whose transition mass towards $B$ exceeds the fraction of their positive-support transition neighbours belonging to $B$. Interface cells have positive transition-matrix support to the other state. The signed statistic is

$$\Delta_{AB} = q_{A\to B} - q_{B\to A}.$$

The interface support is velocity-dependent. This statistic is not a difference between mean cosine similarities.

**A. Source-root conditional estimates.**

| **Pair** $(A, B)$ | $q_{A\to B}$ | $q_{B\to A}$ | $\Delta_{AB}$ | **Null** $\lvert\Delta\rvert$ **95th pct.** | $p$ | **Interface cells A / B** | **Conditional orientation** |
|---|---|---|---|---|---|---|---|
| Mono–Inflam | 0.9952 | 0.0000 | +0.9952 | 0.0317 | 0.0099 | 2,914 / 2,037 | Mono $\to$ Inflam |
| Mono–Scav | 0.9976 | 0.0000 | +0.9976 | 0.1172 | 0.0099 | 3,396 / 8,397 | Mono $\to$ Scav |
| Mono–Foam | 0.9331 | 0.0000 | +0.9331 | 0.1549 | 0.0099 | 3,200 / 1,342 | Mono $\to$ Foam |
| Mono–Res | 0.2583 | 0.2254 | +0.0329 | 0.2943 | 1.0000 | 3,120 / 865 | Unresolved |
| Mono–Fibro | 0.9318 | 0.0031 | +0.9287 | 0.2521 | 0.0099 | 1,599 / 1,295 | Mono $\to$ Fibro |
| Scav–Inflam | 0.6638 | 0.0932 | +0.5706 | 0.0725 | 0.0099 | 10,763 / 2,295 | Scav $\to$ Inflam |
| Res–Inflam | 0.9637 | 0.0000 | +0.9637 | 0.1086 | 0.0099 | 331 / 165 | Res $\to$ Inflam |
| Foam–Fibro | 0.9849 | 0.0008 | +0.9841 | 0.1751 | 0.0099 | 2,047 / 2,657 | Foam $\to$ Fibro |
| Inflam–Fibro | 0.8853 | 0.0000 | +0.8853 | 0.2642 | 0.0099 | 497 / 916 | Inflam $\to$ Fibro |
| Scav–Fibro | 0.9921 | 0.0000 | +0.9921 | 0.0236 | 0.0099 | 7,005 / 2,585 | Scav $\to$ Fibro |
| Res–Fibro | 0.0000 | 0.8006 | -0.8006 | 0.0892 | 0.0099 | 383 / 1,053 | Fibro $\to$ Res |
| Scav–Res | 0.4055 | 0.1538 | +0.2517 | 0.2951 | 0.9604 | 3,623 / 1,086 | Unresolved |
| Res–Foam | 1.0000 | 0.0000 | +1.0000 | 0.0741 | 0.0099 | 302 / 464 | Res $\to$ Foam |
| Inflam–Foam | 1.0000 | 0.0000 | +1.0000 | 0.1352 | 0.0099 | 87 / 191 | Inflam $\to$ Foam |
| Scav–Foam | 1.0000 | 0.0000 | +1.0000 | 0.0660 | 0.0099 | 1,216 / 793 | Scav $\to$ Foam |

**Null and classification.** Each randomization independently reverses cell velocity vectors with probability 0.5 and rebuilds the velocity graph and transition matrix. Two-sided empirical $p$-values compare absolute asymmetry with this sign-flip null, using the plus-one correction and 100 requested randomizations. The minimum attainable value with 100 valid randomizations is $1/101 \simeq 0.0099$. A conditional orientation is assigned when unadjusted $p < 0.05$; its sign determines the orientation. No absolute 0.5 asymmetry cutoff is used. “Unresolved” does not establish biological bidirectionality. Values reproduce the saved four-decimal outputs.

**B. Initialization sensitivity, aligned to the same A-to-B orientation.**

| Pair $(A, B)$ | A-root $\Delta$ | A-root $p$ | B-root $\Delta$ | B-root $p$ | No specified root $\Delta$ | No specified root $p$ | Same resolved orientation |
|---|---|---|---|---|---|---|---|
| Mono–Inflam | +0.9952 | 0.0099 | -0.8581 | 0.0099 | +0.9914 | 0.0099 | No |
| Mono–Scav | +0.9976 | 0.0099 | -0.9050 | 0.0099 | +0.9814 | 0.0099 | No |
| Mono–Foam | +0.9331 | 0.0099 | -0.9993 | 0.0099 | -1.0000 | 0.0099 | No |
| Mono–Res | +0.0329 | 1.0000 | -0.1581 | 1.0000 | -0.5406 | 0.0099 | No |
| Mono–Fibro | +0.9287 | 0.0099 | +0.8913 | 0.0099 | +0.9961 | 0.0099 | Yes |
| Scav–Inflam | +0.5706 | 0.0099 | +0.5680 | 0.0099 | -0.6421 | 0.0099 | No |
| Res–Inflam | +0.9637 | 0.0099 | -0.9939 | 0.0099 | +0.9940 | 0.0099 | No |
| Foam–Fibro | +0.9841 | 0.0099 | +0.9808 | 0.0099 | +0.9941 | 0.0099 | Yes |
| Inflam–Fibro | +0.8853 | 0.0099 | -1.0000 | 0.0099 | +0.9457 | 0.0099 | No |
| Scav–Fibro | +0.9921 | 0.0099 | +0.9935 | 0.0099 | +0.6061 | 0.0099 | Yes |
| Res–Fibro | -0.8006 | 0.0099 | -0.9839 | 0.0099 | +1.0000 | 0.0099 | No |
| Scav–Res | +0.2517 | 0.9604 | +0.1640 | 1.0000 | -1.0000 | 0.0099 | No |
| Res–Foam | +1.0000 | 0.0099 | -0.9871 | 0.0099 | -0.9957 | 0.0099 | No |
| Inflam–Foam | +1.0000 | 0.0099 | -1.0000 | 0.0099 | -1.0000 | 0.0099 | No |
| Scav–Foam | +1.0000 | 0.0099 | -0.9912 | 0.0099 | -0.9950 | 0.0099 | No |

All signs are expressed relative to the printed $A \to B$ order, including models saved in the opposite pair order. “Same resolved orientation” requires the same nonzero sign and $p < 0.05$ in all three settings. Of 13 source-root-resolved pairs, eight reverse their inferred orientation after swapping the specified root. Three pairs retain the same resolved orientation across all three settings: Mono–Fibro, Foam–Fibro and Scav–Fibro. These estimates therefore describe model- and initialization-dependent velocity patterns; significance under the sign-flip null does not establish root-independent temporal transitions.

**Source data.** `table_5/Table_5.csv`, `table_5/Table_5_root_sensitivity.csv`.

#### Supplementary Table 6

##### Gene-level expression changes along OT association gradients.

Gene-level contrasts compare source-population cells in Q4 (highest corrected OT association with the named destination) with Q1 (lowest association). Natural-log1p-normalised expression was transformed back to the linear scale before calculating means. Eligible library–subtype strata contain at least 20 cells in each endpoint quartile. For each gene,

$$\log_2 \mathrm{FC}_{\ell,s} = \log_2\left(\frac{\overline{x}_{Q4,\ell,s} + 0.1}{\overline{x}_{Q1,\ell,s} + 0.1}\right).$$

Subtype effects were averaged equally within each library, followed by the median across libraries. The 123 canonical marker rows are shown below; the full source table contains 1,133 gene–route–role rows covering canonical panels and broad signatures. Association arrows label the analysed source and destination and do not establish temporal direction.

| Gene | Panel role | Median $\log_2$FC | Library-effect IQR | Positive libraries (%) | Mean Q1 | Mean Q4 |
|---|---|---|---|---|---|---|
| **Mono → Scav (17 libraries; 22 library–subtype strata)** | | | | | | |
| *S100A8* | Source | -1.817 | [-1.924, -1.121] | 0.0 | 44.966 | 12.580 |
| *S100A9* | Source | -1.682 | [-2.006, -1.113] | 0.0 | 70.559 | 23.647 |
| *S100A12* | Source | -1.829 | [-2.243, -1.108] | 0.0 | 4.324 | 0.933 |
| *FCN1* | Source | -0.468 | [-0.679, -0.306] | 5.9 | 9.600 | 6.957 |
| *VCAN* | Source | -0.849 | [-1.100, -0.555] | 0.0 | 20.969 | 10.794 |
| *LYZ* | Source | -0.221 | [-0.414, -0.131] | 23.5 | 61.415 | 50.104 |
| *LST1* | Source | -0.062 | [-0.153, 0.178] | 47.1 | 5.576 | 5.476 |
| *COTL1* | Source | -0.364 | [-0.533, -0.130] | 23.5 | 8.693 | 7.492 |
| *SAMSN1* | Source | +0.012 | [-0.206, 0.294] | 52.9 | 10.576 | 11.081 |
| *NAMPT* | Source | +0.150 | [0.029, 0.390] | 76.5 | 24.040 | 27.816 |
| *AREG* | Source | +0.296 | [-0.043, 0.767] | 70.6 | 12.076 | 15.395 |
| *C1QA* | Destination | +1.207 | [0.886, 2.250] | 100.0 | 3.352 | 5.778 |
| *C1QB* | Destination | +1.220 | [0.603, 2.080] | 88.2 | 2.704 | 4.242 |
| *APOE* | Destination | +0.896 | [0.058, 2.097] | 76.5 | 1.407 | 1.223 |
| *SELENOP* | Destination | +1.943 | [0.909, 2.801] | 100.0 | 0.187 | 2.037 |
| **Mono → Inflam (18 libraries; 23 library–subtype strata)** | | | | | | |
| *S100A8* | Source | -0.903 | [-1.610, -0.366] | 11.1 | 48.029 | 30.096 |
| *S100A9* | Source | -0.912 | [-1.580, -0.437] | 11.1 | 72.203 | 48.172 |

*Continued on next page*

| Gene | Panel role | Median $\log_2$FC | Library-effect IQR | Positive libraries (%) | Mean Q1 | Mean Q4 |
|---|---|---|---|---|---|---|
| *S100A12* | Source | -0.809 | [-1.475, -0.362] | 16.7 | 4.969 | 3.020 |
| *FCN1* | Source | -0.106 | [-0.325, -0.012] | 16.7 | 10.883 | 10.034 |
| *VCAN* | Source | -0.532 | [-0.673, -0.309] | 5.6 | 19.612 | 12.945 |
| *LYZ* | Source | -0.063 | [-0.314, 0.066] | 38.9 | 58.341 | 55.928 |
| *LST1* | Source | +0.207 | [-0.003, 0.331] | 72.2 | 6.668 | 7.590 |
| *COTL1* | Source | -0.143 | [-0.327, 0.052] | 33.3 | 9.413 | 8.589 |
| *SAMSN1* | Source | +0.429 | [0.297, 0.626] | 94.4 | 8.952 | 12.352 |
| *NAMPT* | Source | +0.326 | [0.122, 0.475] | 83.3 | 22.218 | 27.064 |
| *AREG* | Source | +0.634 | [0.132, 0.947] | 83.3 | 10.620 | 15.562 |
| *CCL3* | Destination | +0.509 | [0.110, 1.543] | 83.3 | 5.484 | 9.978 |
| *CCL4* | Destination | +0.622 | [-0.047, 1.674] | 66.7 | 7.091 | 8.508 |
| *IL1B* | Destination | +1.108 | [0.533, 1.232] | 94.4 | 8.636 | 17.714 |
| *CXCL8* | Destination | +0.362 | [0.076, 0.879] | 83.3 | 10.234 | 12.070 |
| **Mono $\rightarrow$ Foam (10 libraries; 16 library–subtype strata)** | | | | | | |
| *S100A8* | Source | -0.043 | [-0.888, 0.978] | 50.0 | 43.413 | 17.154 |
| *S100A9* | Source | +0.074 | [-0.474, 0.564] | 50.0 | 46.493 | 31.452 |
| *S100A12* | Source | -0.737 | [-1.226, -0.438] | 10.0 | 4.748 | 1.701 |
| *FCN1* | Source | -0.014 | [-0.339, 0.713] | 50.0 | 9.677 | 8.755 |
| *VCAN* | Source | -0.317 | [-0.752, 0.001] | 30.0 | 13.526 | 9.530 |
| *LYZ* | Source | -0.285 | [-0.430, -0.148] | 10.0 | 71.499 | 55.473 |
| *LST1* | Source | +0.082 | [-0.247, 0.208] | 60.0 | 7.087 | 7.027 |
| *COTL1* | Source | +0.191 | [0.010, 0.598] | 80.0 | 7.244 | 8.634 |
| *SAMSN1* | Source | -0.776 | [-0.891, -0.673] | 10.0 | 10.504 | 6.210 |
| *NAMPT* | Source | -0.653 | [-0.833, -0.313] | 20.0 | 27.313 | 17.740 |
| *AREG* | Source | -0.875 | [-1.332, -0.463] | 10.0 | 14.142 | 7.173 |
| *FABP5* | Destination | +1.923 | [1.021, 2.325] | 100.0 | 0.970 | 2.863 |
| *PLIN2* | Destination | +0.286 | [0.023, 0.783] | 100.0 | 2.654 | 3.667 |
| *CSTB* | Destination | +1.208 | [0.950, 1.324] | 100.0 | 1.484 | 3.410 |
| *LIPA* | Destination | +0.542 | [0.423, 0.908] | 100.0 | 0.734 | 1.210 |

*Continued on next page*

| Gene | Panel role | Median $\log_2$FC | Library-effect IQR | Positive libraries (%) | Mean Q1 | Mean Q4 |
|---|---|---|---|---|---|---|
| **Mono $\rightarrow$ Res (15 libraries; 17 library–subtype strata)** | | | | | | |
| *S100A8* | Source | +0.415 | [-0.193, 0.900] | 66.7 | 41.784 | 46.384 |
| *S100A9* | Source | +0.665 | [0.054, 0.944] | 73.3 | 53.180 | 67.288 |
| *S100A12* | Source | +0.554 | [-0.348, 0.813] | 66.7 | 4.146 | 4.700 |
| *FCN1* | Source | +0.407 | [0.104, 0.555] | 86.7 | 9.930 | 12.499 |
| *VCAN* | Source | +0.162 | [-0.156, 0.531] | 60.0 | 18.391 | 18.338 |
| *LYZ* | Source | +0.104 | [0.032, 0.419] | 80.0 | 63.893 | 73.110 |
| *LST1* | Source | +0.147 | [-0.061, 0.214] | 60.0 | 6.249 | 6.475 |
| *COTL1* | Source | -0.091 | [-0.208, 0.118] | 40.0 | 8.889 | 8.497 |
| *SAMSN1* | Source | -0.270 | [-0.603, -0.022] | 13.3 | 11.419 | 9.136 |
| *NAMPT* | Source | -0.127 | [-0.345, 0.017] | 26.7 | 27.984 | 22.914 |
| *AREG* | Source | -1.213 | [-1.472, -0.240] | 6.7 | 20.073 | 10.438 |
| *LYVE1* | Destination | +0.826 | [0.064, 1.251] | 80.0 | 0.104 | 0.404 |
| *F13A1* | Destination | +1.161 | [0.465, 1.453] | 93.3 | 1.301 | 3.036 |
| *MRC1* | Destination | +0.932 | [0.297, 1.457] | 80.0 | 1.174 | 2.377 |
| *CFD* | Destination | +0.660 | [0.576, 0.810] | 93.3 | 2.892 | 4.781 |
| **Mono $\rightarrow$ Fibro (12 libraries; 14 library–subtype strata)** | | | | | | |
| *S100A8* | Source | +1.051 | [0.649, 1.763] | 100.0 | 29.554 | 80.111 |
| *S100A9* | Source | +1.076 | [0.969, 1.444] | 100.0 | 39.619 | 110.099 |
| *S100A12* | Source | +1.389 | [1.071, 1.835] | 100.0 | 2.565 | 8.355 |
| *FCN1* | Source | +0.481 | [0.242, 0.746] | 100.0 | 10.675 | 16.265 |
| *VCAN* | Source | +0.412 | [0.212, 0.556] | 83.3 | 16.443 | 21.119 |
| *LYZ* | Source | +0.464 | [0.182, 0.655] | 83.3 | 53.817 | 76.200 |
| *LST1* | Source | -0.062 | [-0.211, 0.110] | 41.7 | 6.729 | 6.495 |
| *COTL1* | Source | +0.164 | [-0.023, 0.408] | 66.7 | 8.380 | 9.676 |
| *SAMSN1* | Source | -0.001 | [-0.170, 0.159] | 50.0 | 12.763 | 11.335 |
| *NAMPT* | Source | -0.339 | [-0.459, -0.124] | 16.7 | 33.557 | 24.473 |
| *AREG* | Source | +0.295 | [0.043, 0.496] | 75.0 | 14.629 | 17.843 |
| *SPP1* | Destination | +1.183 | [0.339, 2.204] | 83.3 | 0.326 | 2.521 |
| *FN1* | Destination | +0.423 | [-0.158, 1.554] | 58.3 | 0.552 | 1.157 |

*Continued on next page*

| Gene | Panel role | Median $\log_2$FC | Library-effect IQR | Positive libraries (%) | Mean Q1 | Mean Q4 |
|---|---|---|---|---|---|---|
| *ANXA2* | Destination | +0.571 | [0.061, 0.730] | 75.0 | 3.792 | 5.198 |
| *LGALS1* | Destination | +0.682 | [0.481, 1.027] | 100.0 | 7.851 | 12.721 |
| **Scav → Inflam (26 libraries; 33 library–subtype strata)** | | | | | | |
| *C1QA* | Source | +0.346 | [0.135, 0.441] | 80.8 | 30.892 | 37.911 |
| *C1QB* | Source | +0.467 | [0.250, 0.579] | 100.0 | 28.937 | 38.200 |
| *APOE* | Source | +0.851 | [0.412, 1.632] | 80.8 | 17.060 | 19.383 |
| *SELENOP* | Source | +0.799 | [0.439, 1.394] | 100.0 | 15.036 | 26.645 |
| *CCL3* | Destination | +0.815 | [0.595, 1.636] | 84.6 | 13.396 | 24.480 |
| *CCL4* | Destination | +0.817 | [0.451, 1.789] | 84.6 | 16.517 | 27.366 |
| *IL1B* | Destination | +1.238 | [0.902, 1.842] | 100.0 | 2.516 | 6.975 |
| *CXCL8* | Destination | +1.035 | [0.124, 1.632] | 76.9 | 5.031 | 10.740 |
| **Res → Inflam (5 libraries; 5 library–subtype strata)** | | | | | | |
| *LYVE1* | Source | -0.198 | [-0.314, 0.822] | 40.0 | 11.301 | 9.950 |
| *F13A1* | Source | +0.577 | [0.465, 0.817] | 100.0 | 8.600 | 13.882 |
| *MRC1* | Source | +0.354 | [0.176, 0.534] | 80.0 | 6.917 | 9.473 |
| *CFD* | Source | -0.800 | [-0.892, 0.305] | 40.0 | 18.191 | 10.315 |
| *CCL3* | Destination | +0.899 | [0.632, 4.877] | 80.0 | 0.936 | 3.256 |
| *CCL4* | Destination | +0.670 | [0.643, 3.082] | 80.0 | 0.826 | 1.940 |
| *IL1B* | Destination | +1.775 | [1.085, 2.568] | 100.0 | 0.674 | 2.656 |
| *CXCL8* | Destination | +1.535 | [1.367, 4.269] | 100.0 | 0.759 | 6.199 |
| **Scav → Fibro (21 libraries; 28 library–subtype strata)** | | | | | | |
| *C1QA* | Source | -0.095 | [-0.283, 0.027] | 38.1 | 37.431 | 32.879 |
| *C1QB* | Source | -0.153 | [-0.277, -0.011] | 23.8 | 36.322 | 30.778 |
| *APOE* | Source | +0.501 | [-0.497, 0.659] | 61.9 | 18.327 | 22.253 |
| *SELENOP* | Source | -0.495 | [-0.719, -0.361] | 14.3 | 23.372 | 16.790 |
| *SPP1* | Destination | +2.366 | [1.794, 3.221] | 95.2 | 5.646 | 21.627 |
| *FN1* | Destination | +1.109 | [0.258, 1.552] | 85.7 | 0.568 | 0.813 |
| *ANXA2* | Destination | +0.836 | [0.280, 0.904] | 90.5 | 4.009 | 6.034 |
| *LGALS1* | Destination | +0.877 | [0.613, 1.458] | 95.2 | 7.057 | 12.619 |

*Continued on next page*

| Gene | Panel role | Median $\log_2$FC | Library-effect IQR | Positive libraries (%) | Mean Q1 | Mean Q4 |
|---|---|---|---|---|---|---|
| **Res → Fibro (5 libraries; 5 library–subtype strata)** | | | | | | |
| *LYVE1* | Source | +0.575 | [-1.042, 0.685] | 60.0 | 9.249 | 12.328 |
| *F13A1* | Source | -0.040 | [-1.713, 0.530] | 40.0 | 11.297 | 11.821 |
| *MRC1* | Source | -0.661 | [-0.780, 0.255] | 40.0 | 9.042 | 7.253 |
| *CFD* | Source | +0.665 | [0.241, 1.091] | 100.0 | 11.236 | 20.234 |
| *SPP1* | Destination | +0.000 | [-0.805, 2.514] | 40.0 | 0.512 | 4.215 |
| *FN1* | Destination | +0.540 | [0.000, 0.577] | 60.0 | 5.299 | 8.332 |
| *ANXA2* | Destination | +1.043 | [0.960, 1.314] | 100.0 | 4.293 | 9.220 |
| *LGALS1* | Destination | +1.394 | [1.278, 1.880] | 100.0 | 4.611 | 13.554 |
| **Foam → Fibro (3 libraries; 3 library–subtype strata)** | | | | | | |
| *FABP5* | Source | +0.291 | [0.088, 0.674] | 66.7 | 25.794 | 33.447 |
| *PLIN2* | Source | -0.428 | [-0.434, -0.090] | 33.3 | 17.409 | 15.221 |
| *CSTB* | Source | +0.373 | [0.069, 1.162] | 66.7 | 23.466 | 36.443 |
| *LIPA* | Source | +0.675 | [0.423, 0.980] | 100.0 | 1.810 | 2.519 |
| *SPP1* | Destination | +0.453 | [0.423, 1.258] | 100.0 | 89.632 | 185.239 |
| *FN1* | Destination | -0.404 | [-0.416, 0.138] | 33.3 | 15.952 | 16.560 |
| *ANXA2* | Destination | +0.475 | [0.454, 0.617] | 100.0 | 6.461 | 9.831 |
| *LGALS1* | Destination | +0.576 | [0.394, 0.577] | 100.0 | 22.266 | 29.778 |
| **Inflam → Fibro (6 libraries; 6 library–subtype strata)** | | | | | | |
| *CCL3* | Source | -0.605 | [-0.645, -0.520] | 0.0 | 37.804 | 25.354 |
| *CCL4* | Source | -0.474 | [-0.695, -0.385] | 0.0 | 43.380 | 29.885 |
| *IL1B* | Source | +0.059 | [-0.107, 0.258] | 50.0 | 15.779 | 15.045 |
| *CXCL8* | Source | -0.474 | [-0.669, -0.119] | 16.7 | 13.945 | 11.141 |
| *SPP1* | Destination | +1.419 | [0.531, 2.010] | 83.3 | 4.170 | 9.154 |
| *FN1* | Destination | +1.784 | [1.460, 2.157] | 100.0 | 0.348 | 1.510 |
| *ANXA2* | Destination | +0.561 | [0.403, 0.899] | 83.3 | 2.049 | 3.054 |
| *LGALS1* | Destination | +0.828 | [0.370, 1.148] | 83.3 | 4.159 | 6.819 |

**Reporting.** IQR is the 25th–75th percentile range of library effects, not a confidence interval. Endpoint means separately average eligible subtypes within libraries and then libraries equally; their ratio need not reproduce the median $\log_2$FC. Panel role identifies the source or destination marker set, not exclusive expression or an expected sign. These are descriptive RNA effects; no historical pooled-cell significance tests or four-quartile trend tests are carried over. Protein, liver and held-out-gene validation results use their own documented source tables.

**Source data.** `table_6/Table_6.csv`, `table_6/Table_6_canonical.csv`, `table_6/per_library_gene_effects.csv`, `table_6/per_stratum_gene_effects.csv`, `table_6/coverage.csv`, `table_6/broad_signature_definitions.csv`.

#### Supplementary Table 7

##### Inferred transcription factor activity along OT association gradients.

Transcription factor (TF) activities were inferred from the current atlas using the univariate linear model (ULM) in `decoupler` 1.8.0 and the human CollecTRI network, with complexes retained. The supplied network contains 43,159 interactions and 1,186 regulator entries; 762 regulators passed the analysis requirements, including at least five available target genes. The full result comprises 8,382 regulator–route tests across 11 routes.

Activity differences are Q4 minus Q1 within eligible library–subtype strata (at least 20 cells per endpoint), averaged equally across subtypes within each library. Two-sided Wilcoxon signed-rank tests use library effects; Benjamini–Hochberg adjustment is applied jointly across all 8,382 tests. The displayed entries are the five lowest and five highest median effects per route, regardless of significance. Regulon scores represent inferred activity rather than TF transcript abundance or experimental evidence of regulation.

| Regulator | Median activity difference | Mean activity difference | Positive libraries (%) | $p$ | $q$ (BH) |
|---|---|---|---|---|---|
| **Mono → Scav (17 libraries; 22 strata)** | | | | | |
| *JDP2* | -1.180 | -1.028 | 0.0 | $1.53 \times 10^{-5}$ | $2.83 \times 10^{-4}$ |
| *ZNF335* | -1.029 | -0.900 | 0.0 | $1.53 \times 10^{-5}$ | $2.83 \times 10^{-4}$ |
| *HOXA7* | -0.921 | -0.911 | 5.9 | $3.05 \times 10^{-5}$ | $4.89 \times 10^{-4}$ |
| *MBD1* | -0.877 | -0.762 | 0.0 | $1.53 \times 10^{-5}$ | $2.83 \times 10^{-4}$ |
| *HIF1A* | -0.858 | -0.631 | 29.4 | 0.0267 | 0.0936 |
| *RFXAP* | +3.545 | +3.655 | 94.1 | $3.05 \times 10^{-5}$ | $4.89 \times 10^{-4}$ |
| *RFXANK* | +3.366 | +3.449 | 94.1 | $3.05 \times 10^{-5}$ | $4.89 \times 10^{-4}$ |
| *RFX5* | +3.099 | +3.014 | 94.1 | $3.05 \times 10^{-5}$ | $4.89 \times 10^{-4}$ |
| *CIITA* | +1.809 | +1.946 | 94.1 | $4.58 \times 10^{-5}$ | $6.53 \times 10^{-4}$ |
| *RFX1* | +1.248 | +1.276 | 94.1 | $3.05 \times 10^{-5}$ | $4.89 \times 10^{-4}$ |
| **Mono → Inflam (18 libraries; 23 strata)** | | | | | |
| *JDP2* | -0.979 | -0.858 | 11.1 | $3.81 \times 10^{-5}$ | $5.72 \times 10^{-4}$ |
| *HOXA7* | -0.785 | -0.751 | 5.6 | $2.29 \times 10^{-5}$ | $3.93 \times 10^{-4}$ |
| *FOSB* | -0.713 | -0.682 | 11.1 | $1.07 \times 10^{-4}$ | 0.0013 |
| *ZNF335* | -0.605 | -0.477 | 16.7 | $8.39 \times 10^{-4}$ | 0.0064 |
| *HDGF* | -0.585 | -0.504 | 5.6 | $8.39 \times 10^{-4}$ | 0.0064 |
| *RFXAP* | +1.711 | +1.789 | 88.9 | $3.81 \times 10^{-5}$ | $5.72 \times 10^{-4}$ |
| *RFXANK* | +1.661 | +1.693 | 88.9 | $3.81 \times 10^{-5}$ | $5.72 \times 10^{-4}$ |
| *RFX5* | +1.393 | +1.474 | 88.9 | $3.81 \times 10^{-5}$ | $5.72 \times 10^{-4}$ |
| *DMTF1* | +1.273 | +1.090 | 100.0 | $7.63 \times 10^{-6}$ | $1.72 \times 10^{-4}$ |
| *REL* | +1.132 | +1.455 | 88.9 | 0.0010 | 0.0074 |
| **Mono → Foam (10 libraries; 16 strata)** | | | | | |
| *PAWR* | -0.903 | -1.062 | 0.0 | 0.0020 | 0.0119 |
| *HIPK2* | -0.665 | -0.828 | 0.0 | 0.0020 | 0.0119 |
| *TET1* | -0.603 | -0.555 | 10.0 | 0.0195 | 0.0721 |
| *KLF17* | -0.594 | -0.476 | 10.0 | 0.0059 | 0.0277 |
| *NFKBIB* | -0.593 | -0.527 | 20.0 | 0.0137 | 0.0539 |

*Continued on next page*

| Regulator | Median activity difference | Mean activity difference | Positive libraries (%) | $p$ | $q$ (BH) |
|---|---|---|---|---|---|
| *MYC* | +2.359 | +2.484 | 90.0 | 0.0039 | 0.0201 |
| *SP1* | +1.470 | +1.396 | 90.0 | 0.0039 | 0.0201 |
| *TP53* | +1.015 | +1.019 | 90.0 | 0.0039 | 0.0201 |
| *HOXA7* | +0.992 | +0.933 | 100.0 | 0.0020 | 0.0119 |
| *JUN* | +0.974 | +0.966 | 90.0 | 0.0098 | 0.0414 |
| **Mono $\rightarrow$ Res (15 libraries; 17 strata)** | | | | | |
| *NFKB1* | -1.376 | -1.099 | 26.7 | 0.0054 | 0.0262 |
| *DMTF1* | -1.343 | -0.952 | 26.7 | 0.0034 | 0.0183 |
| *RELA* | -1.217 | -0.944 | 26.7 | 0.0125 | 0.0502 |
| *JUN* | -0.976 | -0.739 | 33.3 | 0.0730 | 0.1640 |
| *REL* | -0.931 | -0.916 | 20.0 | 0.0103 | 0.0430 |
| *ING4* | +0.806 | +0.755 | 100.0 | $6.10 \times 10^{-5}$ | $8.03 \times 10^{-4}$ |
| *LYL1* | +0.566 | +0.580 | 100.0 | $6.10 \times 10^{-5}$ | $8.03 \times 10^{-4}$ |
| *JDP2* | +0.523 | +0.250 | 73.3 | 0.1354 | 0.2623 |
| *PAWR* | +0.515 | +0.249 | 73.3 | 0.2078 | 0.3615 |
| *SIM2* | +0.504 | +0.530 | 100.0 | $6.10 \times 10^{-5}$ | $8.03 \times 10^{-4}$ |
| **Mono $\rightarrow$ Fibro (12 libraries; 14 strata)** | | | | | |
| *RFXANK* | -1.573 | -1.353 | 8.3 | 0.0049 | 0.0241 |
| *RFXAP* | -1.539 | -1.405 | 16.7 | 0.0093 | 0.0404 |
| *RFX5* | -1.413 | -1.127 | 16.7 | 0.0093 | 0.0404 |
| *REL* | -1.263 | -0.967 | 16.7 | 0.0269 | 0.0936 |
| *PAWR* | -1.063 | -1.235 | 0.0 | $4.88 \times 10^{-4}$ | 0.0042 |
| *MYC* | +2.009 | +2.647 | 100.0 | $4.88 \times 10^{-4}$ | 0.0042 |
| *SP1* | +1.319 | +1.395 | 91.7 | $9.77 \times 10^{-4}$ | 0.0071 |
| *ZBTB4* | +1.262 | +1.702 | 100.0 | $4.88 \times 10^{-4}$ | 0.0042 |
| *HOXA7* | +1.078 | +1.128 | 100.0 | $4.88 \times 10^{-4}$ | 0.0042 |
| *ZNF335* | +1.046 | +1.210 | 100.0 | $4.88 \times 10^{-4}$ | 0.0042 |
| **Scav $\rightarrow$ Inflam (26 libraries; 33 strata)** | | | | | |
| *JDP2* | -1.221 | -1.185 | 0.0 | $2.98 \times 10^{-8}$ | $2.75 \times 10^{-6}$ |
| *DACH1* | -1.064 | -1.024 | 3.8 | $5.96 \times 10^{-8}$ | $4.34 \times 10^{-6}$ |
| *ZNF382* | -0.943 | -0.935 | 3.8 | $5.96 \times 10^{-8}$ | $4.34 \times 10^{-6}$ |
| *NAB2* | -0.849 | -0.796 | 7.7 | $1.49 \times 10^{-7}$ | $8.55 \times 10^{-6}$ |
| *HOXA7* | -0.833 | -0.850 | 7.7 | $4.17 \times 10^{-7}$ | $1.83 \times 10^{-5}$ |
| *RFXAP* | +3.228 | +3.234 | 100.0 | $2.98 \times 10^{-8}$ | $2.75 \times 10^{-6}$ |
| *RFXANK* | +3.063 | +3.068 | 100.0 | $2.98 \times 10^{-8}$ | $2.75 \times 10^{-6}$ |
| *RFX5* | +2.630 | +2.725 | 100.0 | $2.98 \times 10^{-8}$ | $2.75 \times 10^{-6}$ |
| *CIITA* | +2.585 | +2.538 | 100.0 | $2.98 \times 10^{-8}$ | $2.75 \times 10^{-6}$ |
| *REL* | +2.151 | +2.532 | 96.2 | $5.96 \times 10^{-8}$ | $4.34 \times 10^{-6}$ |
| **Res $\rightarrow$ Inflam (5 libraries; 5 strata)** | | | | | |
| *HOXA7* | -1.764 | -1.765 | 0.0 | 0.0625 | 0.1447 |

*Continued on next page*

| Regulator | Median activity difference | Mean activity difference | Positive libraries (%) | $p$ | $q$ (BH) |
|---|---|---|---|---|---|
| *JDP2* | -1.722 | -1.738 | 0.0 | 0.0625 | 0.1447 |
| *ZNF335* | -1.623 | -1.405 | 0.0 | 0.0625 | 0.1447 |
| *DACH1* | -1.591 | -1.574 | 0.0 | 0.0625 | 0.1447 |
| *ZNF382* | -1.441 | -1.275 | 0.0 | 0.0625 | 0.1447 |
| *RFXAP* | +3.718 | +3.389 | 100.0 | 0.0625 | 0.1447 |
| *CIITA* | +3.537 | +3.164 | 100.0 | 0.0625 | 0.1447 |
| *RFXANK* | +3.466 | +3.171 | 100.0 | 0.0625 | 0.1447 |
| *RFX5* | +3.166 | +2.779 | 100.0 | 0.0625 | 0.1447 |
| *REL* | +3.050 | +3.088 | 100.0 | 0.0625 | 0.1447 |
| **Scav $\rightarrow$ Fibro (21 libraries; 28 strata)** | | | | | |
| *REL* | -1.545 | -1.603 | 4.8 | $5.25 \times 10^{-5}$ | $7.27 \times 10^{-4}$ |
| *RELA* | -1.351 | -1.178 | 4.8 | $4.10 \times 10^{-5}$ | $6.01 \times 10^{-4}$ |
| *NFKB1* | -1.276 | -1.196 | 9.5 | $1.05 \times 10^{-4}$ | 0.0013 |
| *TBX21* | -1.243 | -1.219 | 0.0 | $9.54 \times 10^{-7}$ | $3.27 \times 10^{-5}$ |
| *IRF1* | -1.121 | -0.932 | 4.8 | $3.15 \times 10^{-5}$ | $4.93 \times 10^{-4}$ |
| *MYC* | +1.600 | +2.053 | 95.2 | $1.91 \times 10^{-6}$ | $5.53 \times 10^{-5}$ |
| *ZBTB4* | +1.586 | +1.551 | 100.0 | $9.54 \times 10^{-7}$ | $3.27 \times 10^{-5}$ |
| *ZBED1* | +1.057 | +0.894 | 100.0 | $9.54 \times 10^{-7}$ | $3.27 \times 10^{-5}$ |
| *HOXA7* | +1.007 | +1.016 | 100.0 | $9.54 \times 10^{-7}$ | $3.27 \times 10^{-5}$ |
| *JDP2* | +0.766 | +0.799 | 100.0 | $9.54 \times 10^{-7}$ | $3.27 \times 10^{-5}$ |
| **Res $\rightarrow$ Fibro (5 libraries; 5 strata)** | | | | | |
| *HBP1* | -1.944 | -1.522 | 0.0 | 0.0625 | 0.1447 |
| *DMTF1* | -1.586 | -1.294 | 0.0 | 0.0625 | 0.1447 |
| *OVOL1* | -1.463 | -1.130 | 20.0 | 0.1250 | 0.2476 |
| *PAWR* | -1.442 | -1.307 | 0.0 | 0.0625 | 0.1447 |
| *HOXB9* | -1.328 | -1.074 | 0.0 | 0.0625 | 0.1447 |
| *JDP2* | +1.610 | +1.625 | 100.0 | 0.0625 | 0.1447 |
| *HOXA7* | +1.376 | +1.443 | 100.0 | 0.0625 | 0.1447 |
| *ZNF335* | +1.127 | +1.101 | 100.0 | 0.0625 | 0.1447 |
| *MYB* | +1.069 | +1.154 | 100.0 | 0.0625 | 0.1447 |
| *ZBTB4* | +1.052 | +1.616 | 100.0 | 0.0625 | 0.1447 |
| **Foam $\rightarrow$ Fibro (3 libraries; 3 strata)** | | | | | |
| *CIITA* | -1.658 | -1.245 | 33.3 | 0.5000 | 0.6423 |
| *IRF1* | -1.346 | -1.096 | 0.0 | 0.2500 | 0.3979 |
| *IRF3* | -1.202 | -1.052 | 0.0 | 0.2500 | 0.3979 |
| *ZFX* | -1.188 | -0.877 | 0.0 | 0.2500 | 0.3979 |
| *BRCA1* | -1.149 | -1.010 | 0.0 | 0.2500 | 0.3979 |
| *MYC* | +3.161 | +3.264 | 100.0 | 0.2500 | 0.3979 |
| *ZBTB4* | +1.271 | +1.772 | 100.0 | 0.2500 | 0.3979 |
| *HOPX* | +1.091 | +1.113 | 100.0 | 0.2500 | 0.3979 |

*Continued on next page*

| Regulator | Median activity difference | Mean activity difference | Positive libraries (%) | $p$ | $q$ (BH) |
|---|---|---|---|---|---|
| *ZBED1* | +1.043 | +0.935 | 66.7 | 0.5000 | 0.6423 |
| *ZNF335* | +0.981 | +0.620 | 66.7 | 0.5000 | 0.6423 |
| **Inflam $\rightarrow$ Fibro (6 libraries; 6 strata)** | | | | | |
| *ZNF202* | -1.043 | -0.921 | 16.7 | 0.0938 | 0.2006 |
| *PAWR* | -0.958 | -1.027 | 0.0 | 0.0312 | 0.1008 |
| *HIPK2* | -0.934 | -0.912 | 0.0 | 0.0312 | 0.1008 |
| *MTA2* | -0.811 | -0.854 | 0.0 | 0.0312 | 0.1008 |
| *CIITA* | -0.799 | -0.923 | 0.0 | 0.0312 | 0.1008 |
| *MYC* | +1.595 | +1.514 | 100.0 | 0.0312 | 0.1008 |
| *MYB* | +1.117 | +1.031 | 100.0 | 0.0312 | 0.1008 |
| *HIF1A* | +1.086 | +1.195 | 100.0 | 0.0312 | 0.1008 |
| *SP1* | +1.009 | +1.053 | 100.0 | 0.0312 | 0.1008 |
| *TP53* | +0.962 | +0.860 | 100.0 | 0.0312 | 0.1008 |

**Inference.** Libraries are the analysis units and are not necessarily independent patients. The limited number of eligible libraries constrains inference for several routes. Effect-ranked rows are retained even when $q \geq 0.05$; no individual regulators pass $q < 0.05$ in the current Res–Inflam, Res–Fibro, Foam–Fibro or Inflam–Fibro comparisons. The complete results and per-library effects are provided as source data.

**Source data.** `table_7/Table_7.csv`, `table_7/Table_7_display_tails.csv`, `table_7/per_library_TF_effects.csv`, `table_7/coverage.csv`, `table_7/network_used.csv`.

#### Supplementary Table 8

**Bashore patient metadata and macrophage composition.**

Clinical metadata and current-atlas cell counts are reported for all 18 mapped Bashore patients (11 symptomatic, seven asymptomatic), using the documented BioSample-to-patient mapping. Counts total 16,336 retained monocytes/macrophages. ADT coverage refers to the 3,707 cells whose identifiers occur in both the current RNA atlas and the saved ADT object, across six patients; these are a subset of the RNA cells and are not added to the RNA total.

**A. Patient metadata and modality coverage.**

| Patient | BioSample | Age (yr) | Sex | Clinical status | RNA cells | RNA libraries | RNA cells with ADT |
|---|---|---|---|---|---|---|---|
| P_46M_S | SAMN39546314 | 46 | M | Symptomatic | 2,068 | 1 | 0 |
| P_59M_S | SAMN39546307 | 59 | M | Symptomatic | 830 | 1 | 0 |
| P_64M_S | SAMN39546316 | 64 | M | Symptomatic | 1,685 | 1 | 0 |
| P_66F_S | SAMN39546315 | 66 | F | Symptomatic | 97 | 1 | 0 |
| P_66M_S | SAMN39546310 | 66 | M | Symptomatic | 534 | 1 | 0 |
| P_75M_S | SAMN39550058 | 75 | M | Symptomatic | 361 | 1 | 360 |
| P_78F_S | SAMN39550056 | 78 | F | Symptomatic | 860 | 1 | 855 |
| P_79M_S | SAMN39550062 | 79 | M | Symptomatic | 294 | 1 | 290 |
| P_80M_S | SAMN39550054 | 80 | M | Symptomatic | 1,060 | 1 | 1,025 |
| P_81M_S | SAMN39550052 | 81 | M | Symptomatic | 1,056 | 1 | 1,055 |
| P_89M_S | SAMN39546318 | 89 | M | Symptomatic | 221 | 1 | 0 |
| P_58M_A | SAMN39546308 | 58 | M | Asymptomatic | 394 | 1 | 0 |
| P_62M_A | SAMN39546312 | 62 | M | Asymptomatic | 1,728 | 1 | 0 |
| P_64M_A | SAMN39546311 | 64 | M | Asymptomatic | 1,250 | 1 | 0 |
| P_67M_A | SAMN39546317 | 67 | M | Asymptomatic | 1,961 | 1 | 0 |
| P_71M_A | SAMN39550060 | 71 | M | Asymptomatic | 154 | 1 | 122 |
| P_75M_A | SAMN39546309 | 75 | M | Asymptomatic | 1,049 | 1 | 0 |
| P_76F_A | SAMN39546313 | 76 | F | Asymptomatic | 734 | 1 | 0 |

**B. Meta-cluster composition: cell count (percentage of patient RNA cells).**

| Patient | Mono | Scav | Res | Inflam | Foam | Fibro |
|---|---|---|---|---|---|---|
| P_46M_S | 358 (17.3%) | 801 (38.7%) | 2 (0.1%) | 279 (13.5%) | 275 (13.3%) | 353 (17.1%) |
| P_59M_S | 218 (26.3%) | 428 (51.6%) | 2 (0.2%) | 16 (1.9%) | 11 (1.3%) | 155 (18.7%) |
| P_64M_S | 404 (24.0%) | 642 (38.1%) | 0 (0.0%) | 352 (20.9%) | 73 (4.3%) | 214 (12.7%) |
| P_66F_S | 47 (48.5%) | 26 (26.8%) | 0 (0.0%) | 14 (14.4%) | 7 (7.2%) | 3 (3.1%) |
| P_66M_S | 100 (18.7%) | 350 (65.5%) | 0 (0.0%) | 18 (3.4%) | 5 (0.9%) | 61 (11.4%) |
| P_75M_S | 148 (41.0%) | 144 (39.9%) | 0 (0.0%) | 63 (17.5%) | 6 (1.7%) | 0 (0.0%) |
| P_78F_S | 315 (36.6%) | 460 (53.5%) | 0 (0.0%) | 17 (2.0%) | 29 (3.4%) | 39 (4.5%) |
| P_79M_S | 33 (11.2%) | 116 (39.5%) | 5 (1.7%) | 25 (8.5%) | 11 (3.7%) | 104 (35.4%) |
| P_80M_S | 275 (25.9%) | 519 (49.0%) | 3 (0.3%) | 60 (5.7%) | 65 (6.1%) | 138 (13.0%) |
| P_81M_S | 96 (9.1%) | 554 (52.5%) | 0 (0.0%) | 319 (30.2%) | 9 (0.9%) | 78 (7.4%) |
| P_89M_S | 18 (8.1%) | 191 (86.4%) | 0 (0.0%) | 8 (3.6%) | 0 (0.0%) | 4 (1.8%) |
| P_58M_A | 10 (2.5%) | 343 (87.1%) | 3 (0.8%) | 7 (1.8%) | 15 (3.8%) | 16 (4.1%) |
| P_62M_A | 117 (6.8%) | 1,023 (59.2%) | 7 (0.4%) | 167 (9.7%) | 27 (1.6%) | 387 (22.4%) |
| P_64M_A | 214 (17.1%) | 840 (67.2%) | 0 (0.0%) | 14 (1.1%) | 8 (0.6%) | 174 (13.9%) |
| P_67M_A | 312 (15.9%) | 408 (20.8%) | 0 (0.0%) | 21 (1.1%) | 27 (1.4%) | 1,193 (60.8%) |
| P_71M_A | 17 (11.0%) | 29 (18.8%) | 0 (0.0%) | 0 (0.0%) | 11 (7.1%) | 97 (63.0%) |
| P_75M_A | 94 (9.0%) | 585 (55.8%) | 0 (0.0%) | 74 (7.1%) | 139 (13.3%) | 157 (15.0%) |
| P_76F_A | 231 (31.5%) | 265 (36.1%) | 0 (0.0%) | 210 (28.6%) | 5 (0.7%) | 23 (3.1%) |

M, male; F, female. Library counts refer to unique RNA libraries represented in the atlas, not the number of sequencing runs. Zero ADT overlap indicates no matched cells in the supplied ADT object; it does not denote zero protein expression. Meta-cluster proportions use the retained RNA cells of each patient as their denominator. Percentages may not sum to exactly 100 after rounding. Clinical associations and their tests are reported separately in the main analysis.

**Source data.** `table_8/Table_8.csv`, `table_8/library_patient_mapping.csv`, `table_8/atlas_ADT_coverage.csv`.

#### Supplementary Table 9

##### Quality control settings and cell accounting.

The recorded QC summaries contain 62 input libraries and 242,433 cells. Fifty libraries retain 201,858 cells across all lineages; downstream selection and annotation yield the 34,533-cell monocyte/macrophage atlas. The difference between all-lineage post-QC counts and final atlas counts includes lineage selection and subsequent exclusions, rather than an additional single QC filter.

**A. Cohort-level cell accounting and depth.**

| Cohort | Input libraries | Retained libraries | Before QC cells | Post-QC cells (all lineages) | Final atlas cells | Median UMI / atlas cell | Median unspliced UMI / atlas cell |
|---|---|---|---|---|---|---|---|
| Alsaigh | 6 | 6 | 54,906 | 44,009 | 7,619 | 10889.0 | 2002.0 |
| Bashore | 18 | 18 | 94,062 | 82,061 | 16,336 | 11596.0 | 2622.0 |
| Fernandez | 6 | 3 | 10,782 | 8,201 | 1,350 | 4858.0 | 888.0 |
| Jaiswal | 9 | 9 | 38,601 | 32,550 | 4,488 | 12197.0 | 4661.5 |
| Pan | 3 | 3 | 12,270 | 10,473 | 2,037 | 12702.0 | 3102.0 |
| Pauli | 12 | 3 | 10,192 | 4,952 | 319 | 3639.0 | 699.0 |
| Wirka | 8 | 8 | 21,620 | 19,612 | 2,384 | 4633.0 | 775.5 |
| **Total** | 62 | 50 | 242,433 | 201,858 | 34,533 | — | — |

Median depth values are calculated from the current atlas's integer-like `counts` and `unspliced` layers, respectively, rather than from all cells entering QC. Input counts refer to the preprocessing stage recorded for each library. Of 62 inputs, 58 are labelled CellBender-corrected and four are labelled standard CellRanger following CellBender failure. All 50 libraries represented in the final atlas are labelled CellBender-corrected.

**B. Executed QC parameters (common to all seven cohorts).**

| Parameter | Value | Application |
|---|---|---|
| MAD multiplier | 5 | Lower-tail outlier filtering of log counts and detected genes; upper-tail mitochondrial-fraction filtering. |
| Mitochondrial cap | 15% | Additional hard upper bound on mitochondrial transcript percentage. |
| Minimum library size | 1,000 cells | Whole-library exclusion below the input cell-count threshold. |
| Minimum median detected genes | 500 | Whole-library exclusion when the post-filter median is below this threshold. |
| Doublet-rate parameter | 0.008 per 1,000 cells | Expected-rate, score-ranked SOLO removal; not a universal SOLO-score threshold or fixed fraction for every library. |

Settings were recovered from completed execution logs whose cohort-level input and output counts match the current QC summaries. Different QC flags can overlap, so per-filter counts must not be summed as disjoint exclusions.

**C. Libraries excluded at the recorded QC stage.**

| Cohort | Library | Input cells | Recorded exclusion |
|---|---|---|---|
| Fernandez | fernandez_GSM7018579 | 191 | Input library below 1,000 cells |
| Fernandez | fernandez_GSM7018581 | 905 | Input library below 1,000 cells |
| Fernandez | fernandez_GSM7018582 | 232 | Input library below 1,000 cells |
| Pauli | pauli_GSM7885766 | 263 | Input library below 1,000 cells |
| Pauli | pauli_GSM7885769 | 232 | Input library below 1,000 cells |
| Pauli | pauli_GSM7885772 | 622 | Input library below 1,000 cells |
| Pauli | pauli_GSM7885773 | 611 | Input library below 1,000 cells |
| Pauli | pauli_GSM7885774 | 1,382 | Post-filter median detected genes 466 (below 500) |
| Pauli | pauli_GSM7885775 | 384 | Input library below 1,000 cells |
| Pauli | pauli_GSM7885777 | 231 | Input library below 1,000 cells |
| Pauli | pauli_GSM7885778 | 168 | Input library below 1,000 cells |
| Pauli | pauli_GSM7885780 | 978 | Input library below 1,000 cells |

Eleven libraries fail the input-size threshold and one fails the post-filter median-gene threshold. Every library with cells retained by these QC summaries contributes cells to the final atlas. Per-library counts, preprocessing stage, overlapping removal flags and log provenance are retained in the source data.

**Source data.** `table_9/Table_9.csv`, `table_9/per_library_QC.csv`, `table_9/QC_removed_libraries.csv`, `table_9/QC_settings_log_evidence.csv`.

#### Supplementary Figures

**Supplementary Figure 1 - Multi-cohort integration: QC and composition balance**

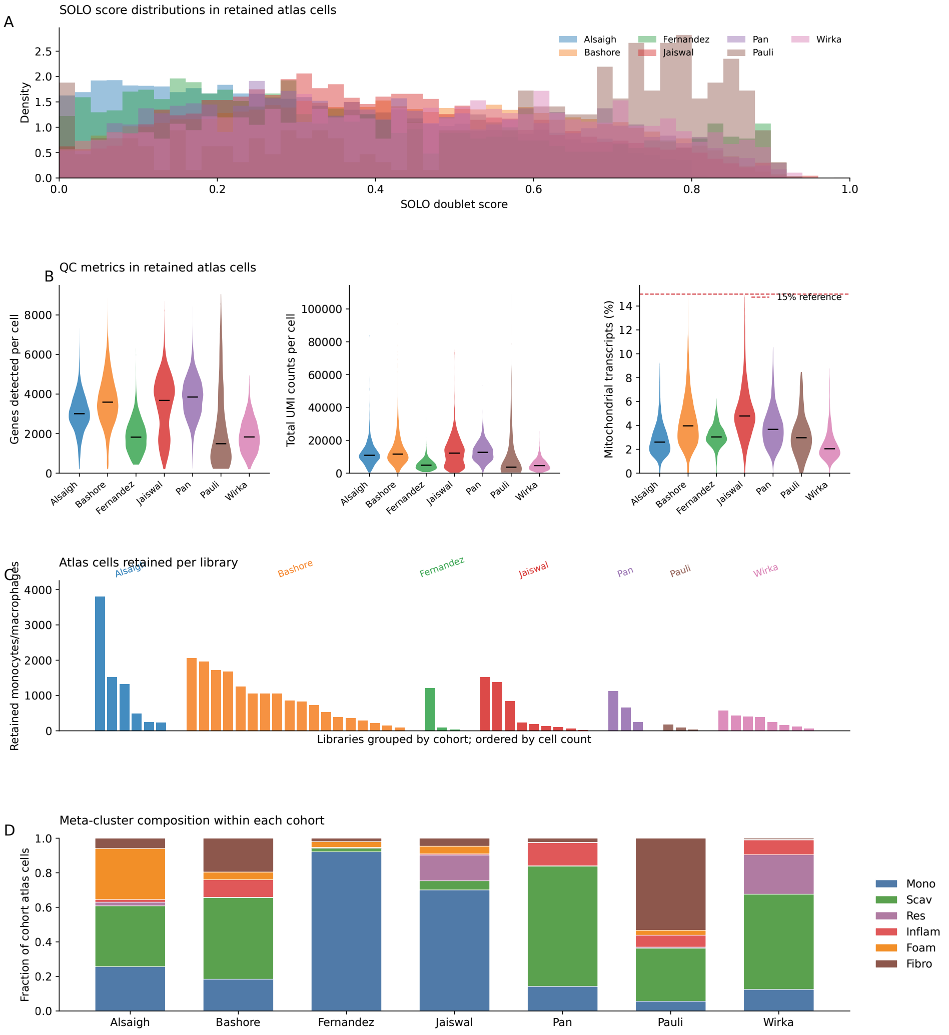


**Supplementary Figure 1 — Quality control and cohort composition of the current atlas. (A)** Saved SOLO doublet-score distributions among retained cells, grouped by cohort. SOLO was trained per library and filtering removed the highest-scoring cells according to the library-specific expected doublet rate; there was no universal 0.5 score cutoff. **(B)** Distributions of detected genes, total UMI counts and mitochondrial percentage among retained cells, grouped by cohort. **(C)** Number of final-atlas cells per library, grouped by cohort. **(D)** Fractions of the six meta-clusters within each cohort. The figure describes 34,533 retained cells across 50 libraries and seven cohorts; executed QC settings and excluded libraries are reported in Supplementary Table 9.

**Supplementary figure 2 - Leiden resolution selection**

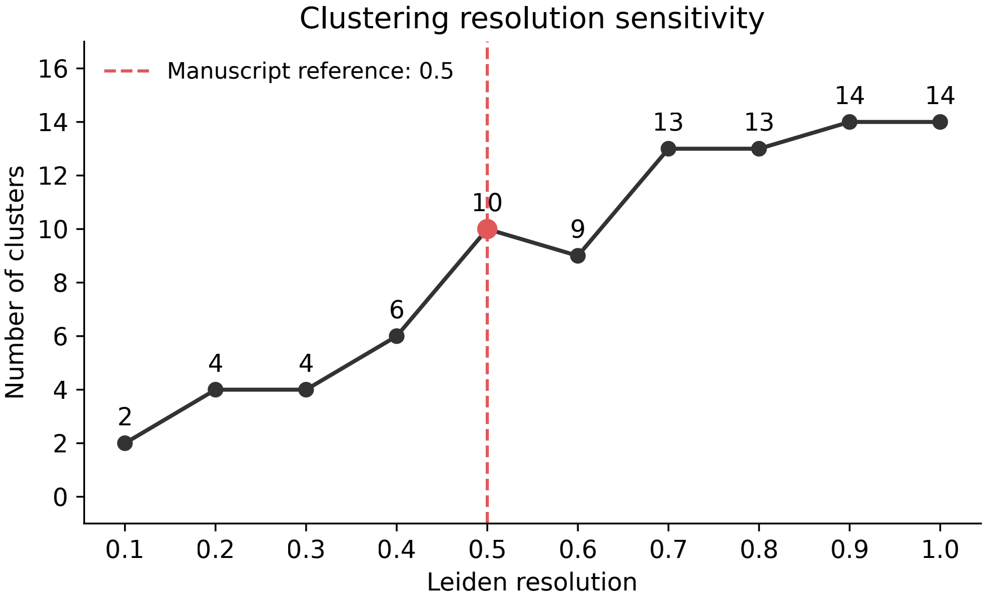


**Supplementary Figure 2 — Sensitivity of cluster number to Leiden resolution.** Number of clusters obtained over resolutions 0.1–1.0 on the saved Scanorama neighbor graph of the current atlas. This diagnostic recomputes partitions on the current cell set without changing the saved biological annotations. Resolution 0.5 is highlighted as a reference point on this diagnostic; the final annotation uses stored resolution-0.8 assignments with additional subclustering (Supplementary Table 3). Cluster counts alone do not measure membership stability.

**Supplementary figure 3 - Statistical robustness of Optimal Transport inference**

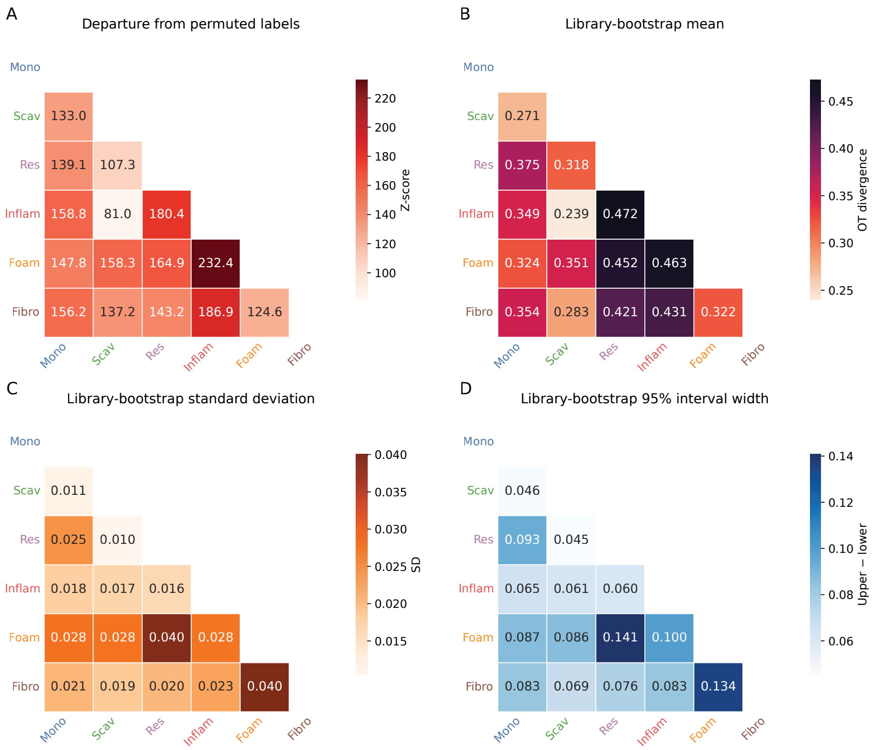


**Supplementary Figure 3 — Label-null separation and library-resampling variability of pairwise OT divergence.** All 15 unordered meta-cluster pairs are shown. **(A)** Standardized deviation from the 50-permutation label-mixing null ($z = 81.02$–232.41). **(B)** Mean OT divergence across library-bootstrap resamples. **(C)** Library-bootstrap standard deviation. **(D)** Width of the 2.5th–97.5th percentile library-bootstrap interval. The library analysis requested 200 resamples, with a maximum of 2,000 cells per state and exclusion of resamples with insufficient state coverage. Standard deviations range from 0.0104 to 0.0400 and interval widths from 0.0452 to 0.1407. These distributions are distinct from the reference calculation using up to 5,000 cells per state. Larger divergence denotes greater transcriptional separation; null $z$-scores are not tests of biological interconversion. Full values and reference estimates are given in Supplementary Table 4.

**Supplementary figure 4 - Leave-One-Cohort-Out (LOCO) statistical robustness**

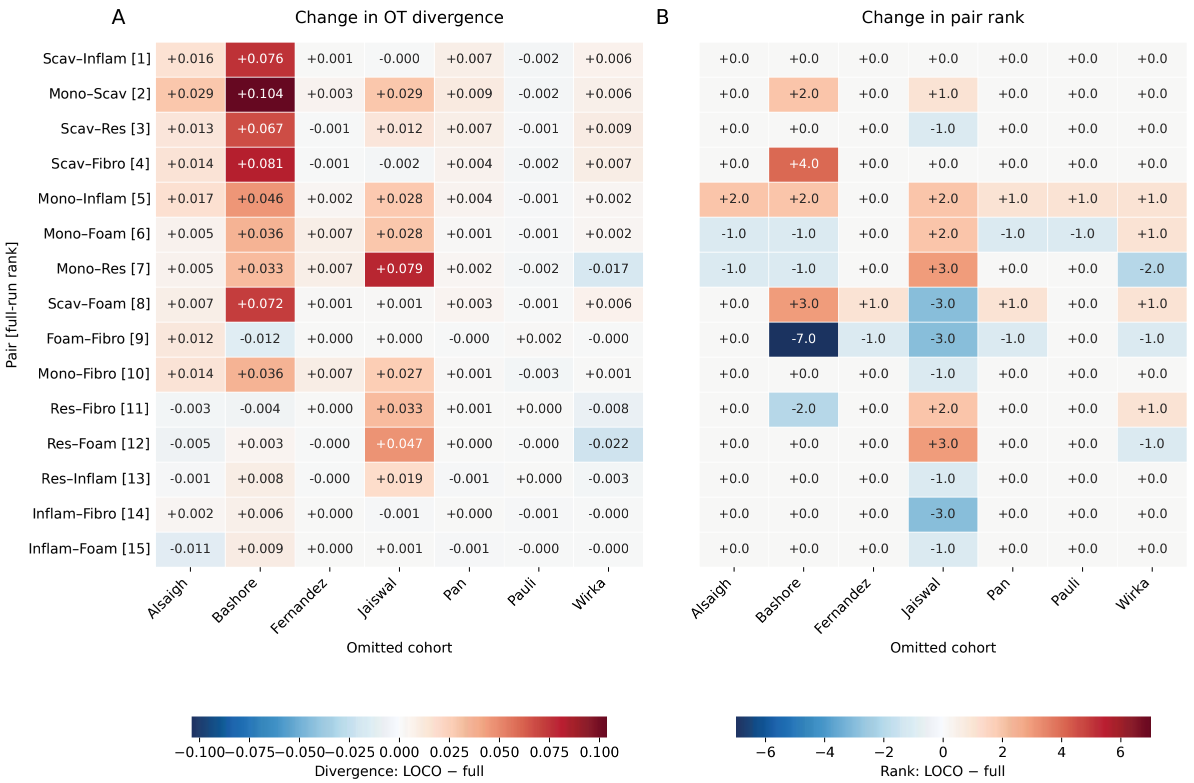


**Supplementary Figure 4 — Sensitivity of OT relationships to omission of individual cohorts.** Each column omits one of the seven cohorts from the existing integrated representation; integration itself was not refitted. Rows show the 15 pairs ordered by full-cohort reference divergence. **(A)** Signed divergence change, LOCO minus full-cohort reference; positive values indicate greater separation after omission. **(B)** Signed rank change using ascending divergence (rank 1, closest); positive values indicate a shift towards a more distant rank. Rank correlations with the full result are 0.989, 0.843, 0.996, 0.889, 0.993, 0.996 and 0.982 for omission of Alsaigh, Bashore, Fernandez, Jaiswal, Pan, Pauli and Wirka, respectively. Scav–Inflam remains the closest pair in every run. The full 105 pair-by-omission estimates are retained in Supplementary Table 4.

**Supplementary figure 5 - UniTVelo RNA velocity stream plots for all pairwise transitions**

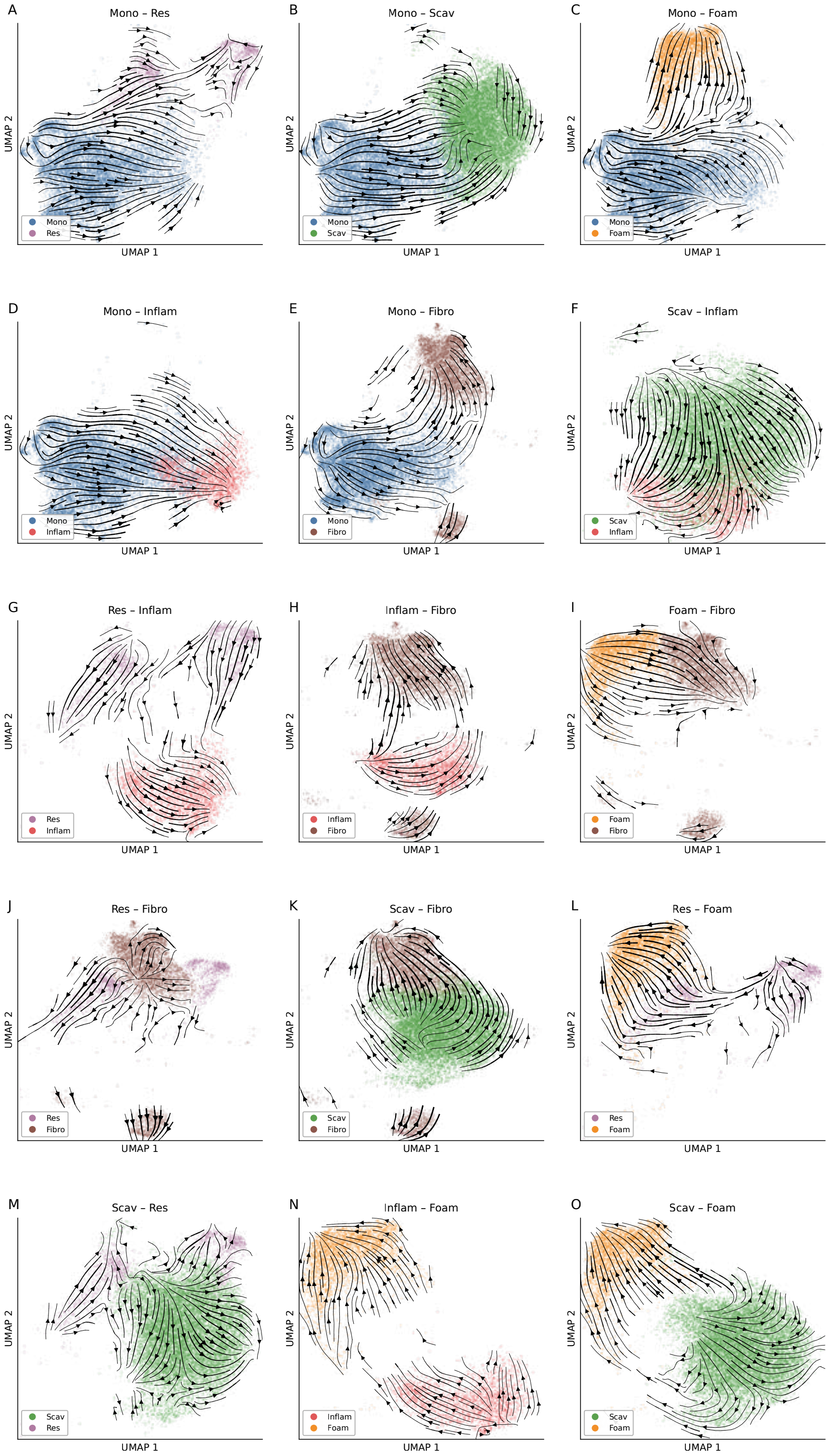

**Supplementary Figure 5 — Pairwise UniTVelo velocity stream displays under the designated source-root initialization.** The 15 panels show the corresponding two-state subsets, colored by meta-cluster, with projected velocities displayed as streamlines on the UMAP coordinates. Panel titles identify the pair and supplied root. Saved fitted models and velocity graphs were reused for visualization; plotting does not constitute a new fit. Quantitative orientation is assessed by the transition-support asymmetry in Supplementary Table 5, not by visual inspection, a UMAP centroid cosine, or an outlined convex-hull interface. Root and unspecified-root sensitivity results are reported alongside the primary estimates in that table. Streamlines depict conditional model output and do not directly track individual cells through time.

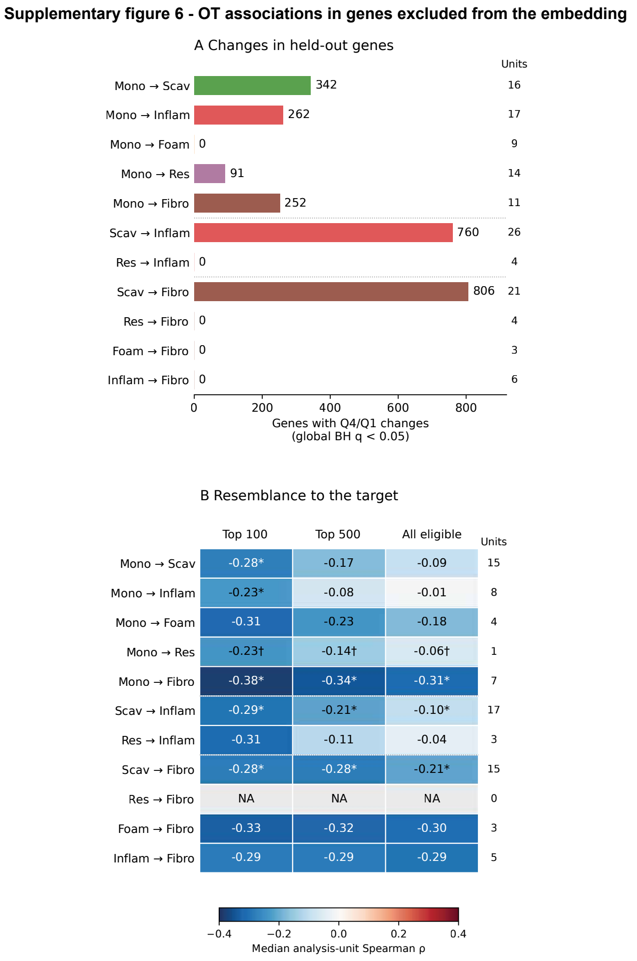


**Supplementary Figure 6 — OT associations in genes excluded from the integration features.** Genes excluded from the OT embedding provide a held-out-feature check of target-associated expression changes and target resemblance, reducing dependence on the genes used to construct the representation. Held-out genes were not selected as highly variable in any cohort and were absent from the final integration feature set. **(A)** Number of held-out genes with Q4/Q1 changes at global Benjamini–Hochberg $q < 0.05$ across 42,415 gene–route tests (2,513 discoveries, including positive and negative effects). Genes required detection in at least 20 and 1% of source cells. RNA effects use linearized normalized expression, pseudocount 0.1, at least 20 cells per endpoint per library–subtype stratum, and equal subtype weighting within libraries. Effects from eligible known paired preparations were then averaged into one analysis unit; other libraries remained separate. Bar colors denote targets; analysis-unit counts are shown at right. **(B)** Median analysis-unit Spearman correlation between corrected OT association and cosine distance to a same-library target centroid in log-expression space. Columns use the top 100, top 500 or all eligible variable held-out genes, ranked without reference to OT scores. References require at least 20 target cells; strata require at least 20 valid source cells per endpoint. Within-stratum correlations were averaged equally within libraries and then within known paired preparations. Negative values indicate closer target resemblance at higher association. Asterisks denote $q < 0.05$ across route–panel tests, a separate testing family from panel A; tests assess mean analysis-unit effects or correlations, whereas panel B displays medians. Two-sided sign flips are exact for 3–12 units and use 9,999 Monte Carlo patterns with a plus-one correction above that range. Daggers mark descriptive estimates based on fewer than three units; NA indicates no eligible comparison. These are held-out-feature controls in the same cells.

**Supplementary figure 7 - SOLO doublet scores across OT commitment quartiles for the three routes to inflammatory activation**

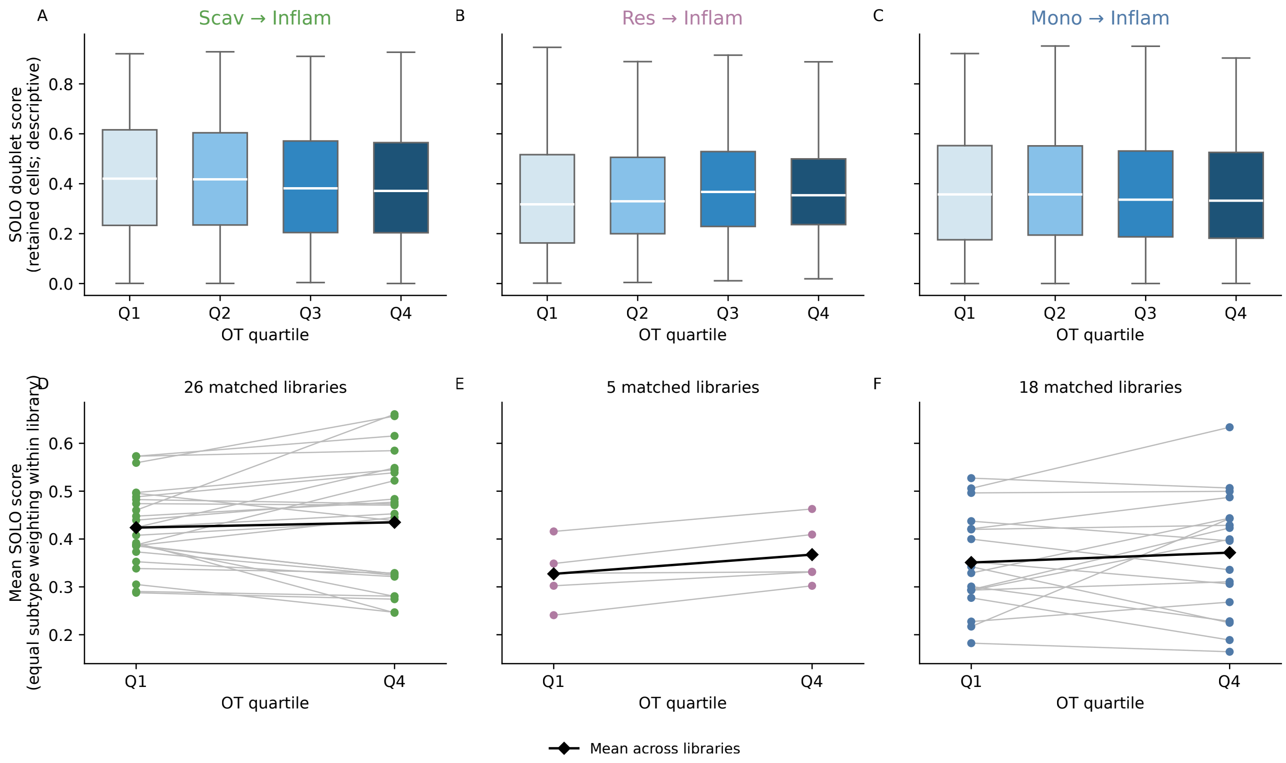


**Supplementary Figure 7 — Saved SOLO scores across inflammatory OT quartiles. (A–C)** Distributions among retained source cells for Scav–Inflam, Res–Inflam and Mono–Inflam. Boxes show the median and interquartile range; whiskers reach the most extreme values within 1.5 interquartile ranges. Outliers omitted from display remain in calculations. These pooled distributions are descriptive. **(D–F)** Matched Q1 and Q4 mean scores in 26, five and 18 libraries, respectively. Means use equal weighting of eligible source subtypes within each library, requiring at least 20 Q1 and 20 Q4 cells per subtype. Gray lines connect each library; black diamonds and connecting lines show arithmetic means across libraries. These are paired library summaries, not confidence intervals. For statistical inference, known paired preparations were averaged into analysis units, yielding 26, four and 17 units for Scav–Inflam, Res–Inflam and Mono–Inflam, respectively. Mean differences, bootstrap intervals and sign-flip tests use these units, as described in the Supplementary Methods. The black diamonds retain the descriptive means across individual libraries and need not equal the grouped estimates. The plot evaluates score variation among retained cells and does not establish that residual doublets are absent.

**Supplementary Figure 8 - Marker gene expression across 9 sub-clusters before meta-clustering**

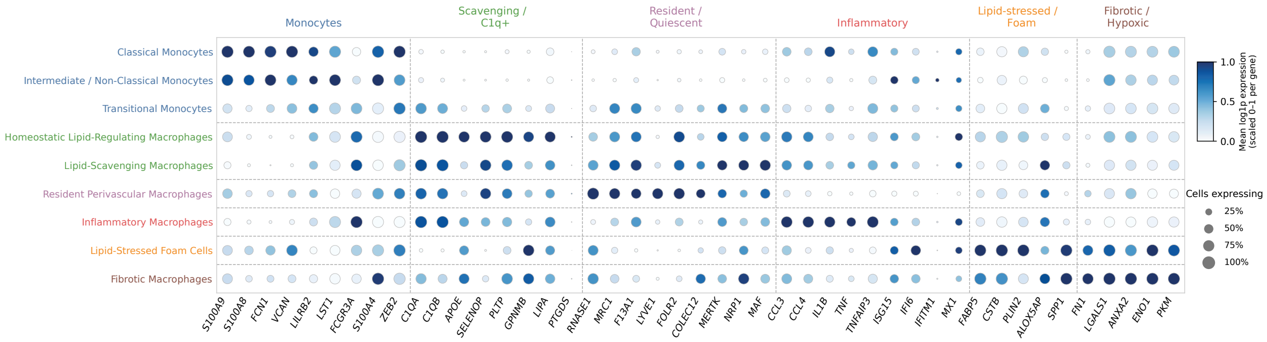


**Supplementary Figure 8 — Canonical marker expression across the nine annotated subtypes.** Dot area represents the fraction of cells with expression above zero; color represents mean log-normalized expression, min–max scaled from zero to one for each gene across subtypes. These are pooled descriptive annotation summaries rather than library-balanced gradient effects. Rows are grouped by the six meta-clusters, with group colors and separators; gene groups follow the corresponding canonical marker panels. All nine nonempty subtype labels and 34,533 current-atlas cells are included. Counts, stored cluster identifiers and subtype-to-meta-cluster assignments are reported in Supplementary Table 3.

#### Supplementary figure 9 - Sensitivity of sinkhorn divergance to main hyperparameter

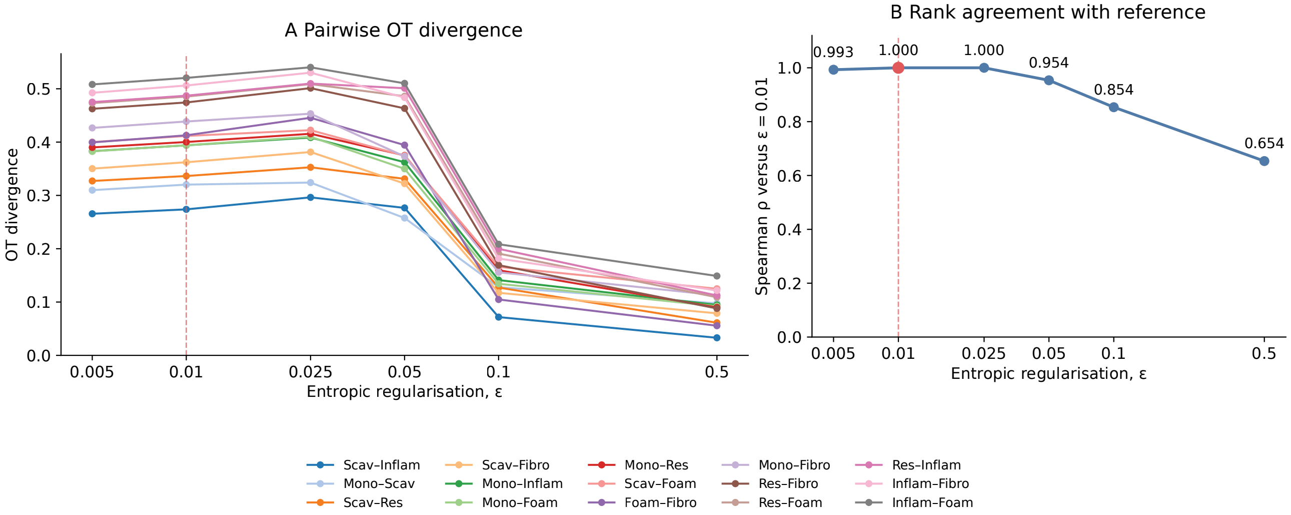


**Supplementary Figure 9 — Sensitivity of OT divergence rankings to entropic regularization.** All 15 meta-cluster pairs were evaluated at $\varepsilon = 0.005$, 0.01, 0.025, 0.05, 0.1 and 0.5 using the same representation, seed and reference sampling scheme (up to 5,000 cells per state). **(A)** Reference OT divergences across regularization values. **(B)** Spearman rank agreement with the selected $\varepsilon = 0.01$ result: 0.993, 1.000, 1.000, 0.954, 0.854 and 0.654, respectively. Rank comparisons use full-precision divergences, avoiding changes in run-specific connectivity scaling. Agreement is strongest near the selected value, while larger regularization changes the ordering of several pairs.

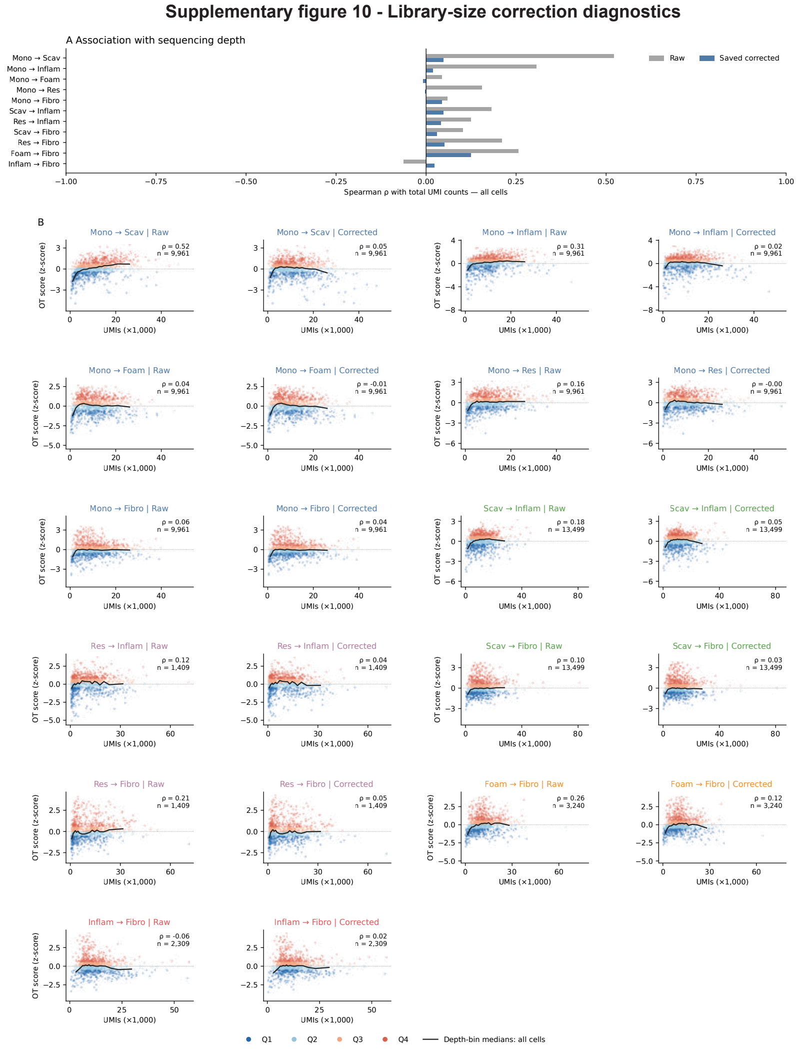


**Supplementary Figure 10 — Sequencing-depth correction of OT target-association scores. (A)** Signed Spearman correlations between total RNA counts and raw or saved OLS-corrected association scores for all 11 comparisons. **(B)** Raw and corrected scores against total counts for each route, using the same deterministically sampled cells in both displays (at most 1,200 plotted cells per route). Colors denote the corresponding saved quartiles. Binned median curves and reported correlations use all matched source cells; rasterization and display subsampling reduce file size without changing calculations. Correction regressed raw scores on total counts with an intercept; no additional rank-based correction was applied. Absolute post-correction Spearman correlations range from 0.002 to 0.124. Pearson correlations are approximately zero by construction. Cell counts are 9,961 for each Mono route, 13,499 for each Scav route, 1,409 for each Res route, 3,240 for Foam–Fibro and 2,309 for Inflam–Fibro. See the Supplementary Note for interpretation.

**Supplementary figure 11 - Broad programme changes across OT-associated routes**

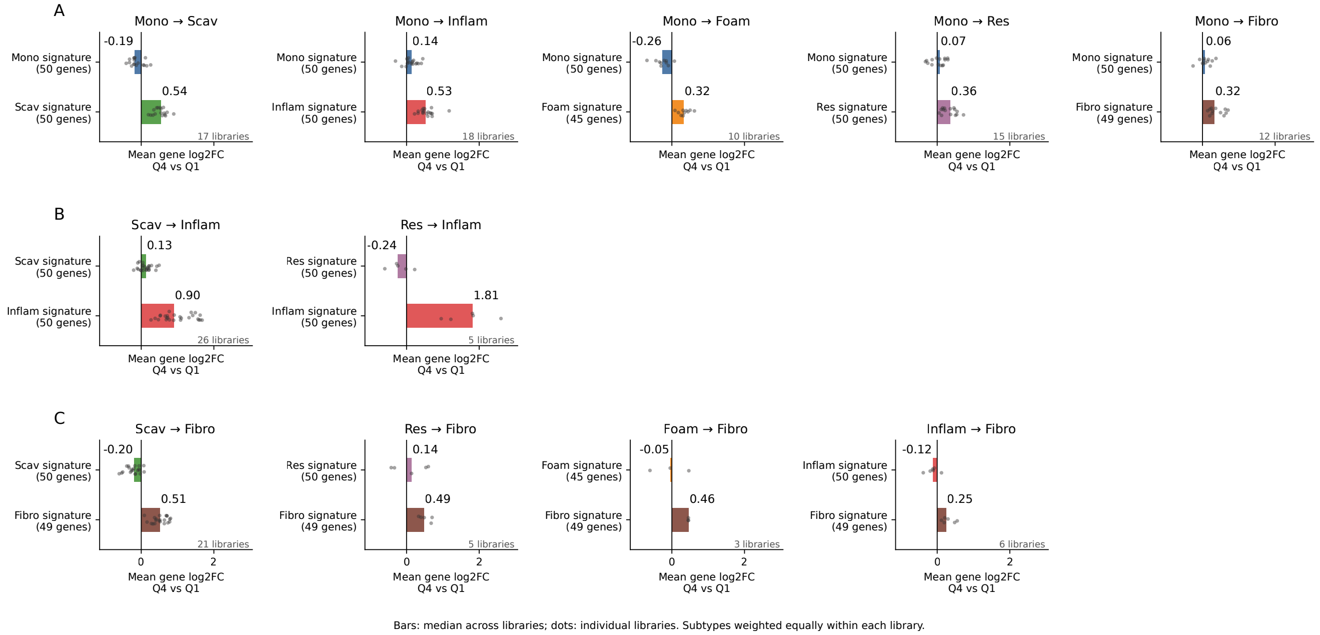


**Supplementary Figure 11 — Broader source and destination signature changes along OT association gradients.** Bars summarize signature-level Q4/Q1 changes as median library effects; small points show individual libraries. Signatures contain 50 genes for Mono, Scav, Res and Inflam, 45 for Foam and 49 for Fibro, selected by the exclusive-signature procedure in the Supplementary Methods. Within each eligible library–subtype stratum, $\log_2$ fold changes were calculated from linearized normalized expression with pseudocount 0.1 and averaged across signature genes. Strata require at least 20 cells in each endpoint; subtype effects are averaged equally within libraries. The panels group 11 unique comparisons into Monocyte-associated comparisons, additional Inflammatory comparisons, and additional Fibrotic comparisons, with source and destination colors. Mono–Inflam and Mono–Fibro are shown once rather than counted twice. Coverage ranges from three to 26 libraries and is route-specific, unlike the jointly eligible 19-library comparison in main Fig. 5D. These are mean gene-level fold-change summaries, not Scanpy module scores, and the atlas-derived signatures are not an independent gene set. Source signatures can show patterns distinct from individual canonical markers; no universal layering/reconfiguration classification is imposed.

#### Supplementary Note — Sequencing-depth correction of OT target-association scores

##### Rationale and implementation

RNA content can covary with both transcriptional state and sampling depth. Association scores derived from an integrated representation are not automatically independent of total counts. Before assigning quartiles, the raw score $s_i$ was regressed on the source cell's total counts $c_i$ within each route,

$$s_i = \alpha + \beta c_i + e_i,$$

and the residual $e_i$ was retained as the corrected score. The current analyses use these saved residuals and quartiles for every route. No switch to rank-based regression was made.

##### Observed count associations

Before correction, Spearman correlations were largest for Mono–Scav (0.522) and Mono–Inflam (0.306); after correction they were 0.048 and 0.019, respectively. Across all 11 comparisons, corrected absolute Spearman correlations ranged from 0.002 to 0.124, the latter for Foam–Fibro. The full values are shown in Supplementary Figure 10 and its source data.

##### Interpretation

Pearson correlations between fitted OLS residuals and the included count covariate are approximately zero by construction, so they are a consistency check on the saved correction rather than independent evidence of biological validity. Spearman correlations describe remaining monotonic association; reduced correlations do not establish the removal of every technical or biological confound. Library–subtype matching addresses an additional source of composition differences when estimating expression effects. Plot subsampling and rasterization alter display only: correlations and binned summaries use all matched cells.
